# Retinal input influences pace of neurogenesis but not cell-type configuration of the visual forebrain

**DOI:** 10.1101/2021.11.15.468630

**Authors:** Shachar Sherman, Koichi Kawakami, Herwig Baier

## Abstract

The brain is assembled during development by both innate and experience-dependent mechanisms^1–7^, but the relative contribution of these factors is poorly understood. Axons of retinal ganglion cells (RGCs) connect the eye to the brain, forming a bottleneck for the transmission of visual information to central visual areas. RGCs secrete molecules from their axons that control proliferation, differentiation and migration of downstream components^7–9^. Spontaneously generated waves of retinal activity, but also intense visual stimulation, can entrain responses of RGCs^10^ and central neurons^11–16^. Here we asked how the cellular composition of central targets is altered in a vertebrate brain that is depleted of retinal input throughout development. For this, we first established a molecular catalog^17^ and gene expression atlas^18^ of neuronal subpopulations in the retinorecipient areas of larval zebrafish. We then searched for changes in *lakritz* (*atoh7*^*-*^) mutants, in which RGCs do not form^19^. Although individual forebrain-expressed genes are dysregulated in *lakritz* mutants, the complete set of 77 putative neuronal cell types in thalamus, pretectum and tectum are present. While neurogenesis and differentiation trajectories are overall unaltered, a greater proportion of cells remain in an uncommitted progenitor stage in the mutant. Optogenetic stimulation of a pretectal area^20,21^ evokes a visual behavior in blind mutants indistinguishable from wildtype. Our analysis shows that, in this vertebrate visual system, neurons are produced more slowly, but specified and wired up in a proper configuration in the absence of any retinal signals.

## Introduction

The assembly of neuronal circuits is classically considered an activity-dependent process^4^. Indeed, an abundance of evidence shows that central brain areas require input from the sensory surface for specification or maintenance of synaptic connections^1,3,5,12,13,22^. In the field of computer vision, machine learning algorithms employ an extreme version of this principle, namely that the architecture of initially randomly wired neuronal networks is entrained entirely by the structure of image data. However, the extent of malleability of biological brains by sensory inputs has not been explored systematically. Modern single-cell RNA sequencing techniques^17^ offer the opportunity to test how the sensory surface influences gene expression and cell-type composition of the brain. Two recent studies of the mouse visual cortex, however, have arrived at divergent conclusions^23,24^, emphasizing the need for more systematic work.

In the visual system, retinal ganglion cells (RGCs) break down the scenery into features and relay those to a range of anatomically well-defined central areas^25–28^. Visual motion is the best studied example of how central circuits are shaped by experience. Experimental manipulation of direction- or orientation-selective RGC inputs disrupts, or otherwise alters, the emergence of properly tuned neurons in cats^2^, ferrets^37^, and Xenopus tadpoles^11^. Even before eye opening, waves of spontaneous activity that sweep across the mammalian retina simulate sequential activation of neighboring RGCs by directional movement^16^; these activity patterns facilitate the formation of visual maps and the segregation of eye-specific inputs in binocular areas^38,39^. In Xenopus tadpoles, experimentally inverting the direction of optic flow prevents the refinement of the retinotectal map^40^. Perhaps most strikingly, directional tuning of mouse RGCs can be reversed by repeatedly showing a stimulus moving in the non-preferred direction^10^. Given that the development of RGCs with different preferred directions is under tight genetic control^41^, these findings imply that sensory activity is able to re-program cellular fates. In addition to imposing patterns of activity on their postsynaptic targets, RGCs secrete mitogens, such as Sonic Hedgehog, and other factors, which could regulate proliferation, commitment, migration and differentiation of downstream neurons and glia^7,8^.

Here we have systematically explored the development of zebrafish visual centers in the absence of RGCs. In larval zebrafish, RGCs project to 12 visual centers in the forebrain and midbrain^29,30^, most of which have mammalian homologs. Moreover, the zebrafish visual brain is rapidly developing, continually growing and plastic^31–35^. Eyes are always open, and visually evoked responses are seen as early as 2.5 days after fertilization^36^, affording sensory experience and spontaneous activity ample opportunity to shape downstream circuitry. Removal of RGCs in the embryo may thus profoundly perturb development of the central brain. We first established a molecular cell-type catalog of the larval thalamus and pretectum, two major divisions of the diencephalon that receive retinal input. We then investigated the cellular composition of these areas in *lakritz* mutants. *The lakritz* mutation disrupts the basic helix-loop-helix transcription factor Atoh7 (Ath5) gene, which is critical for RGC cell fate determination and is not expressed outside of the retina^19^. Homozygous mutants are viable and show normal behavior, except for their blindness. The *lakritz* mutation is completely recessive^19^. Visual experience, spontaneous activity and RGC-derived molecular cues are all eliminated in this mutant. We show that absence of retinal input delays the genesis of central cell types, but, perhaps surprisingly, leaves their final configuration unaffected.

### Generating a molecular atlas of cell types across retinorecipient areas in larval zebrafish

The enhancer-trap line *T*g(*HGn12C:GFP*)^42^ labels most, or all, neurons in pretectum, dorsal thalamus and ventral thalamus, as well as several anterior nuclei of hypothalamus and a subset of neurons in habenula, tectum, nucleus isthmi and medulla. Non-neuronal cells labeled in *Tg(HGn12C:GFP)* include oligodendrocytes and neuronal progenitor cells (Fig. 1a). We transcriptionally profiled 123,224 fluorescently sorted cells with 95,122 passing quality control and clustered glutamatergic and GABAergic neurons independently (Fig. 1b-f). This resulted in 40 glutamatergic and 37 GABAergic clusters, each identifiable by one or more specific marker genes (Fig. 1g,h). The subpopulation-specific expression of markers was verified using a multiplexed wholemount fluorescent in-situ hybridization protocol (Fig. 2; Supplementary Fig. S1). Labeling patterns were registered to a standard reference brain (Fig. 2; Supplementary Fig. S1,S2) and related to classically annotated brain regions in the digital Max Planck Zebrafish Brain Atlas^43^ at https://fishatlas.neuro.mpg.de (Supplementary Fig. S3a-c; Table 1).

**Figure 1.**
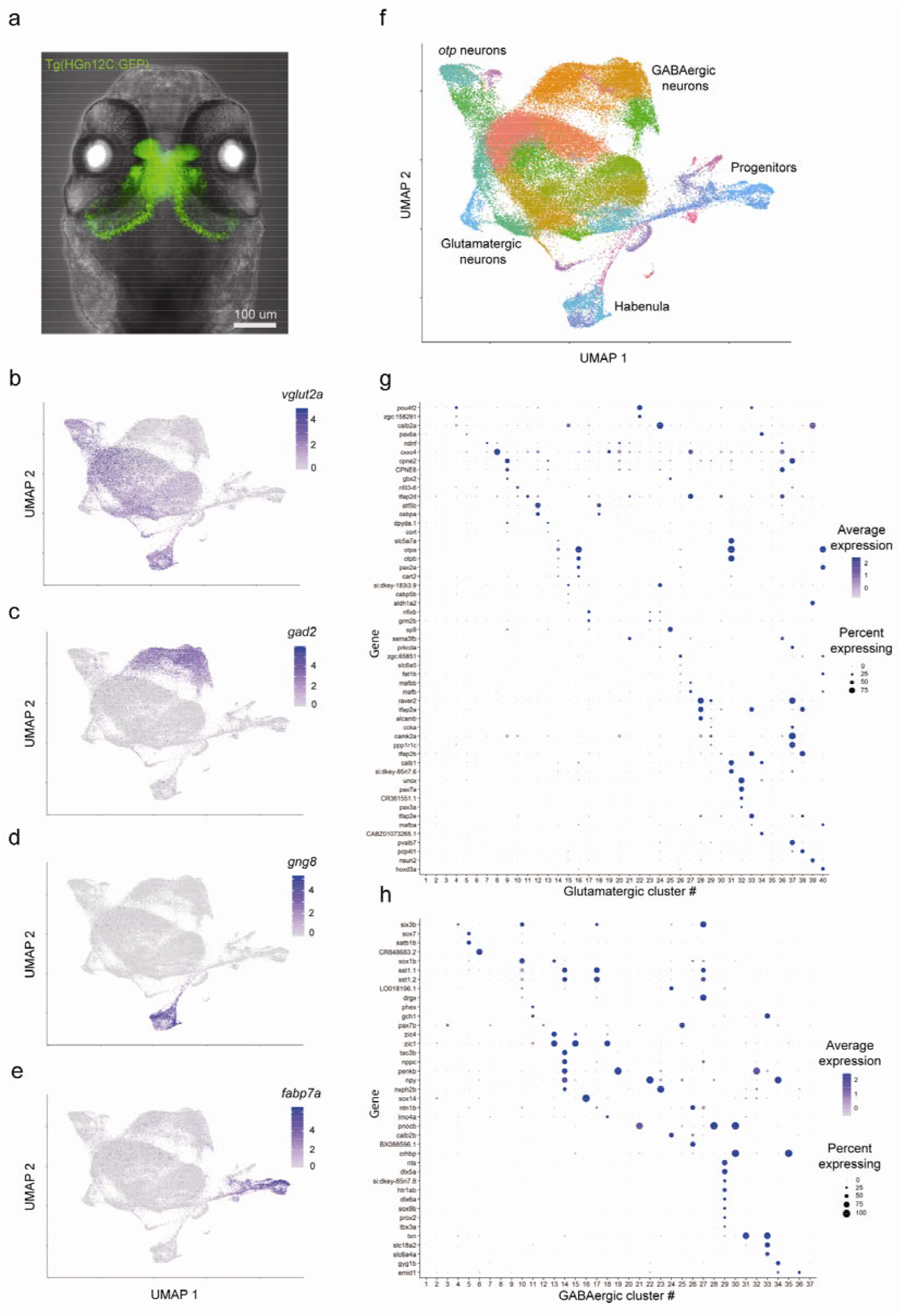
scRNA-seq reveals cell types across visual-processing centers. (a) Maximum Z projection of the *Tg(HGn12C:GFP)* expression pattern in green. In gray, transmitted light. (b-e) Gene expression plots in cells embedded in UMAP space (*vglut2a*, glutamatergic neurons; *gad2*, GABAergic neurons; *gng8*, habenula neurons; *fabp7a*, progenitors). (f) UMAP embedding of all sequenced cells. Color coding represents different clusters. Text labels adjacent cell classes. UMAP space is the same as in (b-e). (g,h) Markers for glutamatergic (top) and GABAergic (bottom) clusters. Color shade represents the average level of marker expression in a cluster (average expression). Dot size represents the percentage of cells expressing the marker in a cluster (percent expressing).

**Figure 2.**
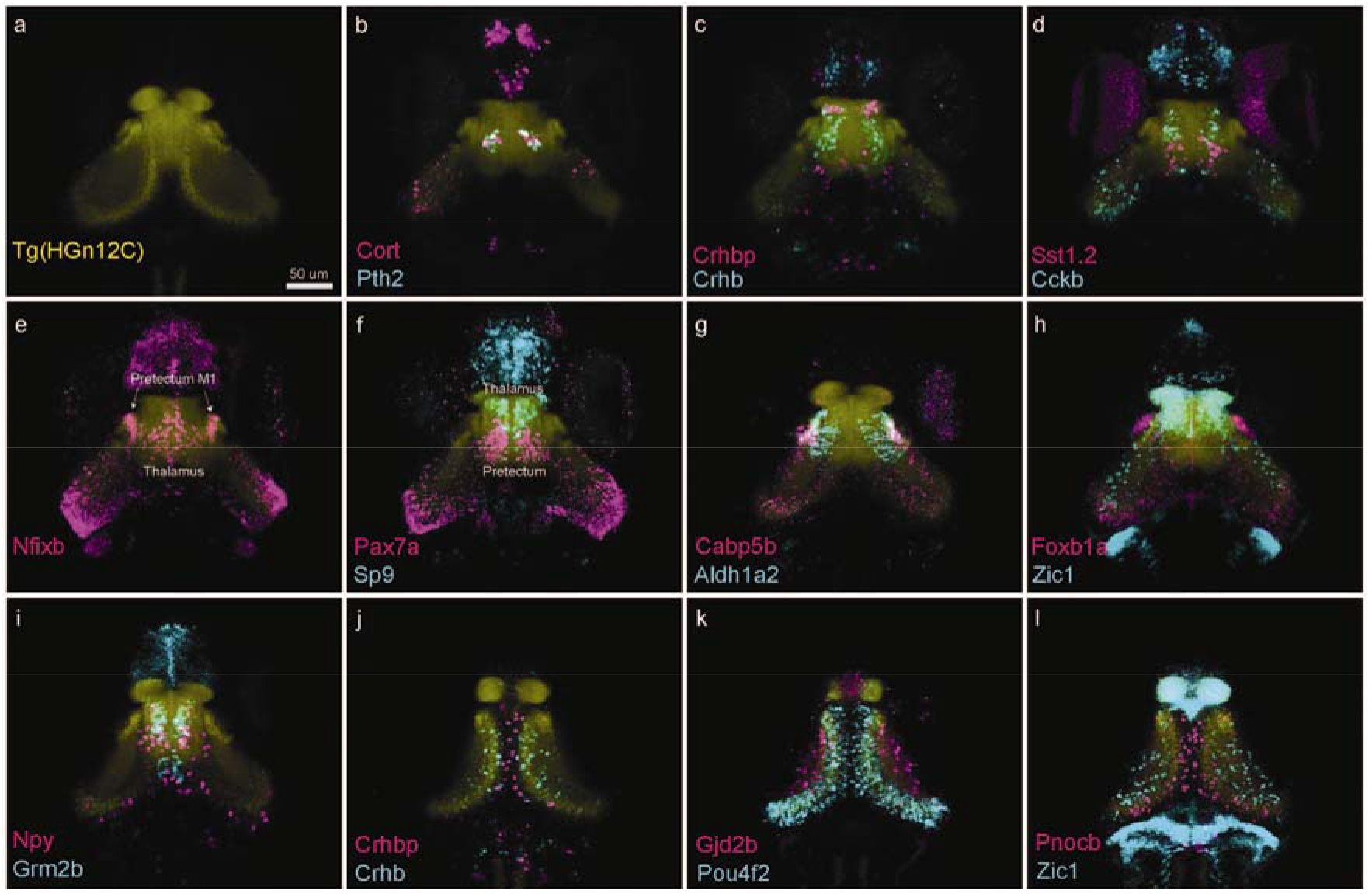
HCR-FISH uncovers the spatio-molecular organization of visual-processing centers. Marker genes validated by HCR in-situ labeling cells in discrete visual-processing areas. (a-i) Substack maximum Z projections of registered HCR-FISH stains. In yellow, Tg(HGn12C:GFP) background stain used for registration. (b-f) selected thalamic markers. From left to right: *cort, pth2, crhbp, crhb, sst1*.*2, cckb, sp9*. (e-i) selected pretectal markers. From left to right: *nfixb, pax7a, cabp5b, aldh1a2, foxb1a, zic1, npy, grm2b*. (j-l) selected tectal markers. From left to right: *crhbp, crhb, gjd2b, pou4f2, pnocb, zic1*.

Our analysis discovered novel type-specific markers and also revealed gene expression domains that had previously escaped detection. For example, it has been reported that the neuropeptide-encoding gene *pth2* is expressed in a small subset of thalamic cells involved in mechanosensation^44^. Our data uncovered that there are two thalamic *pth2+* populations, one also expressing the neuropeptide Cortistatin (encoded by *cort*/*sst7*) and a previously unreported tectal population (Fig. 2a; Supplementary Fig. S2). The gene *pou4f2* encodes a transcription factor specifically labeling glutamatergic neurons of the tectum (Fig. 2k; Supplementary Fig. S2). The gene *gjd2b* encodes a gap-junction protein differentially expressed only in tectal superficial interneurons (SINs; Fig. 2k; Supplementary Fig. S2). The genes *pax7a, aldh1a2*, and *cabp5b* are relatively specific markers for pretectal cells (Fig. 2f,g; Supplementary Fig. S2). Expression of the transcription factor gene *sp9* offers a molecular landmark separating the dorsal thalamus from the pretectum (Fig. 2f; Supplementary Fig. S2). In the thalamus, individual cell types were identifiable by the transcription of *crhb, crhbp, cckb, atf5b, npy, cort* and *sst1*.*2* (Fig. 2b-d; Supplementary Fig. S2). While the adult mouse thalamus has been reported to lack gene expression domains that demarcate separable subdivisions^45^, the larval zebrafish thalamus is organized in spatial clusters likely corresponding to single brain nuclei.

The pretectal migrated area M1 is partially homologous to the mammalian accessory optic system^46^. The genes *calb2a, nfixb, esrrb, sox14* and *zic2a* each label non-overlapping M1 cell populations (Fig. 2e; Supplementary Fig. S2). The gene *pax6a* additionally shows expression in two nuclei bordering on M1 (Supplementary Fig. S2). Lastly, we also identified markers for visual centers in the midbrain, such as *BX088*, an unknown gene, and *sema3fb*, which are both expressed specifically in the nucleus isthmi^47^ (Supplementary Fig. S2, S3). In conclusion, the molecular cell-type catalog and spatial gene expression atlas presented here reveal the complex genetic architectures of the diencephalon and parts of the mesencephalon in larval zebrafish.

### Most, or all, cell types develop autonomously from retinal input

Next we crossed the *Tg(HGn12C:GFP)* reporter into a *lakritz* mutant background and sequenced 20,221 fluorescently sorted mutant cells and 25,687 WT sibling cells, with 17,029 *lakritz* and 18,443 WT sibling cells passing quality control. Strikingly, we could not detect a major difference in the transcriptomic profiles between the different genotypes (Fig. 3a): Every cell cluster that developed in wildtype (WT) is also present in mutants and phenotypically normal siblings. Comparing individual replicates of each genotype excluded a possible confound of batch effects (Supplementary Figs. S4, S5). The similarity of *lakritz* to WT clusters was also robust after bioinformatic separation of glutamatergic and GABAergic neurons (Supplementary Fig. S6-S9), with few exceptions (see Methods). Embedding all samples together in a single UMAP plot did not reveal a difference between samples (Fig. 3b,c). We color-coded *lakritz* cells based on results from *lakritz*-only clustering and were able to see the same clusters forming. This visualization excludes the possibility that a larger WT sample would force the *lakritz* population to embed similarly (Supplementary Fig. S10).

**Figure 3.**
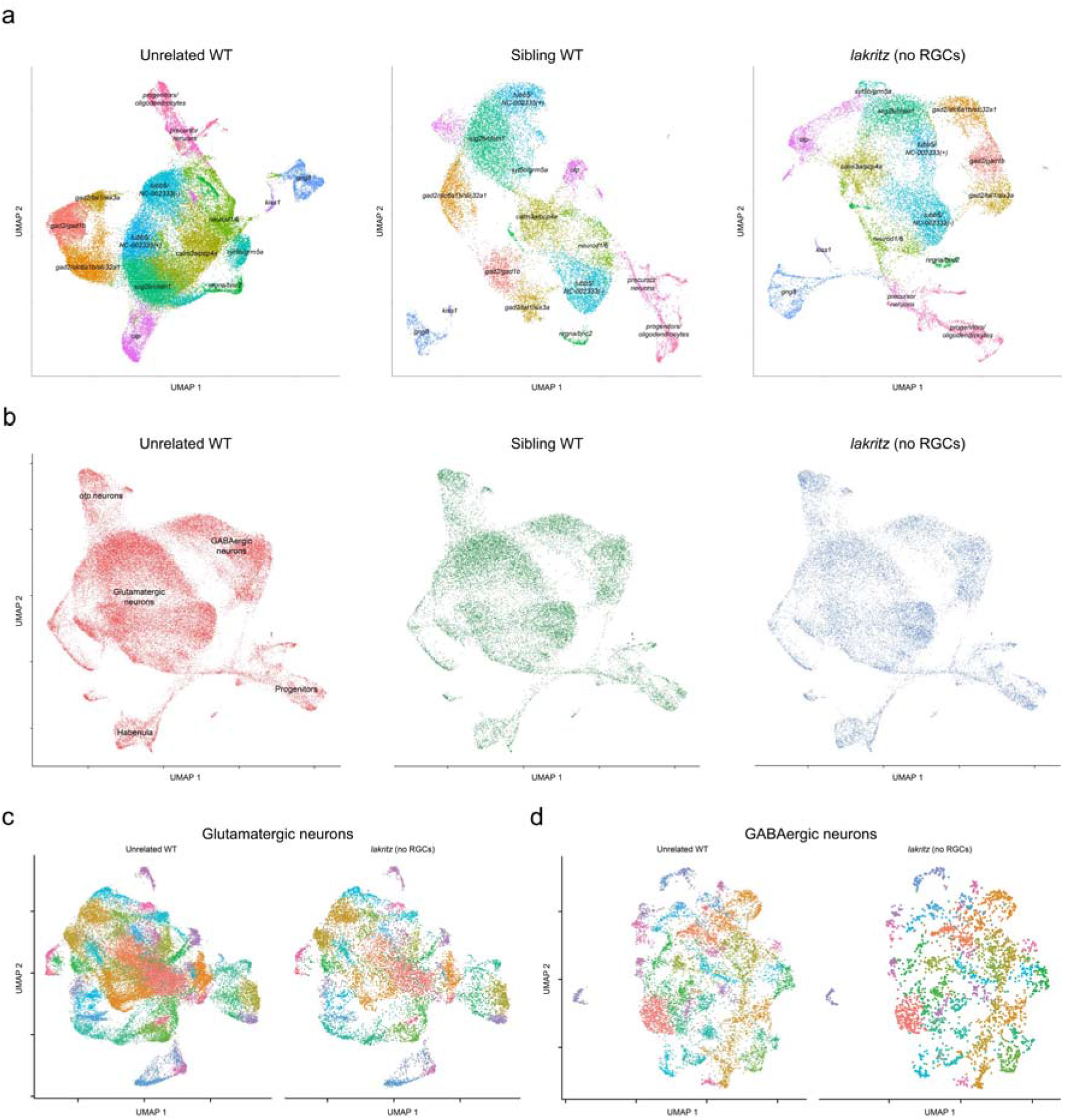
Forebrain cell-type diversity emerges in absence of retinal input. (a) UMAP embedding of different genotypes clustered independently (unrelated WT, left; sibling WT, middle; lakritz, right) color-coded by cluster identity. Clusters are labeled with text. GABAergic clusters express gad2. The rest of the clusters, excluding progenitors and precursor neurons, are glutamatergic clusters. (b) UMAP embedding of all cells color-coded by genetic background. Text labels adjacent cell classes. (c) Clustering of glutamatergic (left) and GABAergic cells (right). Presented side-by-side are WT and lakritz cells of the same clusters. For a quantitative analysis, see supplements.

To explore whether the 2D UMAP dimensionality reduction was sufficiently sensitive to reveal subtle differences, we designed a computational pipeline simulating a cell type ablation experiment. For this, an algorithm omitted *lakritz* cells belonging to a single glutamatergic or GABAergic cluster from the original count matrix. Then, an unbiased pipeline processed the truncated matrix in the standard fashion. We avoided any type of batch correction as that can mask differences. After the pipeline completed, we visualized individual runs and confirmed, or rejected, where we hypothesized an effect was present. In cases where we could not see an effect, we counted the run as negative. Overall, we correctly identified a missing cluster in *lakritz* in 71% of GABAergic clusters and 87% of glutamatergic clusters (Supplementary Fig. S11-S14). Together these clusters contain 90% of all cells we sequenced. In cases of clusters where we could not detect an effect, we noticed that these clusters often embedded poorly in a 2D UMAP; they show a dispersed pattern, apparently mixed among many other clusters, making it difficult to visualize these cells as a single cluster (Supplementary Fig. S11d, S13d). Thus, our “in-silico cluster ablation” approach further validates the conclusion that the vast majority of forebrain cell types, if not all, are present in *lakritz* mutants.

### Individual genes are dysregulated in the forebrain of *lakritz* mutants

Next we asked if we could detect global, or cluster-specific, differences between samples. In the absence of a generally agreed-on statistical test for differences in these kinds of multidimensional datasets, we carried out a straightforward three-way comparison between mutants, heterozygotes, and unrelated WT. We searched for genes that were differentially expressed between the three groups. A large number of genes indeed varied between *lakritz* and WT (Supplementary Fig. S15a). These differences are slightly less pronounced between the two WT groups. However, only a few of the potentially dysregulated genes pass a threshold defining markers (see Methods).

The small variation in gene expression extended to the principal components (PCs) (Supplementary Fig. S15b). While, in some cases, PCs are switched, the top PCs are identical across groups. If the expression of individual genes were systematically different between genotypes then the nearest neighbors of *lakritz* cells in our UMAP embedding should more often be other *lakritz* cells rather than a mix of both. This statistical effect should be enhanced if the differentially expressed genes are also marker genes and would create local ‘hotspots’ in the UMAP space, populated preferentially by either *lakritz* or WT cells. Alternatively, changes in gene expression might be more distributed, with most, or all, clusters affected similarly. The latter scenario applied to our data: Altered neighborhoods exist throughout the *lakritz* dataset (Supplementary Fig. S15c), suggesting that the differences are broadly distributed among the clusters. The magnitude of overall changes between pairs were highly similar, i.e., WT clusters were not more similar to phenotypically normal siblings than to mutants. Any difference between mutant and WT did not translate into a difference between mutant and heterozygous siblings, as would be expected for a correlation with phenotype (the absence of RGCs).

Lastly, we tested whether we could identify differences in the relative proportion of specific clusters or clusters that are particularly sensitive to the absence of RGCs. We could not detect significant differences in the abundance of specific clusters between samples (Supplementary Fig. S16). To test whether some cell types show more severe transcriptome alterations than others by the absence of RGCs, we applied an analysis used before to uncover such clusters^48^ (see Methods). We used p-values calculated in this analysis to color-code our UMAP, highlighting cell types showing the most pronounced transcriptional changes (Supplementary Fig. S17). We then compared the expression patterns of cluster-specific markers in WT and *lakritz* by semi-quantitative HCR in-situ labelings^18^. This analysis revealed local differences in expression of some genes (Supplementary Fig. S18). For example, *aldh1a2* and *cabp5b* show strong expression in cells of the dorsolateral central pretectum and, additionally, the *aldh1a2* probe labels more medial pretectal cells. In *lakritz* mutants, the level of expression in the latter population is diminished, while the former population shows downregulation only in *cabp5b* (Supplementary Fig. S18b,c). The *calb2b* gene is downregulated in *lakritz* mutants across the tectum (Supplementary Fig. S18d). The *crhbp* marker is expressed in multiple nuclei across the forebrain, but noticeably downregulated in a single thalamic nucleus (Supplementary Fig. S18e).

### Differentiation trajectories are unaffected, but cell cycle exits are delayed, in the *lakritz* brain

In addition to neurons and glia, the *HGn12C:GFP* reporter also labels progenitors and differentiating precursor cells. These groups can be clearly distinguished based on their transcriptional profiles (Fig. 1e,f). In our UMAP embedding, progenitors and differentiated neurons are connected by a single “thread-like” cluster of precursors, reflecting the gradual transition of the single-cell transcriptomes from uncommitted, mitotic to postmitotic, fate-committed terminal stage. Further visualization revealed a split of late-stage precursors into a glutamatergic (expressing *neurog1*) and a GABAergic (expressing *ascl1b*) branch in all samples (Fig. 4a; Supplementary Fig. S19a,b). In addition, a subset of the glutamatergic precursors branches off to a habenula fate (expressing *cxcr4b*; Supplementary Fig. S19c). Confirming a previous report^49^, mature habenula neurons fall into one of two subpopulations: *gng8*-positive or *kiss1*-positive cells (Fig. 4a; Supplementary Fig. S19c). Subpopulations of precursor neurons express two kinds of marker genes: “cell-type markers”, which are characteristic of differentiated neuronal clusters, probably indicating commitment to a specific fate, and “transient markers”, which are downregulated in mature neurons and may contribute to the developmental transition (Table 2).

**Figure 4.**
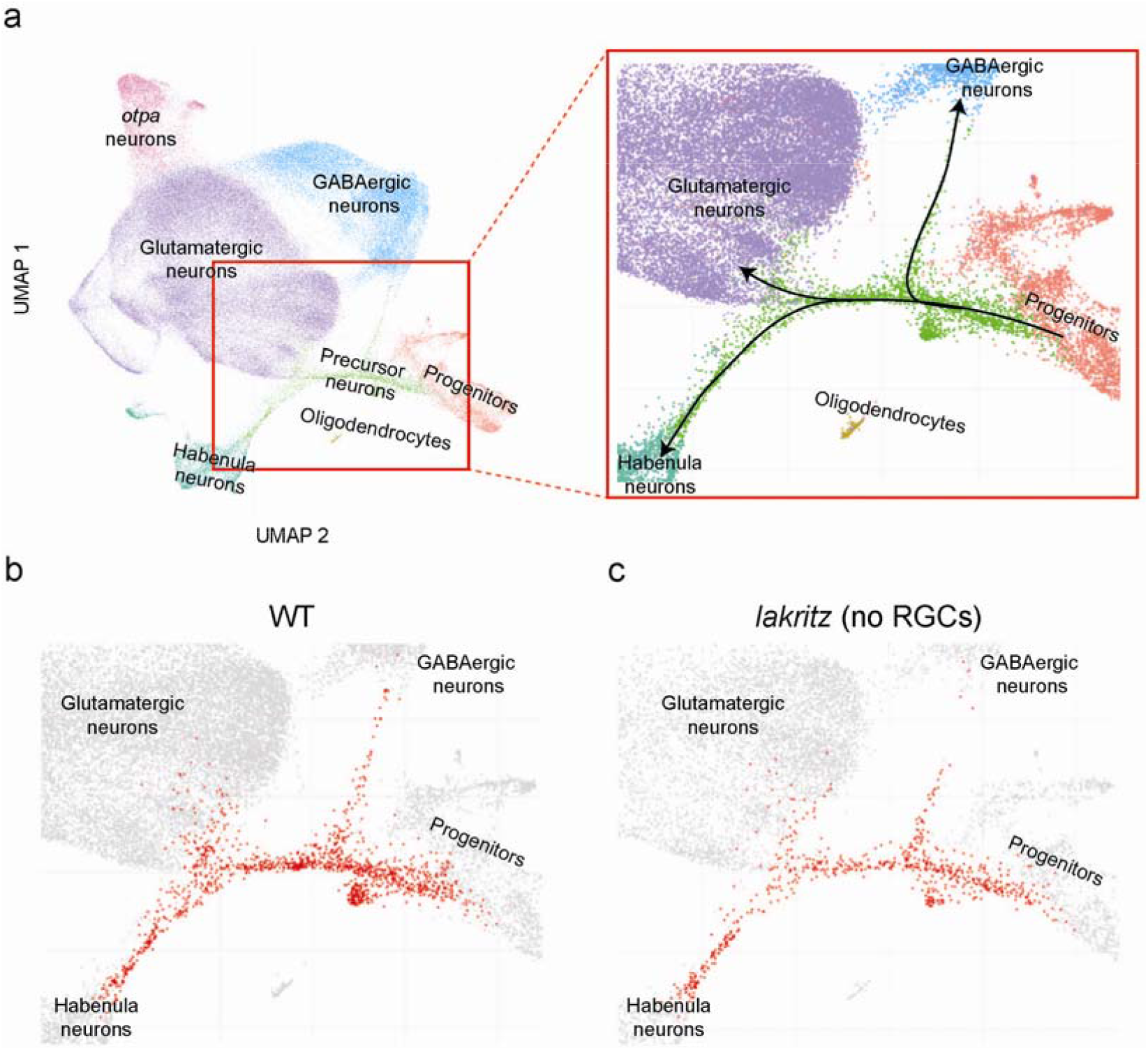
Differentiation trajectories are conserved in absence of RGCs. (a, left) UMAP embedding of all cells color-coded by cell identity. Text labels adjacent cell classes. (right) Enlarged area (red square, left) showing a class of neuronal precursor cells differentiating into the major cell classes. (b,c) same enlarged area as in (a, right), but showing only WT cell (b), or *lakritz* (no RGCs) cells. In red are highlighted all neuronal precursor cells showing splits along the same differentiation trajectories.

This resolution afforded us the opportunity to investigate if the absence of RGCs altered developmental trajectories. In *lakritz* mutants, differentiation pathways are akin to WT (Fig. 4b,c). Strikingly, however, progenitors are enriched relative to neurons in *lakritz* mutants (WT: 82.6% neurons, 4.4% progenitors, ratio = 19; *lakritz*: 73.5% neurons, 7.9% progenitors, ratio = 9; p-value < 2.2#10^16^ Chi-squared test). We used semi-quantitative HCR in-situ stains to explore the location and expression of marker genes expressed in progenitors (*pcna, her4*.*1*, and *fabp7a*) and differentiating precursor neurons (*ascl1b, neurod1* and *neurod2*) (Supplementary Fig. S20). In *lakritz* mutants, the mitotic marker *pcna* was upregulated, while both *fabp7a* and *her4*.*1* were downregulated. Markers of newly committed precursors, such as *neurod1* and *neurod2*, were reduced in *lakritz* mutants. These results indicate that absence of RGCs increases the progenitor-to-precursor ratio, probably by delaying cell-cycle exit.

### Pretectum circuitry forms in the absence of retinal input and supports behavior

We set out to test whether functional visual networks form during development without the input from RGCs. Optogenetics was used to probe synaptic circuitry in *lakritz* mutants. Previous work had identified the direction-selective circuitry in the ventro-anterior pretectum as a hub driving the optokinetic-response (OKR) in zebrafish larvae^20,21,46^. Photostimulation of channelrhodopsin (ChR2)-expressing neurons in the transgenic line *Gal4s1026t* via an optic fiber elicited a sequence of slow pursuit eye movements and saccades typical of the OKR^21^. We crossed *Gal4s1026t; UAS:ChR2-mCherry* into a *lakritz* mutant background and exposed fish larvae sequentially to moving gratings in a visual arena and to targeted photostimulation (Fig. 5a,b). As expected, WT larvae, but not *lakritz* mutants, respond to visual stimulation (Fig. 5c-e). Optogenetic activation of the pretectal area, on the other hand, evokes full OKR-like behavior in both WT and mutants (Fig. 5c-e; Supplementary Fig. S21). Activation of a different population of pretectal projection neurons has recently been shown to trigger prey-capture bouts. Consonant with our finding, this behavior could also be elicited in *lakritz* mutants in the absence of visual stimuli^50^.

**Figure 5.**
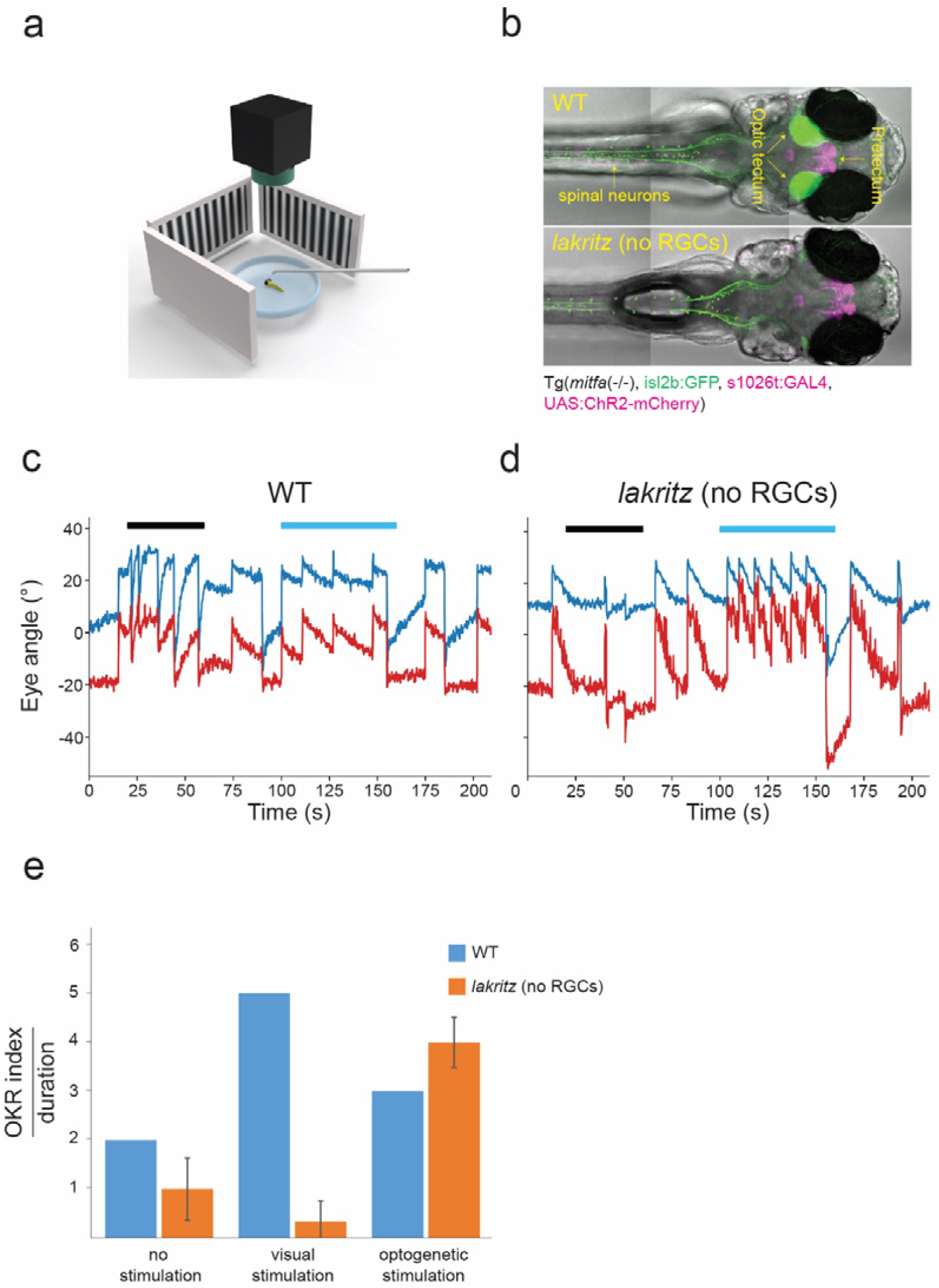
The pretectal OKR circuitry assembles to generate relevant behavior without retinal input. (a) Illustration of experimental setup. Larvae embedded in agarose on a small transparent dish facing screens showing moving gratings (generating optic-flow). A camera records eye movement and an optogenetic fiber can shine light into the brain. (b) Selected images of larvae used for experiment. Maximum Z-projection of either WT (top) or *lakritz* (no RGCs; bottom) 6 dpf transgenic larvae Tg(mitfa (-/-; transmitted light), *isl2b*:GFP (green), Gal4s1026t, UAS:Chr2-mCherry (magenta)). GFP expression was used to phenotype *lakritz* (no RGCs) mutants. (c,d) Eye movement traces of larvae in (b; blue, left eye; red, right eye) either WT (c), or *lakritz* (no RGCs; d). Black line shows interval of visual stimulation (gratings moving). Cyan line shows interval of optogenetic stimulation. (e) OKR index for the three experimental phases (no stimulation, visual stimulation, and optogenetic stimulation). Increase in OKR index indicates a shift from spontaneous eye movement to repetitive synchronized eye movement (for *lakritz* n=3)

To verify that the direction-selective cells driving expression of ChR2 in *Gal4s1026t* overlap with the *HGn12C:GFP* reporter line, we sequenced them and introduced their transcriptomes into our cell-type catalog (Supplementary Fig. S22a). We also co-expressed the two reporters in a triple-transgenic fish (Supplementary Fig. S22b) and co-registered them within the Max Planck Zebrafish Brain Atlas atlas (Supplementary Fig. 22c). Together, these analyses showed that the cells expressing neuronal markers in the *Gal4s1026t* domain were a subset of those present in the *HGn12C:GFP* forebrain catalog. While *Gal4s1026t* labels additional non-neuronal cells (likely glia; Supplementary Fig. S22a), its expression is restricted to a smaller pretectal subvolume than *HGn12C:GFP*.

### Conclusions

This work has explored the development of visual forebrain centers in absence of any retinal signals, be it synaptic activity or secreted molecules. We resolved, for the first time, the diversity of cell types in the thalamus and pretectum. We compared this to the population of cell types that develop in mutants specifically lacking RGCs at all stages of development. Strikingly, we could not detect a single cell type that failed to develop, or one that developed along an altered differentiation pathway. However, without retinal input, progenitors remained on average longer in the cell cycle. Thus, RGC-derived signals facilitate the transition from uncommitted, cycling progenitors to differentiating neurons, without affecting terminal fate selection. Lastly, we and others^50^ show that a forebrain devoid of retinal input assembles into circuits capable of generating visual behavior, ranging from OKR to hunting.

The zebrafish visual system develops rapidly, grows through adulthood and shows substantial plasticity^31^. Nevertheless, the sensory surface appears to play a subordinate role in shaping the composition or function of central targets. In broad agreement with our findings, a recent publication demonstrated that visual experience in mouse is not necessary for development of visual cortex cell types^23^. In contrast, a preprint^23b^ reported that cells do not differentiate properly in a specific layer (L2/3) of the mouse visual cortex, demonstrating that some aspects of mammalian neocortex development rely on signals from the sensory surface. Single-cell transcriptomics, ideally combined with circuit reconstruction and functional reconstitution via optogenetics, is slated to shed light on the relative contributions of experience, spontaneous activity and genetic programs to assembly of the brain.

## Acknowledgements

We would like to thank the Max Planck Campus Martinsried sequencing facility for their assistance, especially Markus Oster for FACS processing and troubleshooting, and Marja Driessen and Rin Ho Kim for next-generation sequencing and initial bioinformatics. We would also like to thank Yvonne Kölsch for initial help in sample preparation. Irene Arnold-Ammer provided superb technical assistance. We thank Gregory Marquart, Manuel Stemmer, Inbal Shainer, Johannes Larsch, Fumi Kubo, Karthik Shekhar, Joshua Hahn and Salwan Butrus for critical feedback and discussions.

## Methods

### Zebrafish husbandry

Adult and larval zebrafish were maintained on a 14:10 hour light:dark cycle at 28°C. Embryos were kept in Danieau’s solution (17 mM NaCl, 2 mM KCl, 0.12 mM MgSO4, 1.8 mM Ca(NO3)2, 1.5 mM HEPES). All animal procedures conformed to the institutional guidelines set by the Max Planck Society, with an animal protocol approved by the regional government (Regierung von Oberbayern).

For single-cell experiments one male and one female were placed in individual breeding tanks with a divider the evening before spawning. Dividers were removed at 9.00 am the next morning and fish let spawn until 10.00 am. Eggs were collected shortly after and mixed together. *Tg(HGn12C:GFP)* larvae were sorted for GFP expression 24 hours after fertilization. All single-cell experiments were performed at 6 days after fertilization. For WT *Tg(HGn12C)* experiment, pigmented larvae were used from *Tg(HGn12C)* pigmented males or females crossed with WT Nacre (*mitfa* -/-). For *lakritz* and sibling WT experiments, *Tg(HGn12C:GFP) lakritz* heterozygote adults were crossed with adult *lakritz* heterozygotes. For experimetns with *lakritz* larvae, only *lakritz* were collected, for other experiments they were discarded. either. For *Tg(1026t)* experiment, *Tg(Gal4s1026t)* were crossed with *Tg(UAS:GFP)*. All larvae were reared at a maximum density of 60 individuals in 10 cm petri dish.

### Cell-dissociation for single-cell RNA sequencing

Six-day-old larvae expressing GFP under *Tg(HGn12C)* or *Tg(Gal4s1026t)* were used to label neuronal cell types in the forebrain. Ames medium (Sigma A-1420) was used throughout and prepared according to the manufacturer’s guidelines. Before animal handling, 300 ml Ames medium was oxygenated in room temperature for one hour. Buffer pH was adjusted to 7.2-7.3 using HCl and filtered through a standard Steritop filter. Buffer pH was checked again after filtration for a desired range of 7.4-7.5. Oxygenated Ames medium was then placed on ice. Larvae were anesthetized in oxygenated Ames ice slush and rapidly decapitated. Brain material from a maximum of 110 larvae was collected using a broad glass pipette and transferred into chilled oxygenated Ames in a 2 ml tube on ice. Tube medium was replaced with fresh Ames after every material transfer. In addition to transgenic larvae expressing fluorescent protein, 30 non-transgenic larvae were used to adjust FACS gates. Cell suspensions from both samples were prepared in-parallel.

Tissue was dissociated into single-cells using a modified protocol from (Kölsch et al. Neuron 2021). Papain solution [25 U/ml final] was prepared by mixing 4810 µl of oxygenated Ames with 89.3 µl papain stock 42.8[mgP/ml], 32.7[U/mgP], 50 µl DNaseI [13K U/ml] (Sigma D-4527, 40K Units), and 50 µl L-cysteine [152.2 mM] (Sigma C-1276). Papain solution was then placed for 15 minutes in a tabletop spinning incubator preheated to 34°C spiining at 10 RPM. The solution was then examined: though initially milky, the papain solution becomes transparent when activated. Papain solution was not filtered pre- or post-activation. Ames buffer was removed from sample tubes and replaced with 2 ml activated papain solution. Samples were placed in the same tabletop incubator at 34°C for one hour spinning at 10 RPM. After 20 minutes, samples were carefully triturated five times with a narrow glass pipette flamed at the tip to avoid sheering. After one hour, the sample was placed shortly on a bench to let the biological material sink to the tube’s bottom. Papain solution was removed and replaced with 1 ml papain inhibitor solution (4450 µl oxygenated Ames, 500 µl ovomucoid stock, 50 µl DNaseI [13K U/ml]). 10x ovomucoid stock was prepared as follows: 150 mg BSA (Sigma A-9418), 150 mg ovomucoid (Worthington LS003087), 10 ml Ames buffer; pH adjusted to 7.4, filtered and then stored at -20°C. After resuspension in inhibitor solution, tissue cells were completely dissociated by triturating a maximum of 30 times with a p1000 pipette (not broad-end) set to 850 µl. A good indicator of a successful dissociation was that there were no observable white particles (brain matter) in solution. Intact eyes were a good indicator that the dissociation was sufficiently gentle to allow high cell-survival. After mechanical dissociation, 1 ml of inhibitor solution was added to each tube. Samples were passed through a pre-wet 30 um filter (Sysmex). Wetting the membrane with Ames buffer allows liquid and small particles to smoothly pass through the mesh filter. Samples were pelleted in a centrifuge pre-cooled to 4°C for 10 minutes at 300 g. Supernatant was removed and the pellet resuspended in 2 ml Ames with non-acetylated BSA (4.5 ml oxygenated Ames, 500 µl 4% non-acetylated BSA, 0.5 µl DNaseI [13K U/ml]). The resuspended solution was filtered through a pre-wet 20 um filter into a new 2 ml tube and short spun to get all liquid through the filter. 2 µl of calcein blue [1 µl/ml] was added to the solution to stain live-cells. Calcein blue was not added to control samples. Suspensions were kept on ice and processed further by FACS.

### FACS

BD FACSAria III was used to sort cells. FACS gates were set after 500,000 recorded events to sort live single-cells expressing GFP. Similar gates were used across experiments. Cells were sorted using a 100 um nozzle (∼20 PSI) into 2 ml protein LoBind Eppendorf tubes (Eppendorf 0030108132) placed in a cooling holder. Tubes were treated pre-FACS for one hour with 2% BSA in Ames while spinning. Before FACS all liquid was removed. The combination of treatment with LoBind plastic ensured cells will not adhere to the collection tube after FACS. Collection tube was filled before cell-sorting began with 500 µl FACS collection medium containing: 750 µl oxygenated Ames, 250 µl non-acetylated BSA (stock 4%), 0.1 µl DNaseI [13K U/ml]. In total, we aimed to collect 200,000 cells, as in our hands this would fill a 2 ml tube completely. After FACS completed, cell-suspension was centrifuged at 4°C for 5 minutes at 300 g. Centrifuge was not pre-cooled. We found it helpful to memorize the tube’s orientation, as often it was difficult to visualize the pellet. After centrifugation, the pellet was resuspended in 60 µl of Ames with non-acetylated BSA diluted 1:10 in oxygenated Ames (0.04% non-acetylated BSA, final). The medium was slowly let slide over the pellet multiple times, and then the tube was gently tilted back-and-forth to push the suspension over the location of the pellet a few more times. The cell suspension was placed over ice until used for single-cell sequencing.

### Calculating cell-suspension density

13 µl oxygenated Ames was supplemented with 2 µl trypan blue (stain for dead cells) and 5 µl cell suspension (1:4 suspension dilution). 20 µl were loaded into a Fuchs-Rosenthal chamber (NanoEnTek, DHC-F01). A minimum of four large squares were counted for live and dead cells using either a dark-field or DIC microscope. Cell density was routinely between 500-1000 cells/µl with viability at ∼90% (live cells / live+dead cells).

### Single-cell RNA sequencing

Droplet RNA sequencing experiments using the 10X chromium platform were performed according to the manufacturer’s instructions with no modifications. Single-chromium chip channels were loaded routinely with 8000 cells aiming to capture 5000 cells with a doublet rate <5%. In our hands, we noticed that loading 8000 cells would usually result in capturing of 3000-3500 cells. For experiments with WT *Tg(HGn12C)*, 17 replicates were collected across 4 experiments. For *lakritz Tg(HGn12C)*, 10 replicates were collected across 4 experiments. For WT *lakritz* siblings *Tg(HGn12C)*, 8 replicates were collected across 3 experiments. For *Tg(Gal4s1026t)*, 4 replicates were collected from 1 experiments. The cDNA libraries were sequenced on an Illumina HiSeq 2500 to a depth of ∼50,000 reads per cell.

### Alignment of gene expression reads and initial cell filtering

Initial preprocessing was performed using the cellranger software suite (version 3.1.0, 10X Genomics) following standard publisher guidelines. Reads for each channel were aligned to the zebrafish reference genome (GRCz11.98). Further analysis was performed as described below using the Seurat R package (Satija et al., 2015) on the filtered cellranger output matrices.

### Initial analysis using Seurat

Output from cellranger was loaded into Seurat allowing for 200-4000 genes/cell, 400-8000 UMIs/cell, and a maximum detection of mitochondrial genes of 12% of all transcripts detected in cell. Unless otherwise stated, batch correction was performed using harmony on experiment and genotype. For analysis of marker genes, the data was first clustered and separated into glutamatergic and GABAergic neuronal datasets, and then batch-corrected and reclustered. For each dataset we recalculated the 2000 most variable genes (“vst”). We used 18 PCs to cluster our glutamatergic dataset and 20 to cluster our GABAergic dataset. We used a clustering resolution of 1.6 for our glutamatergic dataset and 1.8 for our GABAergic dataset after determining the best resolution using clustree (Zappia et al. GigaScience 2018). Marker genes were extracted using the command FindAllMarkers(…, only.pos = TRUE, logfc.threshold = 0.75), filtered for adjusted p-value < 0.05, and inspected individually using the FeaturePlot visualization tool.

Integration of all three genotypes followed the same pipeline, except that in addition to batch correcting using harmony, we also tested batch correction using the Seurat anchoring method. In short, each combination of experiment and genotype was SCTransformed independently. For each transformed dataset, the 3000 most variable genes were selected as possible integration features. We followed the standard integration pipeline using SelectIntegrationFeatures, PrepSCTIntegration. We used the unrelated WT dataset as our integration space, as it was our largest dataset. We integrated our datasets finally using FindIntegrationAnchors, and IntegrateData. Using this pipeline, we came to the same conclusion as when using Harmony – that we cannot see a difference in cell types between samples.

### Independent clustering of datasets from different genotypes

Data from three genotypes was loaded into Seurat and independently processed and clustered. Separation of the GABAergic and glutamatergic datasets was done as described before, but independently for each genotype. Batch-correction was performed for each dataset using Harmony on experimental replicates. For clustering of all cells, 14 PCs were used for all datasets with a resolution of 1. For the glutamatergic dataset, 18 PCs were used with a resolution of 2. For the GABAergic dataset, 22 PCs were used with a resolution of 2.5.

Matching of clusters between datasets was performed based on unrelated WT marker expression identified using the Seurat command: FindAllMarkers(…, only.pos = TRUE, min.pct = 0.25, logfc.threshold = 0.75). For each cluster, we visualized the best marker genes on the UMAP. Upon confirmation that they were descriptive of the cluster (or additional clusters in the case of over clustering), we visualized the expression in the other datasets and assigned all clusters a name based on the expression of a single marker or multiple markers. In the case where no markers could be verified for a cluster, we looked for the nearby clusters on the UMAP, and merged the cluster with another cluster that showed the fewest distinguishing markers.

In some cases, there were clusters we could not assign to all datasets. For the glutamatergic datasets these were: *tubb5/chd4a, nhlh2/zic2a, pvalb6, pax6a, dlx5a, bhlhe23/grm2b, cabp5b, foxb1a/tfap2e*. For the GABAergic datasets these were: *aldocab, BX088, adarb2, CR34551, emx2, gata2a/nxph1, gyg1b, irx1b, crhbp, otp, onecut1, penkb, pnocb, txn/nr4a2a, uncx4*.*1*. These differences were often the result of the larger cell numbers in the unrelated WT dataset, allowing for the clustering algorithm to more accurately separate cell types. In the cases where a cell type is a rare one, the cells were often merged with larger nearby clusters. Some notable exceptions: *adarb2* cells clustered strongly in lakritz and its siblings, but not in the larger unrelated WT dataset. *Otp*, neurons which are mainly glutamatergic appeared in the unrelated WT GABAergic dataset. *Uncx4*.*1* formed a significant cluster in the unrelated WT dataset, but was largely missing from the *lakritz* and sibling datasets, as was the same for the glutamatergic *dlx5a* cluster. However, there was never a case where cells expressing these markers were completely missing from all other datasets. For closer inspection, see Supplementary Fig. S6, S8.

### Determination of confounding batch-effects

WT and *lakritz* datasets were first analyzed independently. To determine whether there are batch effects within each dataset across experiments, each dataset was first analyzed independently. Normalization, variable feature selection, scaling, and principal component analysis were performed with either WT or *lakritz* cells. The DimPlot visualization tools (Seurat) was used to inspect different combinations of the first 5 PCs where cells were color-coded by experiment. For the WT dataset, where we could detect a batch effect, we tested whether batch-correction using Harmony can correct the effect. We did not test this for the *lakritz* dataset, as we could not detect such a batch effect. To test whether WT and *lakritz* datasets were integrated properly also in absence of batch-correction, we followed standard procedure, but omitted a batch-correction step.

To test whether *lakritz* cluster structure is preserved following integration with WT cells, we first clustered the *lakritz* data independently and saved the results as a large matrix containing cell and “*lakritz*-only” cluster identities. We then followed standard procedure to cluster WT and *lakritz* cells together. Lastly, we plotted *lakritz* cells according to their new UMAP coordinates (embedded with WT) and color-coded the cells according to their “*lakritz*-only” clusters.

### Analysis of globally differentially expressed genes between genotypes

For analysis of genes differentially expressed globally between genotypes, datasets were batch-corrected using Seurat’s anchoring pipeline as described before. We used the FindMarkers command with default parameters comparing either unrelated WT with *lakritz* or unrelated WT with sibling WT. Globally differentially expressed genes were filtered using adjusted p-values < 0.05. We highlighted genes for which the log fold change was either larger than 1 or lower than -1, or where the ratio of expressing cells across groups was either higher than 2 or lower than 0.5. We also ran this analysis using Harmony for batch-correction, but this led to no differentially expressed genes.

### Alignment of PCs between genotypes

Datasets were batch-corrected using Seurat’s anchoring pipeline as described before. PCs were calculated using the Seurat RunPCA function for the first 15 PCs. For each comparison, only genes that appeared across PCs in both groups were compared. PCs were compared by calculating the cosine similarity between all PCs. PCs were compared be calculating the cosine similarity and extracting the highest value for each PC generating a distribution of values for most similar PCs. We generated our null distribution, by randomly dividing our WT dataset into two groups similar in ratio to *lakritz* / unrelated WT. We repeated this ten times to generate a null distribution for PC similarity across different datasets. We used a Wilcoxon signed-rank test to compare the different distribution.

### Comparing neighborhood embedding between genotypes

For each cell, the neighborhood was determined by extracting the 19 nearest neighbors’ genotype composition and calculating the cosine distance between the observed neighborhood genotype composition (unrelated WT / sibling WT / *lakritz*) and the expected neighborhood genotype composition. The null distribution was generated by randomly shuffling the genotype labels and recalculating the distance between observed and expected neighborhood genotype composition. From this, we extracted the standard deviation and calculated a Z-score for each cell in the original dataset. We color-coded our UMAP based on the Z-score to uncover areas in the UMAP (potentially clusters) that show significantly altered neighborhoods in *lakritz*.

### In-silico cell type ablation analysis

Cell identity and cluster information were imported from a completed clustering analysis. Iteratively, *lakritz* cells belonging to a single cluster were removed from the count matrix. After, removal of the cells, the matrix was processed following the standard Seurat pipeline without batch-correction. To determine whether we could observe an area missing in *lakritz* cells, we visualized all three genotypes together in UMAP space. Upon determination of a missing area, we opened an image where cells in the cluster missing *lakritz* cells were highlighted. This either confirmed our suspicion (if the cell type were missing, we could detect the effect), or denied it. We applied this analysis separately to datasets containing glutamatergic or GABAergic neurons for all clusters.

### Finding clusters with altered transcriptomes in absence of RGCs

The analysis was applied as reported in (Sharma et al. Nature 2020) on datasets batch-corrected using Seurat’s anchoring points pipeline as reported before. In short, for each cluster we measured the ratio between WT and *lakritz* for the set of marker genes and for all genes detected in the cluster. We then compared these two distributions using a Wilcoxon signed-rank test and corrected the p-values for multiple testing (depending on the number of clusters) using Bonferroni’s correction. We extracted the WT identities belonging to the cluster and projected them into the formerly analyzed WT dataset in which we performed our marker analysis. We color-coded the cells based on their p-value for transcriptome alteration. From this, we extracted a list of markers expressed in clusters predicted to be the most altered in *lakritz* compared to WT. We performed this analysis also by applying the same processing pipeline starting with only glutamatergic or GABAergic cells, essentially applying normalization within the separated dataset. We also tested this pipeline with Harmony batch-correctio.

### Comparing cell type proportions

For each experiment, we calculated the relative proportion of cells in each cluster as a fraction of the total number of cells captured across replicates in a single experiment. We compared the distribution of proportion of each cluster between WT and *lakritz* using a Wilcoxon signed-rank test and corrected the p-values for multiple testing using Bonferroni’s correction based on the number of clusters.

### Developmental trajectory analysis

The identities of cells belonging to the neural precursor cluster (expressing *ngn1* and *ascl1b*) were extracted and highlighted. Marker genes for this cluster were calculated using the FindMarkers Seurat function. Each gene was visually inspected using the FeaturePlot visualization tool to determine whether it is expressed only in the cluster (transient expression), or also in other clusters (continuous expression). For each gene we also summarized whether it is expressed in other cell classes (GABAergic neurons, glutamatergic neurons, habenula neurons, and progenitors).

### HCR fluorescent in-situ imaging

HCR stains were performed according to the manufacturer’s instructions (Molecular Instruments) with no modifications. Larvae used for HCR stains were reared in PTU from 24 hours to 6 days. At 6 days, larvae were fixed following instructions. All larvae were Tg(HGn12C:GFP) and stained for a maximum of two transcripts labeled with either/both Alexa546 and Alexa647. All probes were purchased from the manufacturer. Imaging was performed on a commercial Zeiss confocal (LSM780) For WT and *lakritz* comparative stains, larvae were generated from parental *Tg(HGn12C:GFP*; *isl2b:RFP, lakritz*+/-) crossed with Tg(*lakritz*+/-). *Lakritz* (no RGCs) phenotype was selected for, or against, using *isl2b:RFP* expression. Comparative stains were performed only on siblings from the same clutch stained in-parallel for each gene. Imaging in *lakritz* and WT was performed using the same imaging parameters in both samples when comparing the expression of the same gene. Images were collected from a minimum of two replicates for each condition. For each individual either one or two tiles were imaged.

### Morphological registration using ANTs

Brain registration using ANTs was performed as described before (Kunst et al. Neuron 2019). In short, a standard brain was generated using 12 HCR stained Tg(HGn12C:GFP) larvae using ANTs. All HCR stains were registered to the Tg(HGn12C:GFP) standard using Tg(HGn12C:GFP) background stain present in all HCR stacks. In case of drift during image acquisition, the transmitted-light channel was used to correct drift using the MultiStackReg ImageJ plugin. Once the Affine transformation was saved, it was applied to all other channels.

For comparative WT and *lakritz* stains, individual confocal stacks were registered first to the Tg(HGn12C:GFP) standard and then differences were inspected by eye. In the case where differences were detected, the original stacks were then inspected to validate differences.

Anatomical masks from the MapZeBrain atlas were registered to the *Tg(HGn12C:GFP)* standard using a bridging *Tg(HuC:H2b-RFP)* from the atlas. *Tg(HuC:H2b-RFP, HGn12C:GFP)* larvae were independently registered to the *Tg(HGn12C:GFP)* standard to generate an H2b-RFP stain in the standard morphological space. Transgenic fluorescence was successfully preserved in *Tg(HuC:H2b-RFP)* by modifying the HCR fixation protocol as reported in (Lovett-Barron et al., Nature Neuroscience 2020).

### HCR image analysis

Individual confocal stacks of HCR stains were aligned to a standard *Tg(HGn12C:GFP)* expression pattern as described before. Color-coded depth projections were generated using an ImageJ script written by Kota Miura. For analysis of HCR signal, individual confocal stacks were binarized using the ImageJ “RenyiEntropy” algorithm applied to individual slices using each slice’s histogram. In slices where the algorithm produced sudden spikes in the total number of pixels compared to surrounding slices, pixels were instead binarized using the “maxEntropy” algorithm. This step improved the binarization process where the “RenyiEntropy” algorithm was sensitive to sudden changes in noise. However, both algorithms were sensitive to different kinds of noise which allowed to generate a combined but smooth binarization mask. The binarized pixels were then multiplied by the image intensity in each pixel. The morphological signal analysis was performed on the product of these images. Registered anatomical masks were generated as described before. For total expression analysis, the sum of the intensity across the binarized pixels was calculated for each anatomical mask. This analysis was performed for replicates of the same stain and averaged across replicates. The final heat matrix was calculated by normalizing the expression of each gene to the highest value for the specific gene. For background normalization, the background was defined as the signal intensity in pixels inside the anatomical mask that are not HCR positive binarized pixels. For predictions of best matches between brain areas and gene expression, the background-normalized signal values were normalized in the heatmap to the highest value in each brain region.

### Optogenetic stimulation and behavioral tracking

Optogenetic stimulation and behavioral tracking were performed as reported in (Wu et al. Neuron 2020). In short, after embedding and removal of agarose close to the eyes, an optic fiber was placed on top of the fish targeting the pretectum. In order to track the eyes while in dark, larvae were illuminated from the bottom using an 850 nm infrared LED. Light emitted by optogenetic illumination was filtered out by an IR filter in fromt of the recording camera (Thorlabs absorptive filter, ND = 1.0). For focal optogenetic activation with ChR2, a 50um optic fiber (AFS50/125Y, Thorlabs) was used to shine blue light (473 nm, 20-40 mW/mm2, Omicron Lighthub) onto the right or left pretectum. In each experiment, larvae were presented with a phase of stationary gratings, followed by moving gratings (40s), followed by stationary gratings, followed by blue light illumination (60s) and stationary gratings. For optogenetic activation of the *lakritz* pretectum Tg(*mitfa* (-/-), *isl2b*:GFP, Gal4s1026t, UAS:Chr2-mCherry lakritz (+/+)) larvae were used. For control, the same larvae lacking ChR2-mCherry expression were used. Larvae were identified as *lakritz* via *isl2b*:GFP expression.

For eye tracking, the angle of each eye was calculated relative to the body midline. During visual stimulation (gratings moving), the eyes of a fish almost exclusively saccade in a single direction intermittent with a smooth movement in direction of motion. When there is no visual stimulation (stationary gratings), the eyes of a fish will saccade in one direction and then in the opposite direction. Hence, saccades are a reliable readout of the optokinetic response (OKR). To calculate an OKR index the saccades in one direction were subtracted from the saccades in the other direction for each eye – producing higher values during OKR. The average of both eyes was taken as the OKR index.

### Validation of *lakritz* genotype

For larvae, *lakritz* mutants were identified by extreme pigmentation. When nacre (*mitfa* -/-) or PTU treated larvae needed to be identified, either *Tg(Isl2b:RFP)*, or *Tg(Isl2b:GFP)* were used. In WT *Tg*(*isl2b*) drives expression in RGCs, trigeminal ganglia, and spinal neurons. In *lakritz*, there are no RGCs, but *Tg*(*isl2b*) still drives expression in the trigeminal ganglia and spinal neurons. In adults, fin clips were used to determine carriers of the *lakritz* mutation as described in (Kay et al. Neuron 2001). In short myTaq extract-PCR kit (Bioline) was used to amplify a 300 bp fraction of the *ath5* gene containing the *lakritz* mutation using the following primers: *Fw_ccggaattacatcccaagaac, Rv_ ggccatgatgtagctcagag*. The amplified product was digested using the StuI restriction enzyme over night at 37°C. Carriers of the *lakritz* mutation would show three products on an agarose gel at: 300 bp, 200 bp, and 100 bp. WT fish show two products at: 200 bp and 100 bp.

## Figure legends

*<u>Supplementary Figure 1. HCR-FISH validates HGn12C marker expression and specificity</u>*

*Panels organized by gene expression from Fig. 2. 2D UMAP expression plots are presented side by side with co-expression plots. Upper row shows UMAPs for glutamatergic neurons. Bottom rows show UMAPs for GABAergic neurons*.

*<u>Supplementary Figure 2. HCR-FISH stains for HGn12C markers</u>*

*Maximum projections color-coded by depth for each HGn12C marker gene. All stains were registered using the HGn12C expression pattern. Last panel shows colors depth scale in microns.*

*<u>Supplementary Figure 3. Heatmap of HCR-FISH signal in MapZeBrain brain regions</u>*

*(a) Total intensity detected from gene expression in each of the anatomical brain regions defined in the MapZeBrain atlas. The expression of each gene is normalized to the area with highest expression. (b) Same as in (a), but the signal is first normalized to the background intensity. (c) Values are same as in (b), but normalized in the end to the brightest intensity of each brain area. The heat map shows for each brain area genes best expressed in that area.*

*<u>Supplementary Table 1. Table of top hits for HCR-FISH stains from the Max Planck Zebrafish Brain Atlas</u>*

*<u>Supplementary Figure 4. Batch effects in WT population are limited and properly corrected</u>*

*(a) First 5 principal components of WT cells plotted in order and in all combinations. Color coding and numbering refers to all single-cell samples prepared on the same day. (b) UMAP embedding of all WT cells split according to day of sample preparation. Numbers are the same as in (a). (c) UMAP embedding of all cells before (left) and after (right) batch correction using Harmony. Numbers and colors are the same as in (a).*

*<u>Supplementary Figure 5. Lakritz (no RGCs) and WT cells can be fully integrated with no observable confounding effects</u>*

*(a) First 5 principal components of lakritz (no RGCs) cells plotted in order and in all combinations. Color coding and numbering refers to all single-cell samples prepared on the same day. (b) UMAP embedding of all lakritz (no RGCs) cells without batch correction. Numbers and colors are the same as in (a). (c) Embedding of WT and lakritz (no RGCs) cells in the same UMAP space without batch correction.*

*<u>Supplementary Figure 6. Independent clustering of glutamatergic neurons uncovers similar clusters across samples</u>*

*UMAP embedding of three different samples (unrelated WT, top; sibling WT, middle; lakritz, bottom) after independently processing and clustering each sample. For detailed cluster differences, see Supplementary Fig. S6 and methods.*

*<u>Supplementary Figure 7. Independent clustering of glutamatergic neurons uncovers similar clusters across samples</u>*

*Gene expression plots in UMAP embedded cells after independent processing and clustering of three different samples (unrelated WT, sibling WT, lakritz). The expression plots show the expression of marker genes for clusters which are not named across all three samples. (a-e) marker genes for clusters which are named by two markers. Green and red are the expression of individual markers (unrelated WT, top; sibling WT, middle; lakritz, bottom). (f,g) marker genes for clusters which are named by a single marker (unrelated WT, left; sibling WT, middle; lakritz, right). For detailed cluster differences, see methods.*

*<u>Supplementary Figure 8. Independent clustering of GABAergic neurons uncovers similar clusters across samples</u>*

*UMAP embedding of three different samples (unrelated WT, top; sibling WT, middle; lakritz, bottom) after independently processing and clustering each sample. For detailed cluster differences, see Supplementary Fig. S8 and methods.*

*<u>Supplementary Figure 9. Independent clustering of GABAergic neurons uncovers similar clusters across samples</u>*

*Gene expression plots in UMAP embedded cells after independent processing and clustering of three different samples (unrelated WT, sibling WT, lakritz). The expression plots show the expression of marker genes for clusters which are not named across all three samples. (a-c) marker genes for clusters which are named by two markers. Green and red are the expression of individual markers (unrelated WT, top; sibling WT, middle; lakritz, bottom). (d-p) marker genes for clusters which are named by a single marker (unrelated WT, left; sibling WT, middle; lakritz, right). For detailed cluster differences, see methods.*

*<u>Supplementary Figure 10. Larger WT population does not coerce lakritz (no RGCs) cell-type identities to resemble WT identities</u>*

*Lakritz (no RGCs) cells embedded with all other cells. Color coding is drawn from cluster identity following clustering of only lakritz (no RGCs) cells. Shown are only lakritz cells in WT-lakritz mutual UMAP space.*

*<u>Supplementary Figure 11. In-silico cell type ablation of a subset of glutamatergic neuronal clusters</u>*

*A select number of clusters processed via an in-silico cell ablation pipeline. (a-c) Example of three clusters in which we could positively identify which cell type was ablated. (d) Example of a cluster where we were unable to identify which cell type was ablated.*

*<u>Supplementary Figure 12. In-silico cell type ablation of all glutamatergic neuronal clusters</u>*

*Panels organized in sequential pairs. Left panel of each cluster shows UMAP embedding of all glutamatergic neurons. In red are all WT cells belonging to the ablated (omitted) cluster. Right panel shows the same UMAP as the left panel, but with color-coding representing the experimental genotype (red, unrelated WT; green, sibling WT; blue, lakritz (no RGCs))*

*<u>Supplementary Figure 13. In-silico cell type ablation of a subset of GABAergic neuronal clusters</u>*

*A select number of clusters processed via an in-silico cell ablation pipeline. (a-c) Example of three clusters in which we could positively identify which cell type was ablated. (d) Example of a cluster where we were unable to identify which cell type was ablated.*

*<u>Supplementary Figure 14. In-silico cell type ablation of all GABAergic neuronal clusters</u>*

*Panels organized in sequential pairs. Left panel of each cluster shows UMAP embedding of all GABAergic neurons. In red are all WT cells belonging to the ablated (omitted) cluster.*

*Right panel shows the same UMAP as the left panel, but with color-coding representing the experimental genotype (red, unrelated WT; green, sibling WT; blue, lakritz (no RGCs))*

*<u>Supplementary Figure 15. Lakritz (no RGCs) population shows a global transcriptomic drift from WT population</u>*

*(a) Volcano plots of global markers (gray dots) detected between unrelated WT and WT (left) and lakritz (no RGCs) and unrelated WT (right). Colored dots with text represent marker genes differentially expressed both in their level and in the proportion of cells expressing the gene (blue, control; red, lakritz; no RGCs). (b) PC similarity across groups. Red, iterative random shuffle of control cells into two groups generating a null distribution. Green, unrelated WT compared with WT. Blue, unrelated WT compared with lakritz (no RGCs). P-values calculated using a wilcoxon signed-rank test. (c) UMAP embedding of all cells. Color coding shows cells with altered nearest-neighbor neighborhoods. Color intensity shows Z-score for neighborhood alteration. P-values calculated using a wilcoxon signed-rank test comparing either WT (left, green) or lakritz (no RGCs; right, red) to a random label shuffle distribution.*

*<u>Supplementary Figure 16. There is no significant change in relative cluster proportions in absence of RGCs</u>*

*Bar plots showing for each cluster the variance in cluster’s relative percentage across replicates (a, glutamatergic clusters; b, GABAergic clusters). For each cluster, the variance is shown for all genotypes (red, unrelated WT; green, sibling WT; blue, lakritz (no RGCs)). P-values calculated using a wilcoxon signed-rank test and corrected for multiple testing using the Bonferroni correction.*

*<u>Supplementary Figure 17. A subset of clusters shows altered transcriptome despite correct fate in absence of RGCs</u>*

*(a) Left, UMAP embedding of WT glutamatergic cells color-coded by cluster. Right, cells embedded in UMAP space, same as on the left, color coded by the p-value exponent calculated from analysis measuring cluster-specific transcriptome changes. (b) same as in (a), but for GABAergic cells.*

*<u>Supplementary Figure 18. A small subset of markers shows morphologically-specific altered expression in absence of RGCs</u>*

*(a) Registration of Tg(HGn12C) expression patterns from WT (left, yellow), and lakritz (no RGCs; middle, magenta). Right panel shows registered expression patterns to standard HGn12C pattern. (b-e) Comparative stains between WT (left), and lakritz (no RGCs; right). Arrows (b,c,e) show areas where expression is altered in absence of RGCs. In (d) the tectum shows down-regulation.*

*<u>Supplementary Figure 19. Precursor neurons cluster markers reveal canonical differentiation pathways</u>*

*(a) UMAP embedding of all cells color-coding shows expression levels of neurog1 in precursor neurons differentiating into either glutamatergic neurons (including habenula neurons). (b) UMAP embedding of all cells color-coding shows expression levels of ascl1b in precursor neurons differentiating into GABAergic neurons. (c) UMAP embedding of all cells color-coding shows expression levels of cxcr4b in precursor neurons differentiating into habenula neurons.*

*<u>Supplementary Table 2. Putative differentiation markers</u>*

*Table of markers from neuron precursor cluster. Each gene is categorized according to its expression leading to a cluster (transient expression) or labeling in the cluster (maintained expression).*

*<u>Supplementary Figure 20. Progenitor and non-commited neuron markers are altered in absence of RGCs</u>*

*Comparative HCR in-situ stains performed on lakritz (right panels) and its WT siblings (left panels). Shown side-by-side are stains registered to Tg(HGn12C:GFP) standard brain color-coded by depth from dorsal (bright) to ventral (dark).*

*<u>Supplementary Figure 21. Optogenetic pretectal activation of OKR in lakritz is robust and specific</u>*

*(a) Control lakritz (no RGCs) larvae same as in (fig 5c,d), but lacking Chr2 expression. (b) Two more examples of lakritz (no RGCs) larvae as in (fig 5d). (c) RFLP analysis confirming a lakritz (no RGCs) mutation in larvae selected for pretectal optogenetic stimulation. From left to right: 1,2 lakritz; 3,4 WT controls. DNA ladder is in bp.*

*<u>Supplementary Figure 22. The Tg(HGn12C) pattern encompasses Tg(s1026t) pretectal neurons</u>*

*(a) UMAP embedding of scRNA-seq results collected from Tg(s1026t; red, left) together with all cells from WT Tg(HGn12C; cyan, left). On the right, color coding of elavl3 expression level across all cells in UMAP. (b) pretectal neurons are co-labeled by Tg(HGn12C) and Tg(Gal4s1026t). Shown is a pretectal plane from a single larva. In magenta Tg(Gal4s1026t, UAS:Chr2-mCherry). In green, Tg(HGn12C:GFP). Small panel shows transmitted light and field-of-view. All scale bars are 100 um. (c) Registered to standard brain images of transgenic lines Tg(HGn12C; cyan, left) and Tg(s1026t; red, middle). In magenta (right) mask of the pretectum over the standard brain (gray).*

