## Supplementary figures and images for "Retinal input influences pace of neurogenesis but not cell-type configuration of the visual forebrain"

### Supplementary Figure 1

##### Supplementary Figure 1

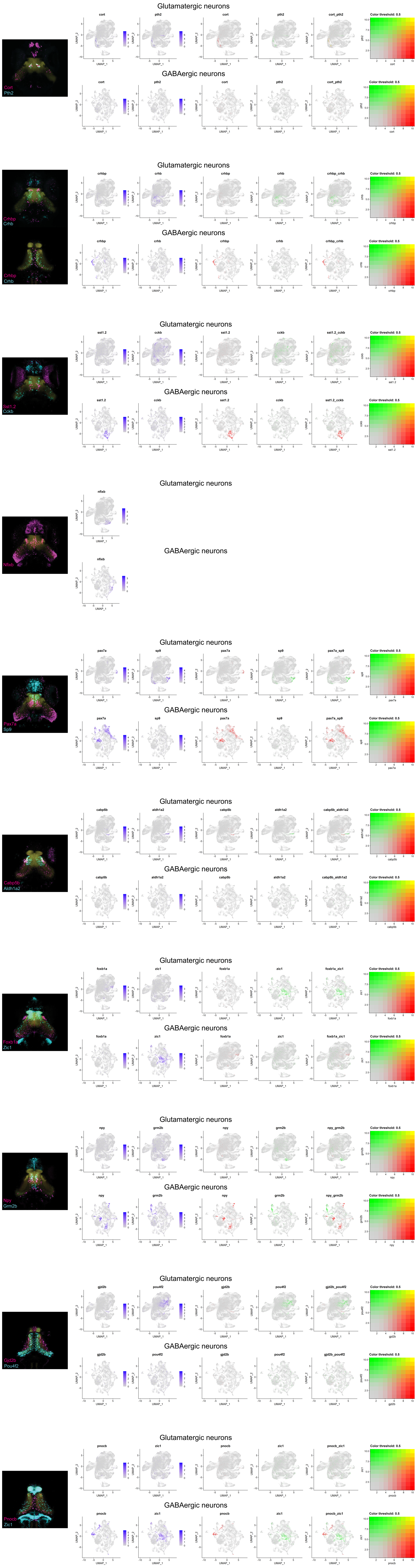

### Supplementary Figure 2

#### Supplementary Figure 2

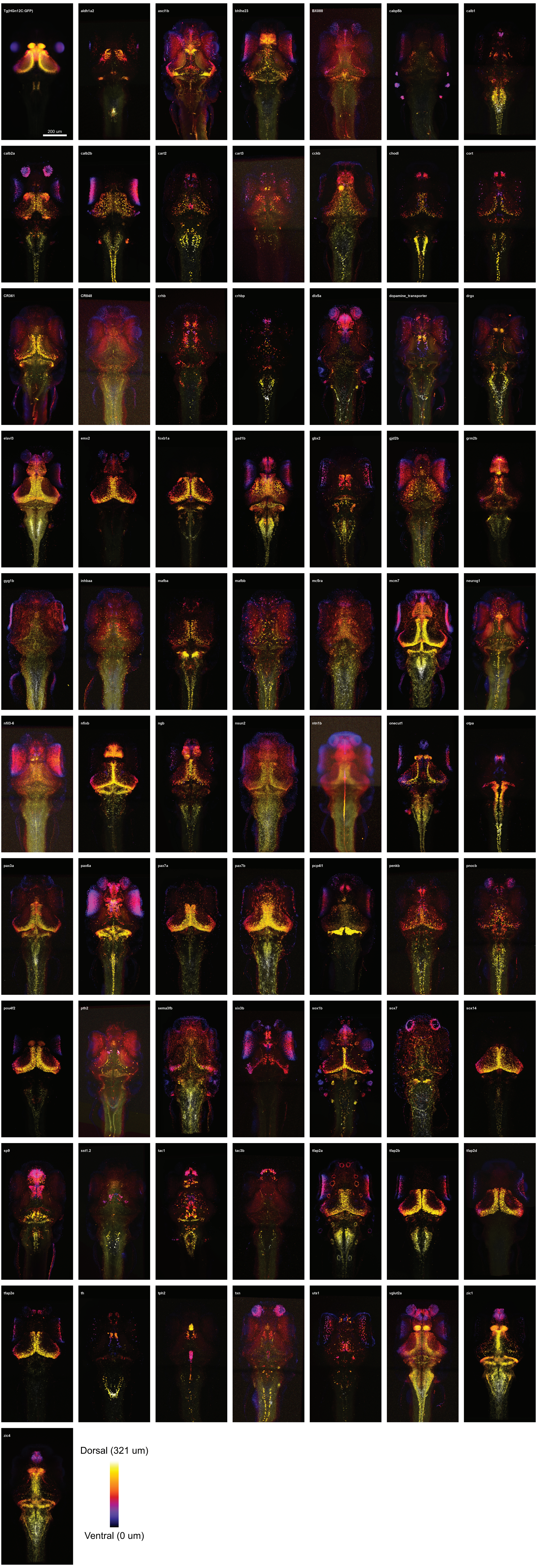

### Supplementary Figure 3

#### Supplementary Figure 3

a

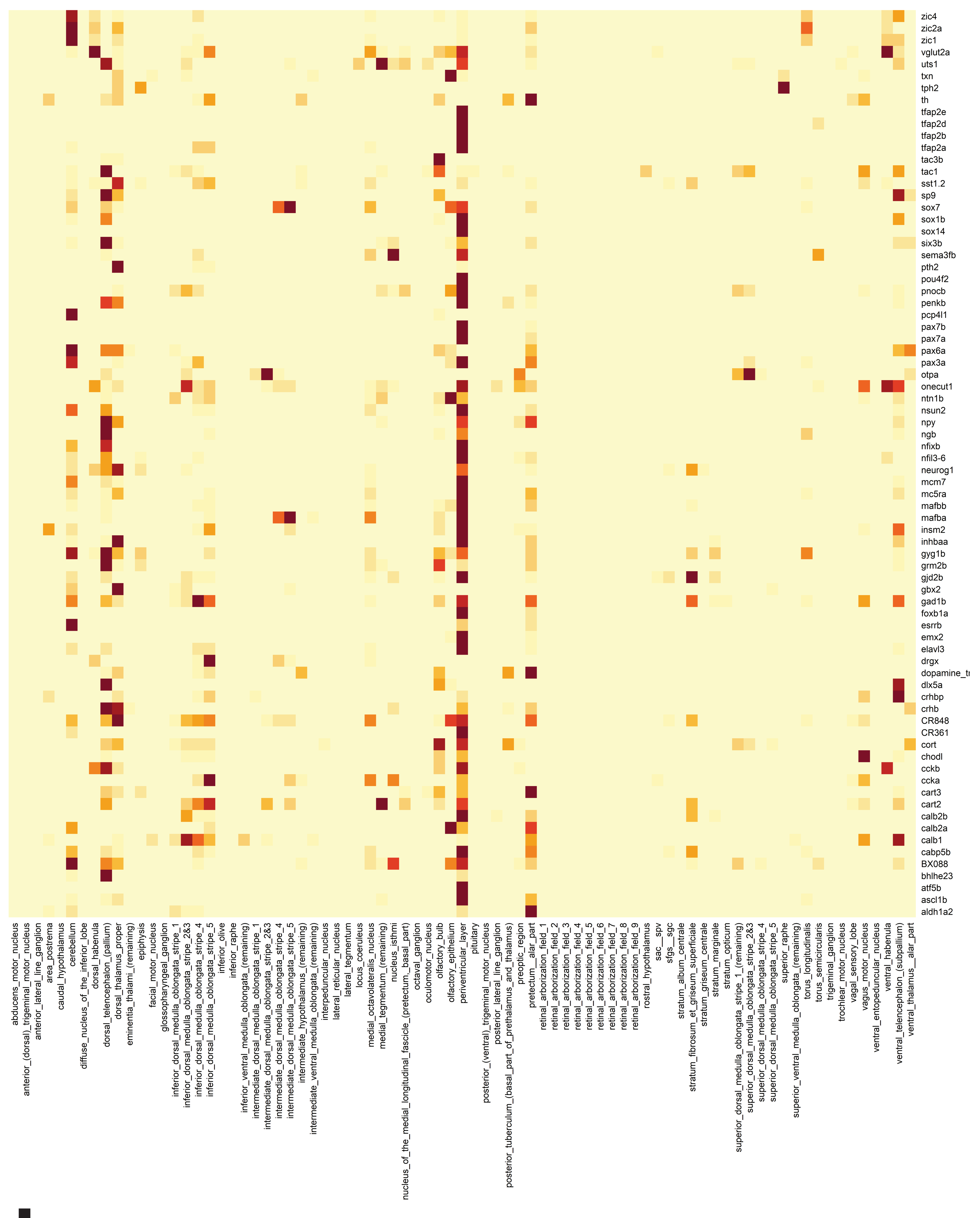

b

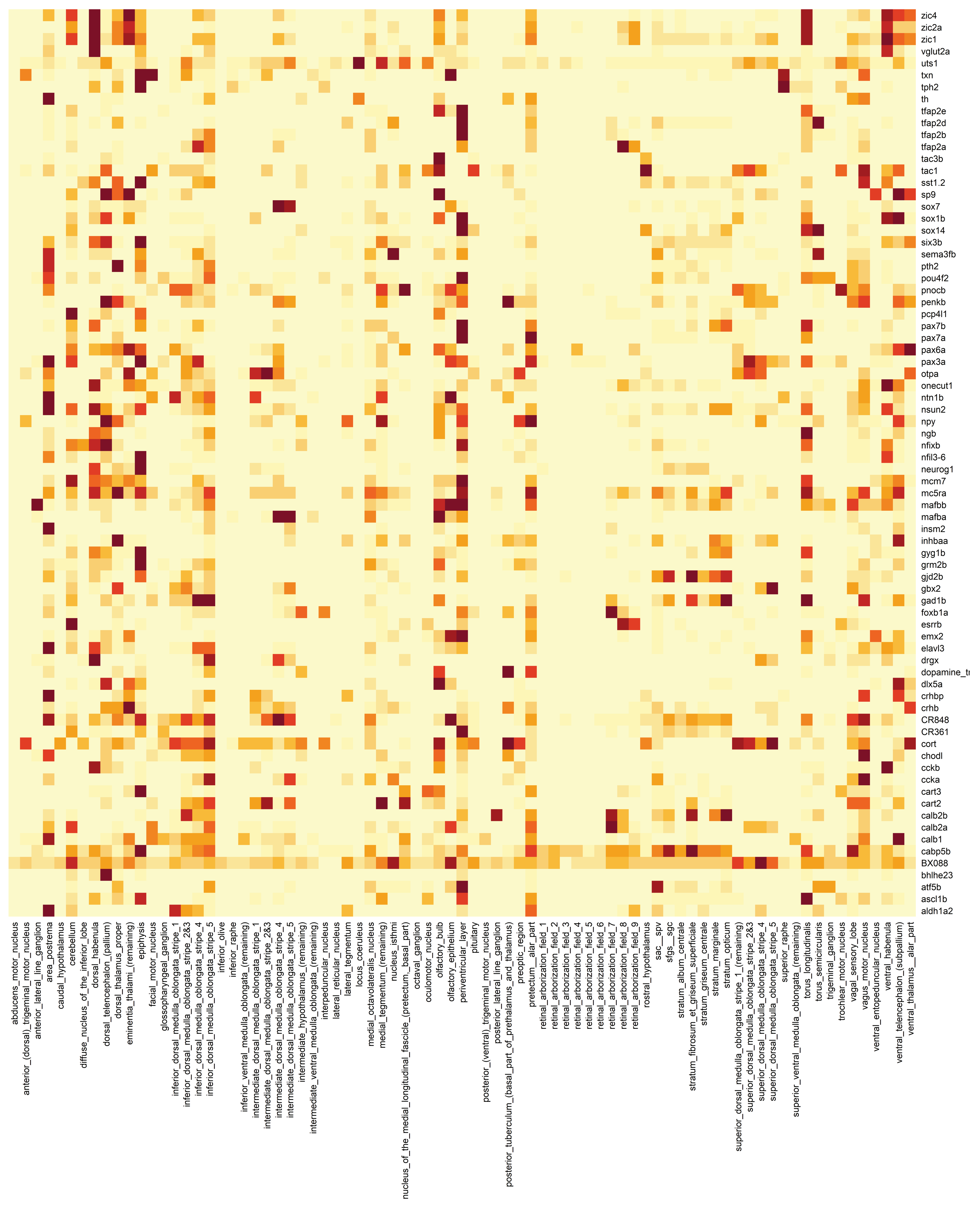

c

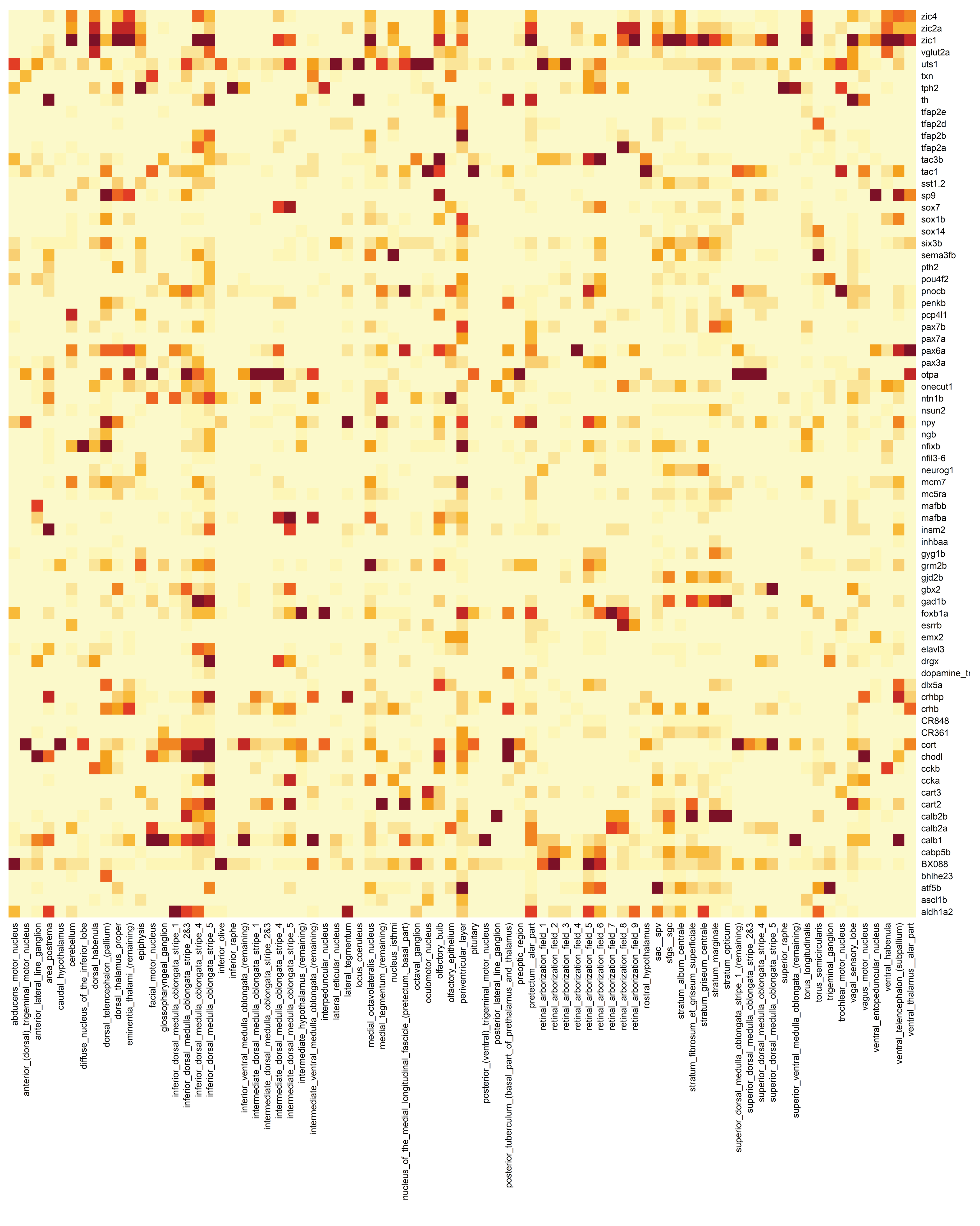

### Supplementary Figure 4

Supplementary Figure 4

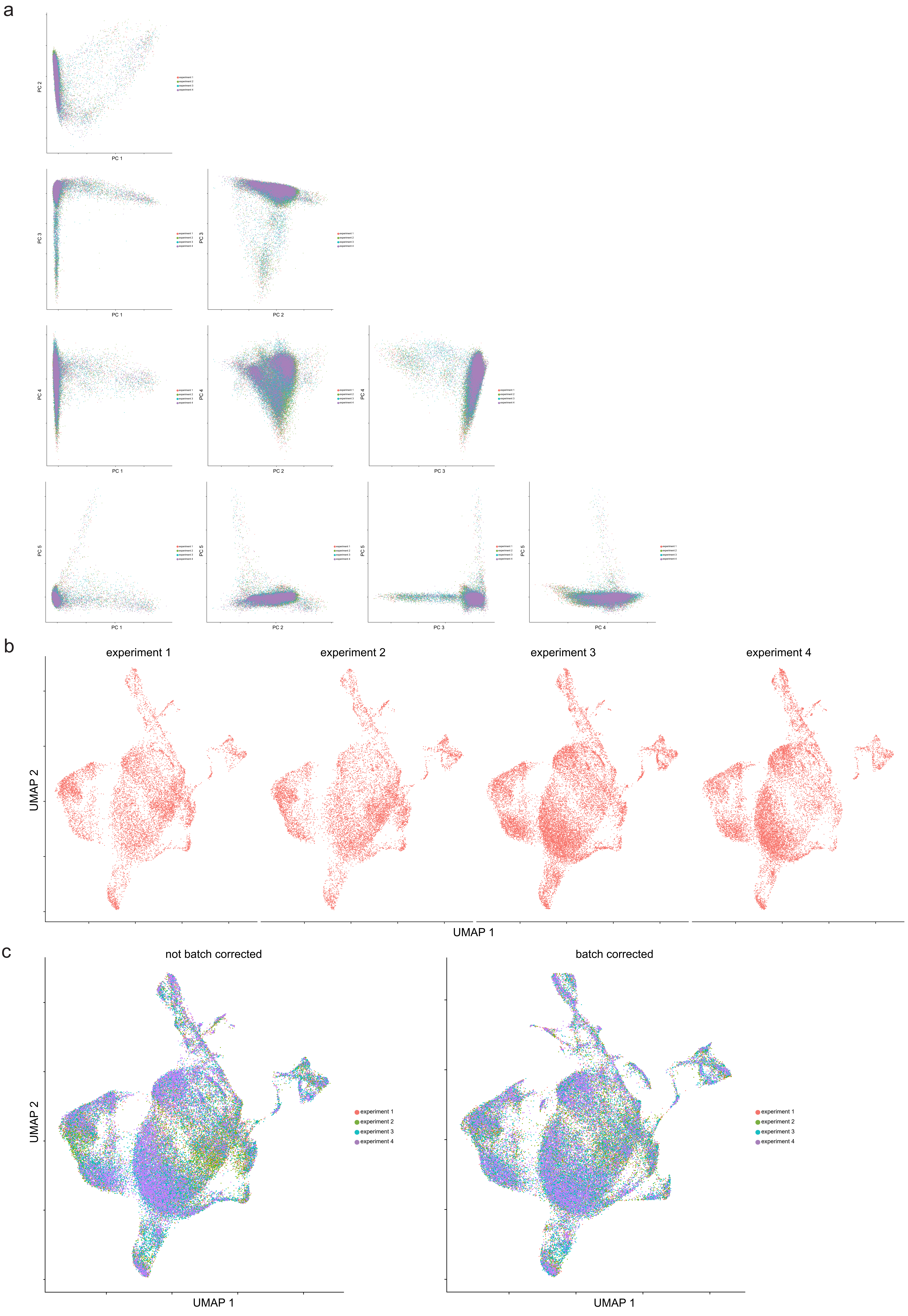

### Supplementary Figure 5

Supplementary Figure 5

a

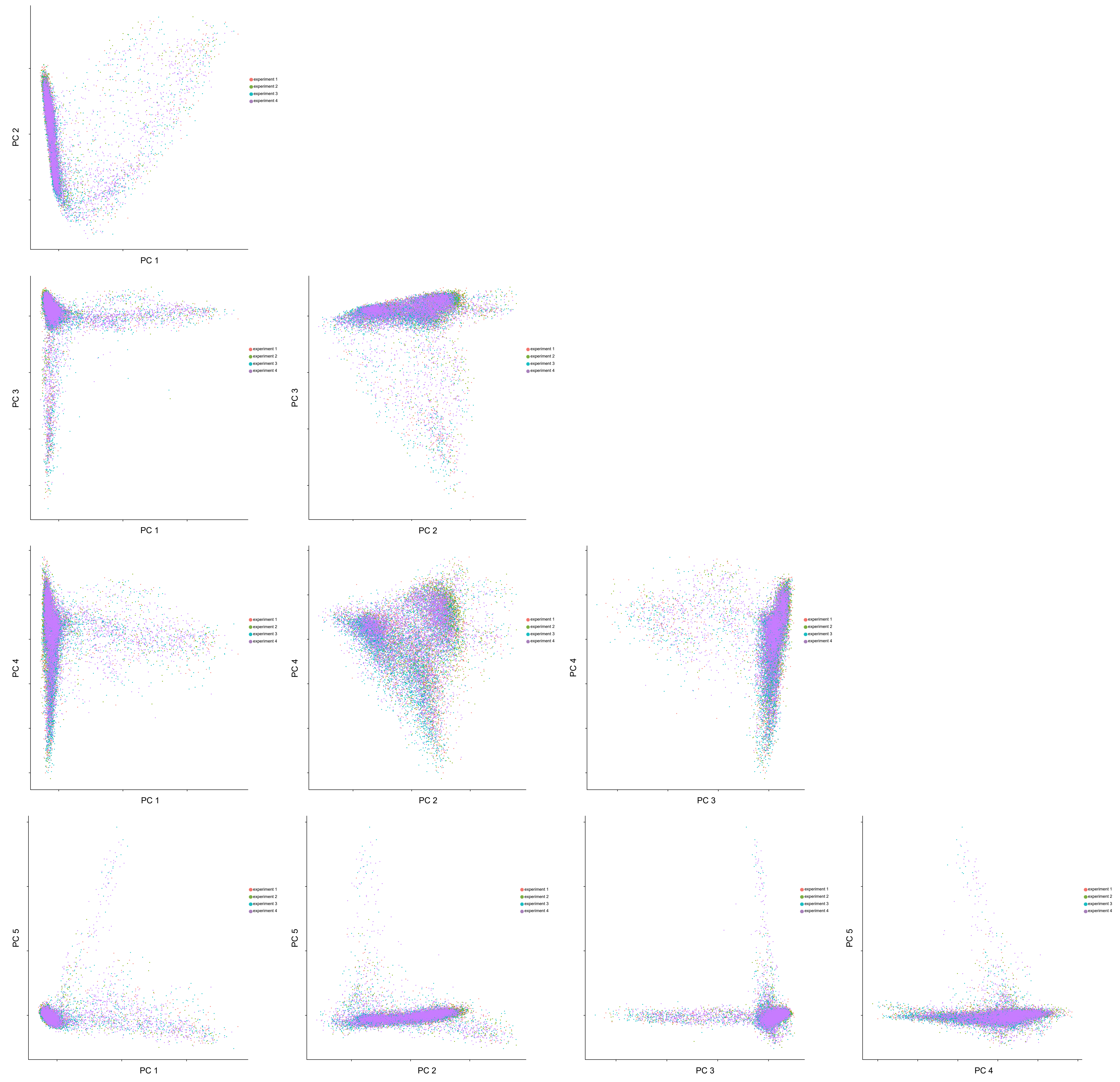


b

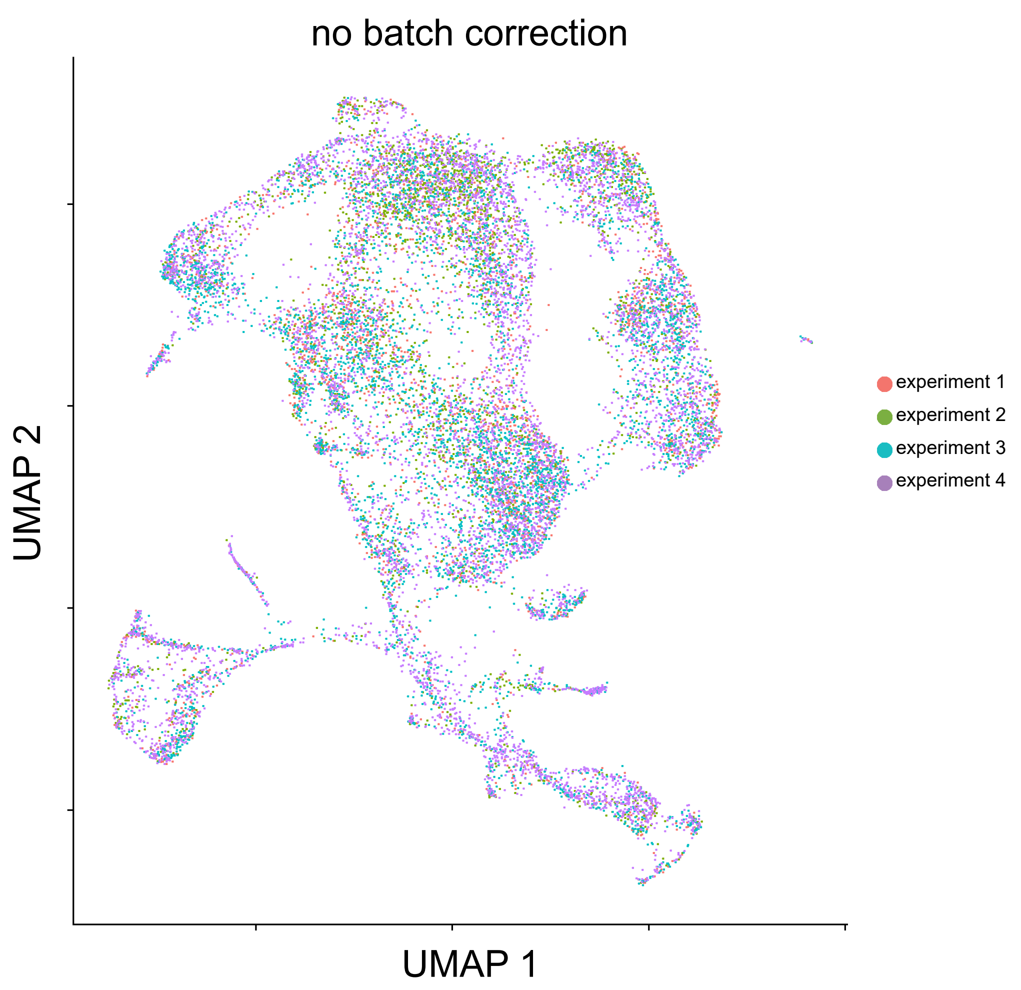


c

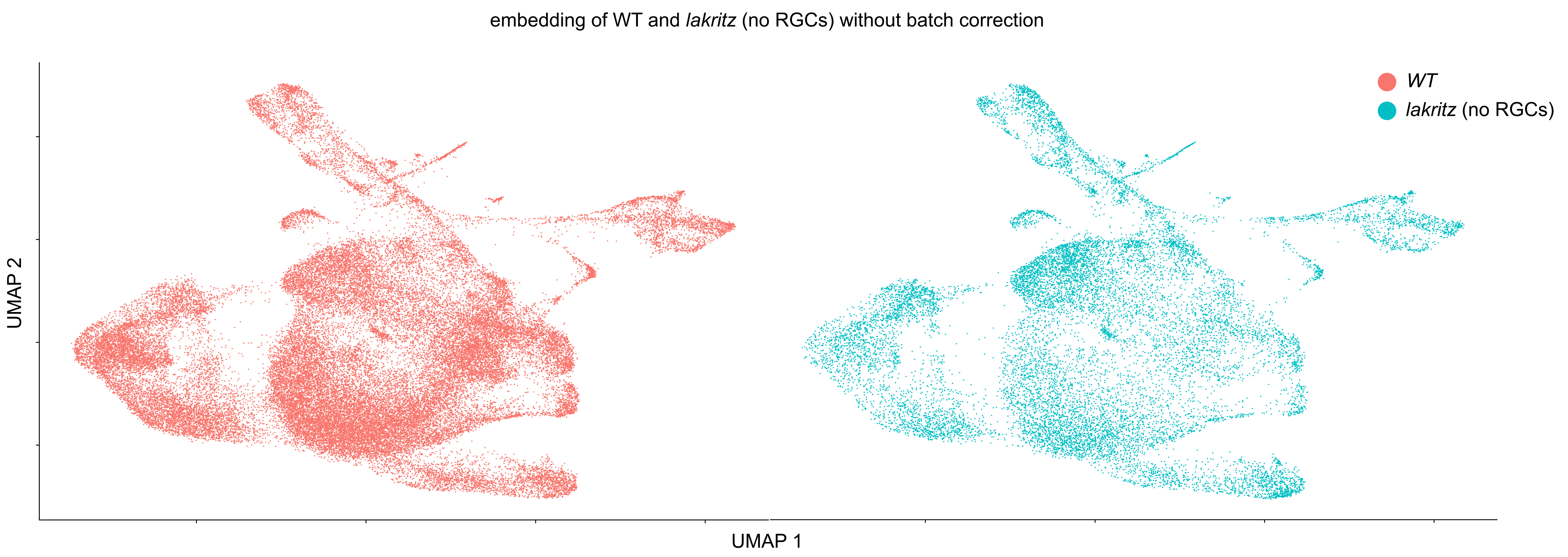

### Supplementary Figure 7

#### Supplementary Figure 7

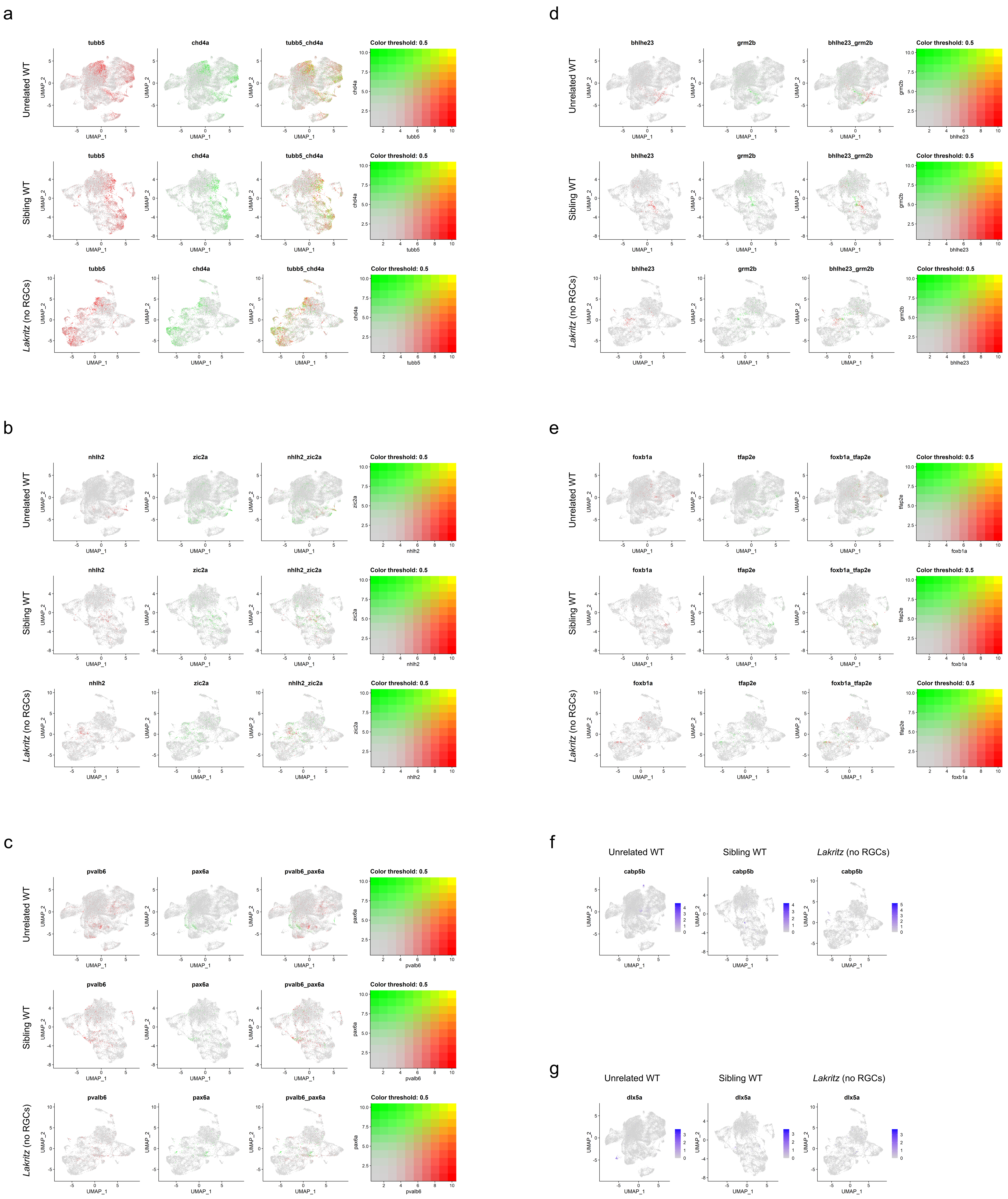

### Supplementary Figure 9

Supplementary Figure 9

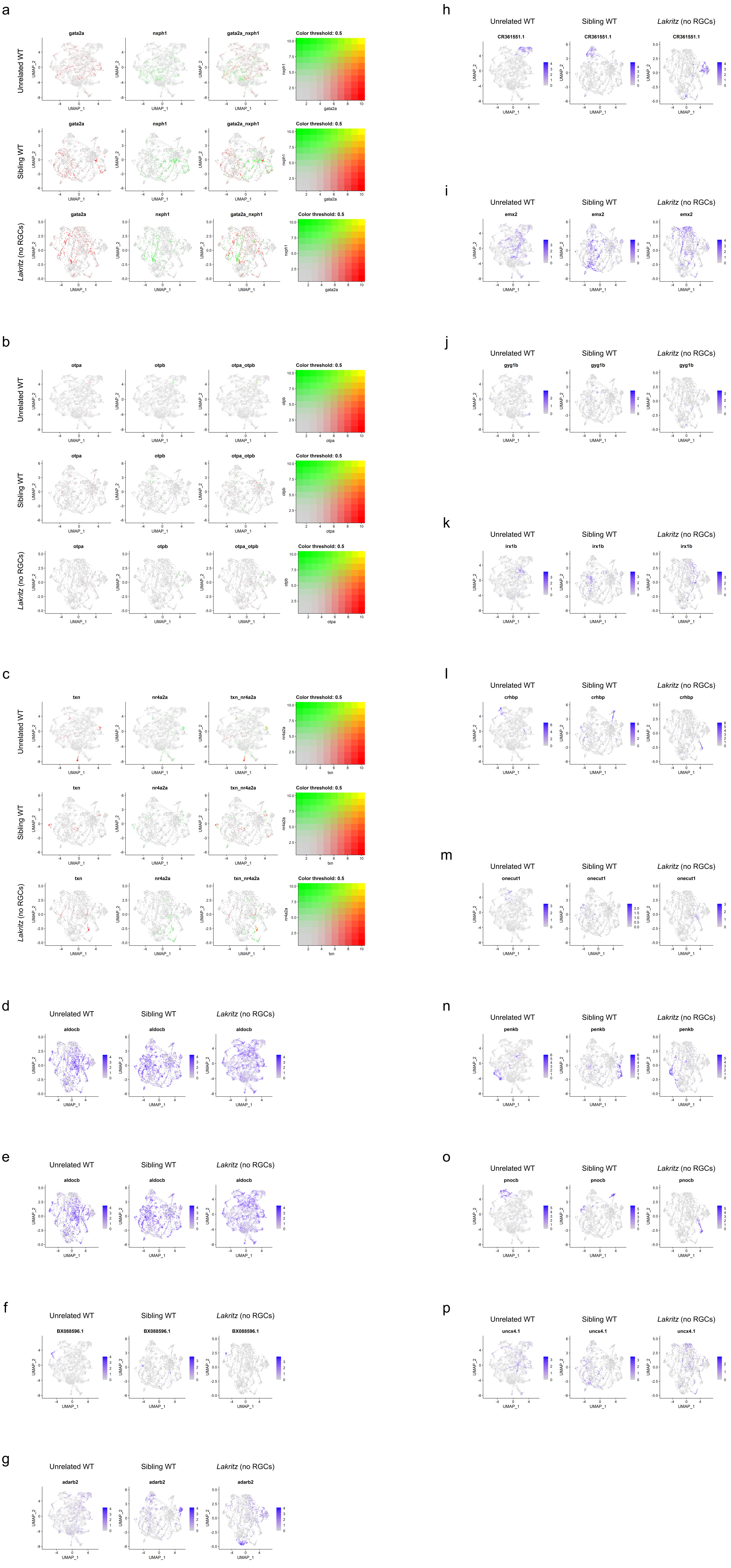

### Supplementary Figure 10

Supplementary Figure 10

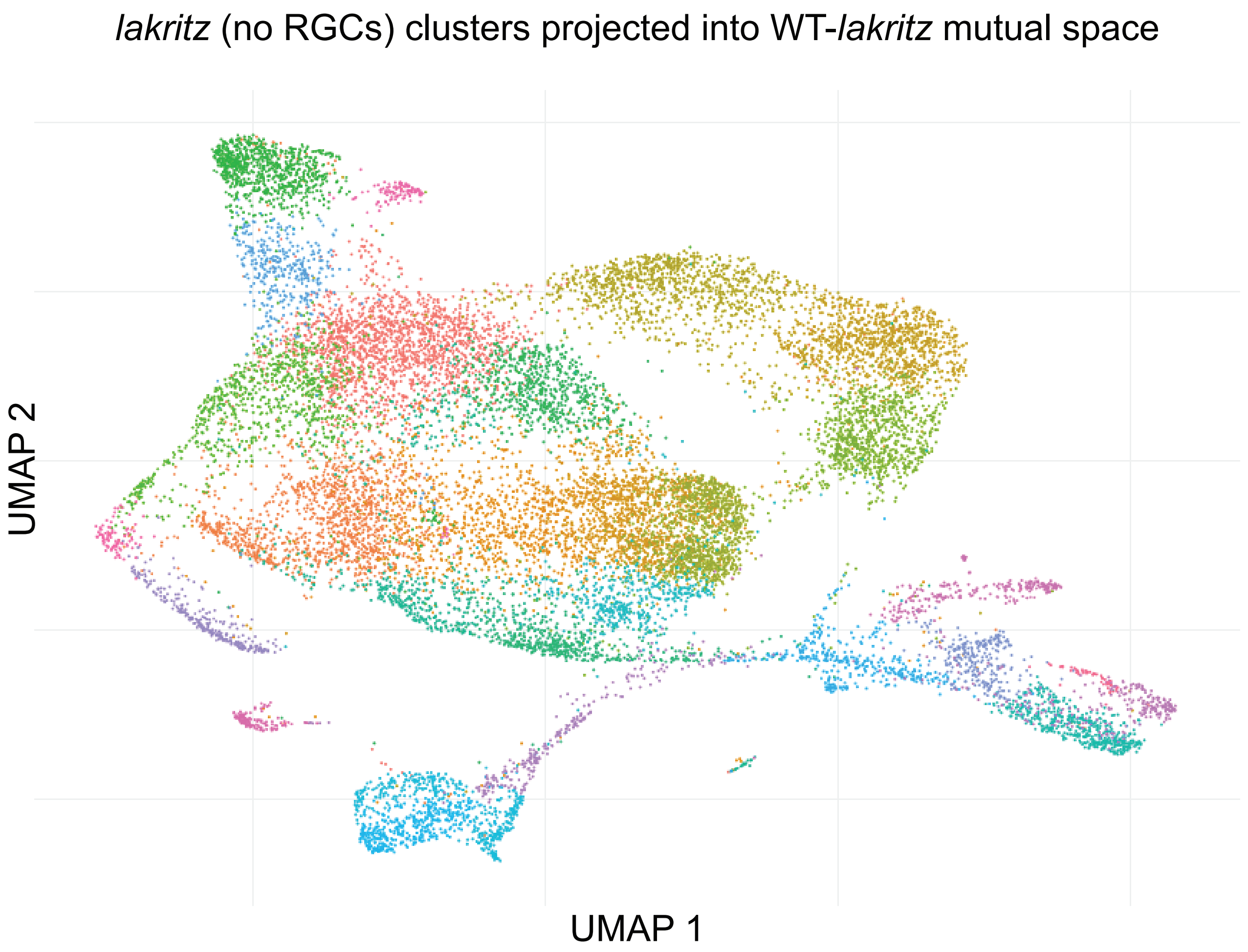

### Supplementary Figure 11

#### Supplementary Figure 11

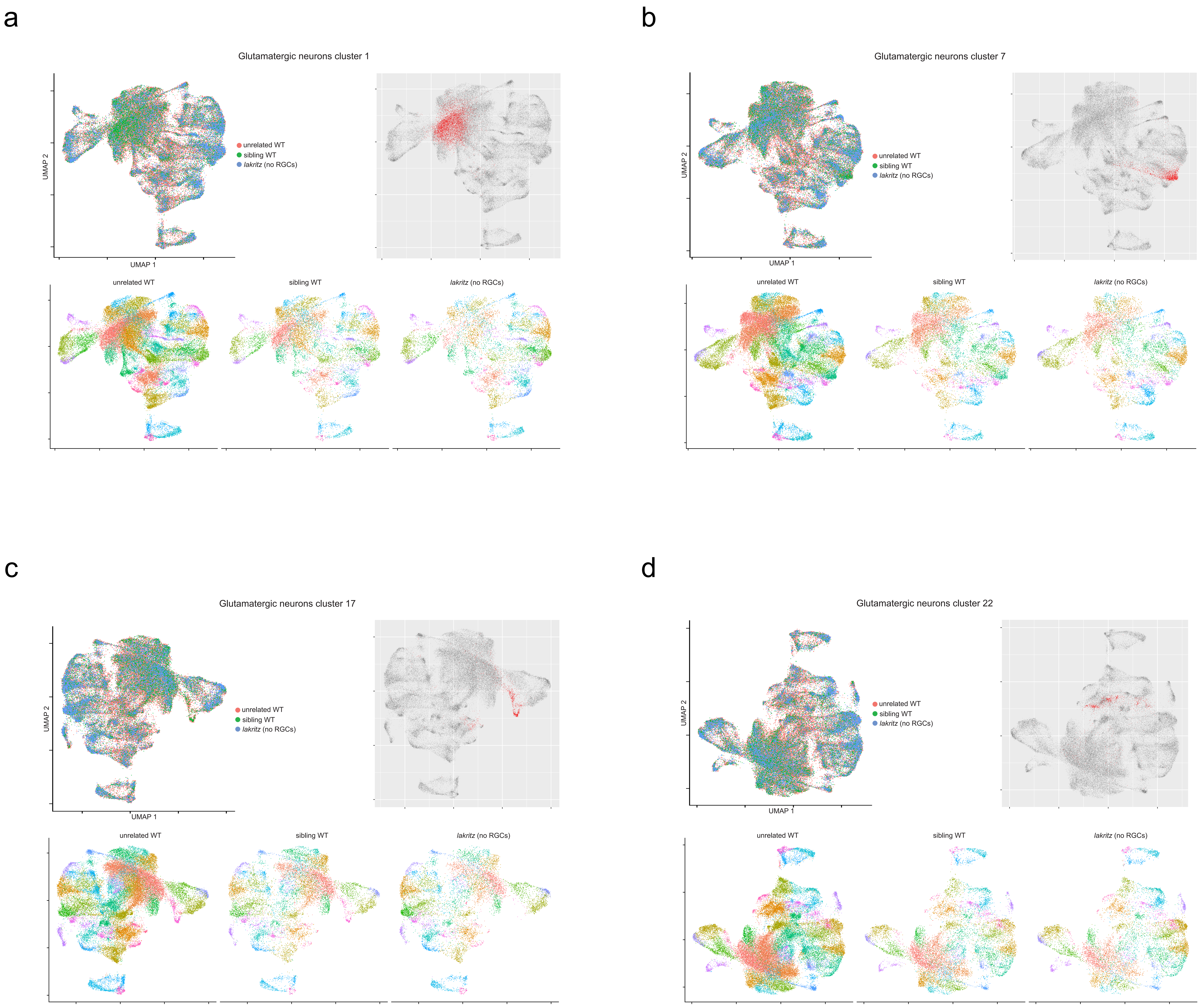

### Supplementary Figure 12

#### Supplementary Figure 12

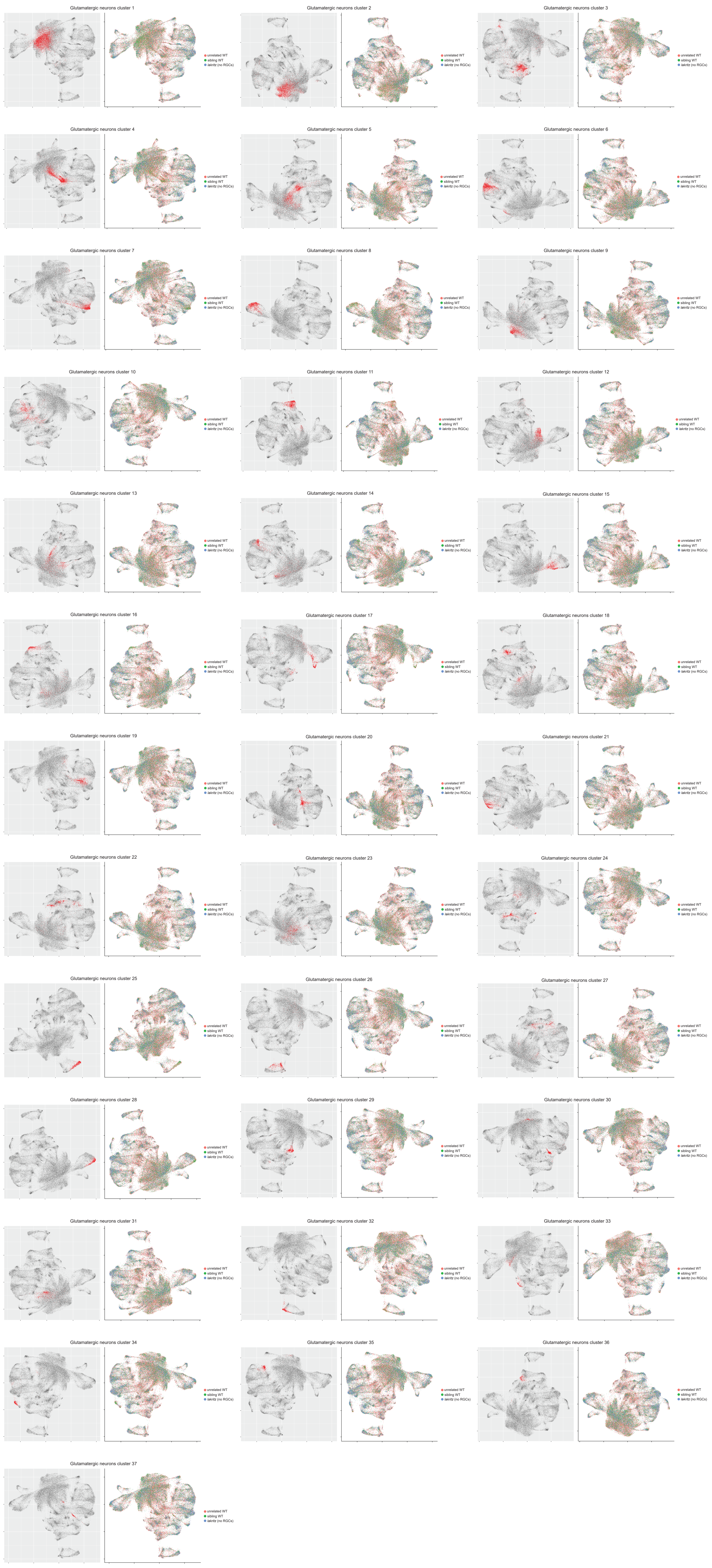

### Supplementary Figure 13

#### Supplementary Figure 13

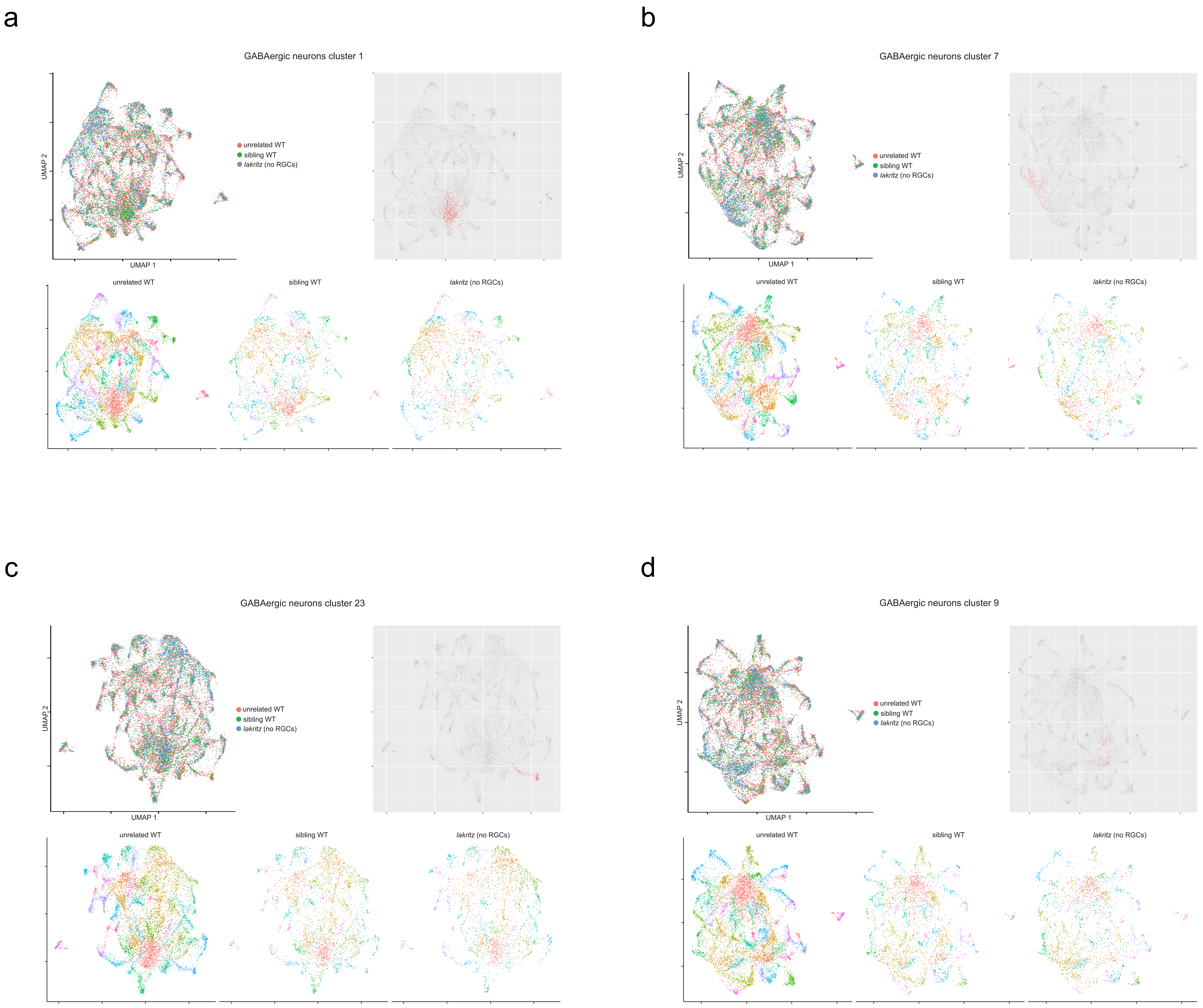

### Supplementary Figure 14

#### Supplementary Figure 14

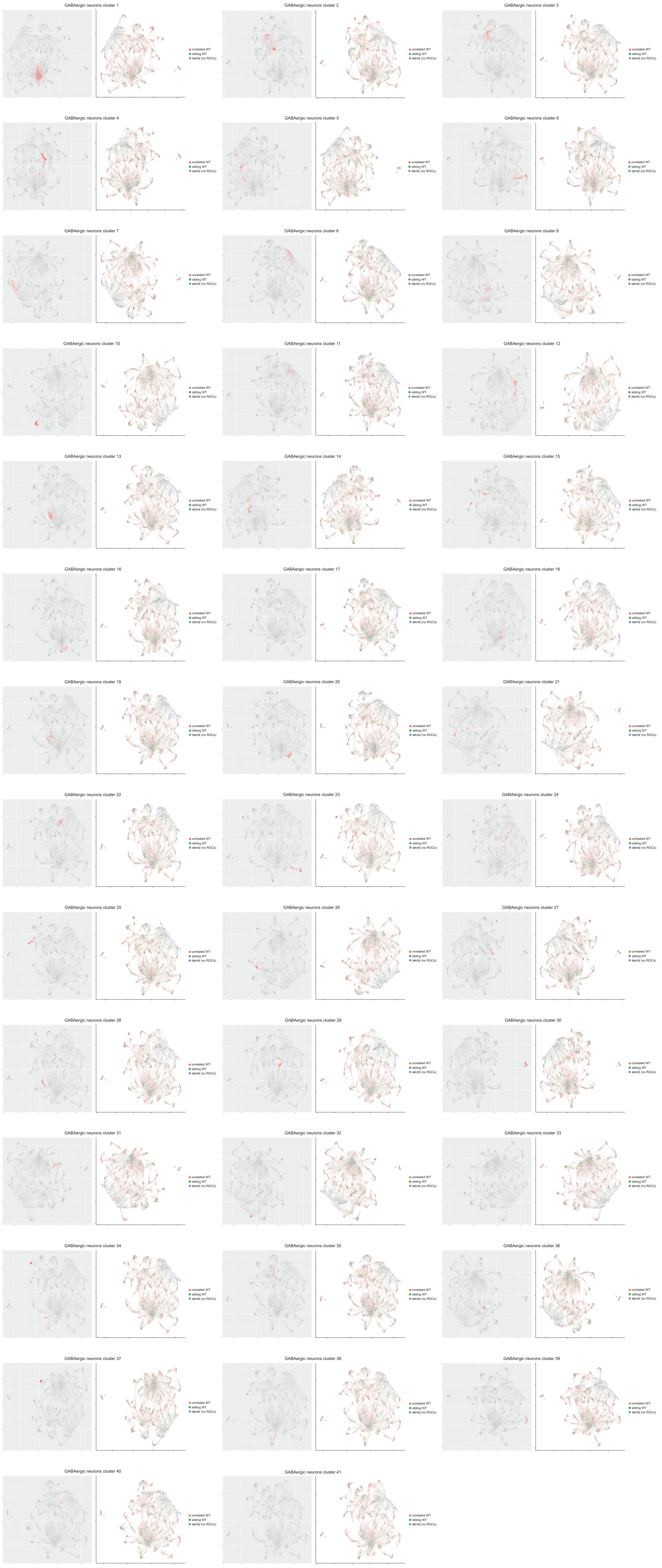

### Supplementary Figure 15

#### Supplementary Figure 15

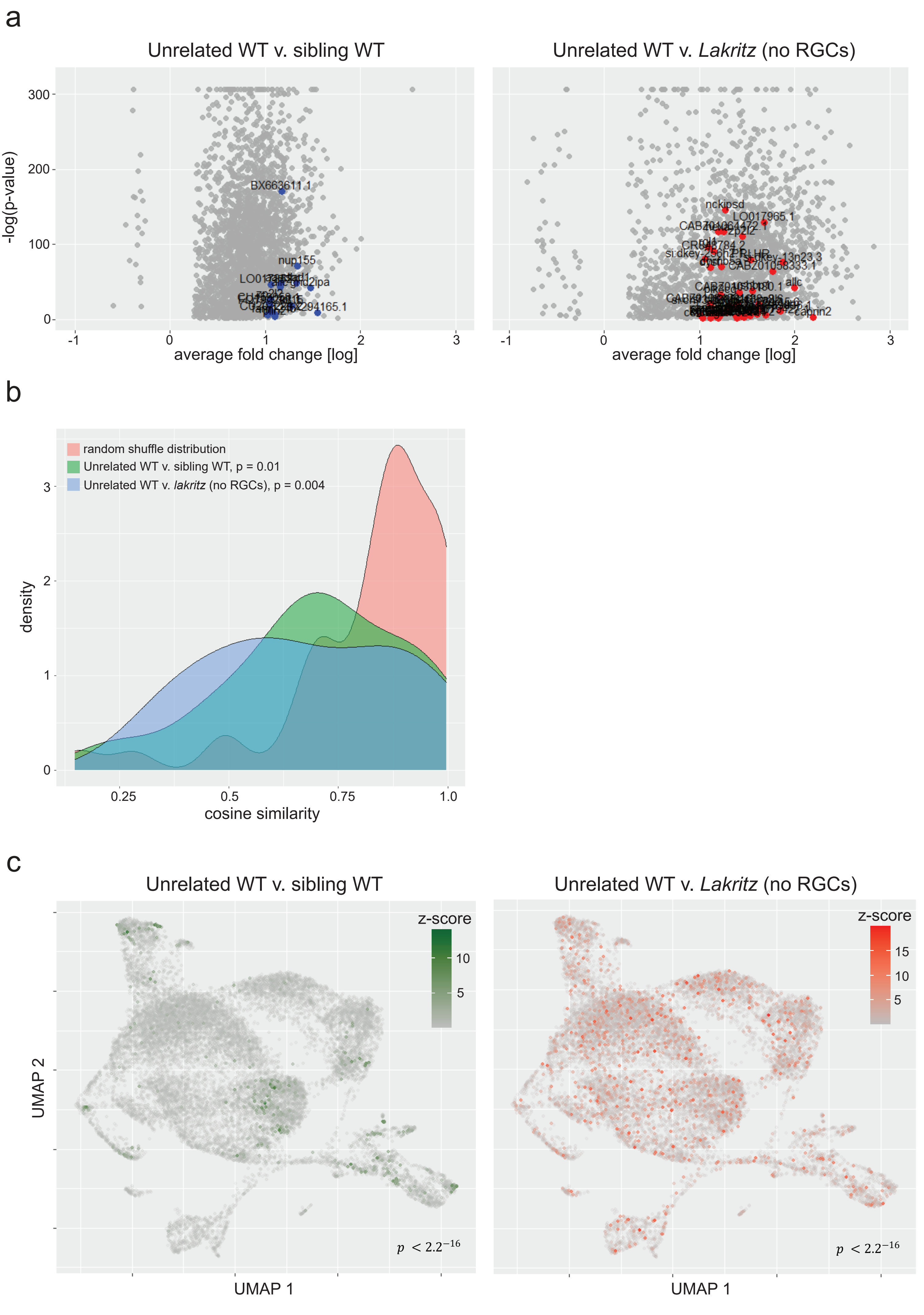

### Supplementary Figure 16

#### Supplementary Figure 16

a

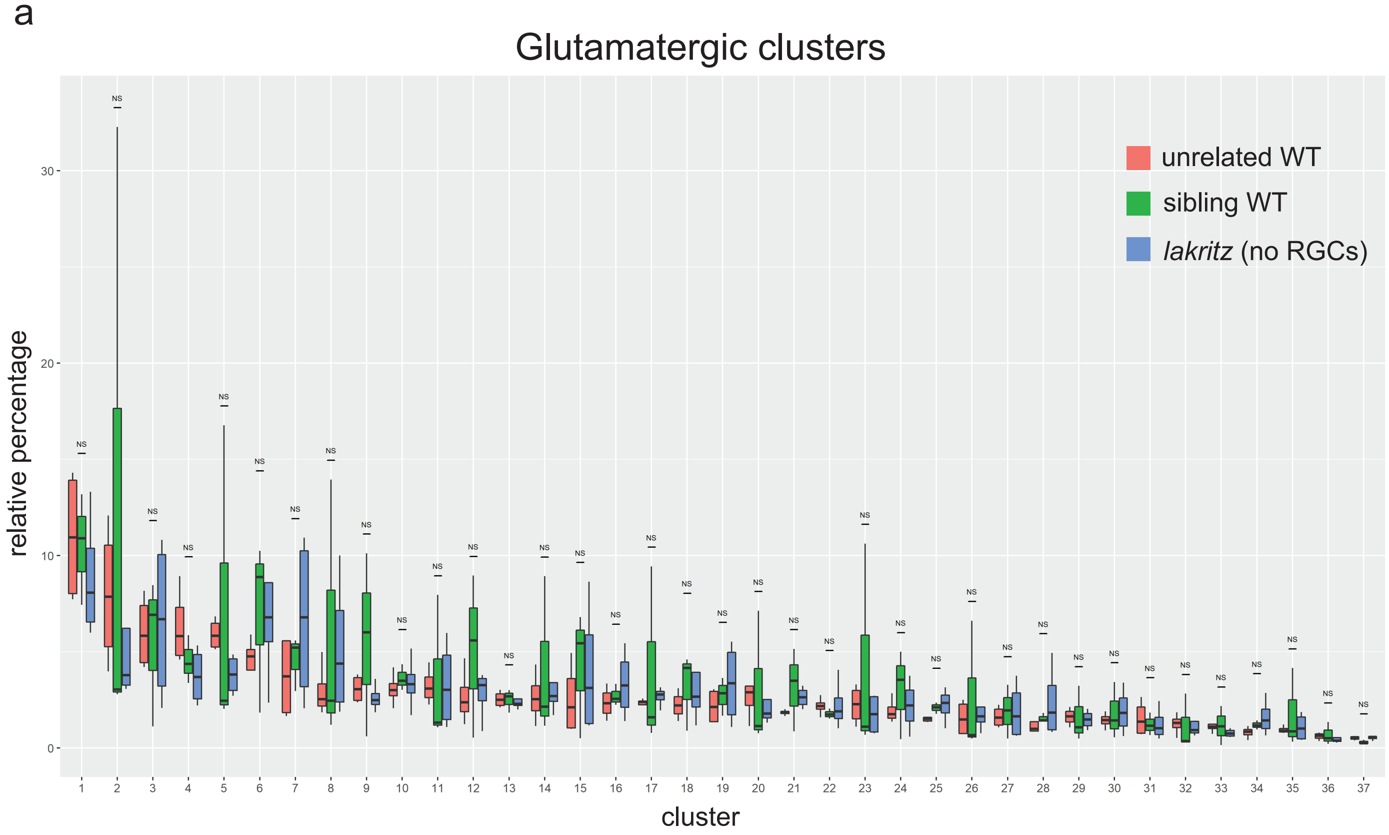


b

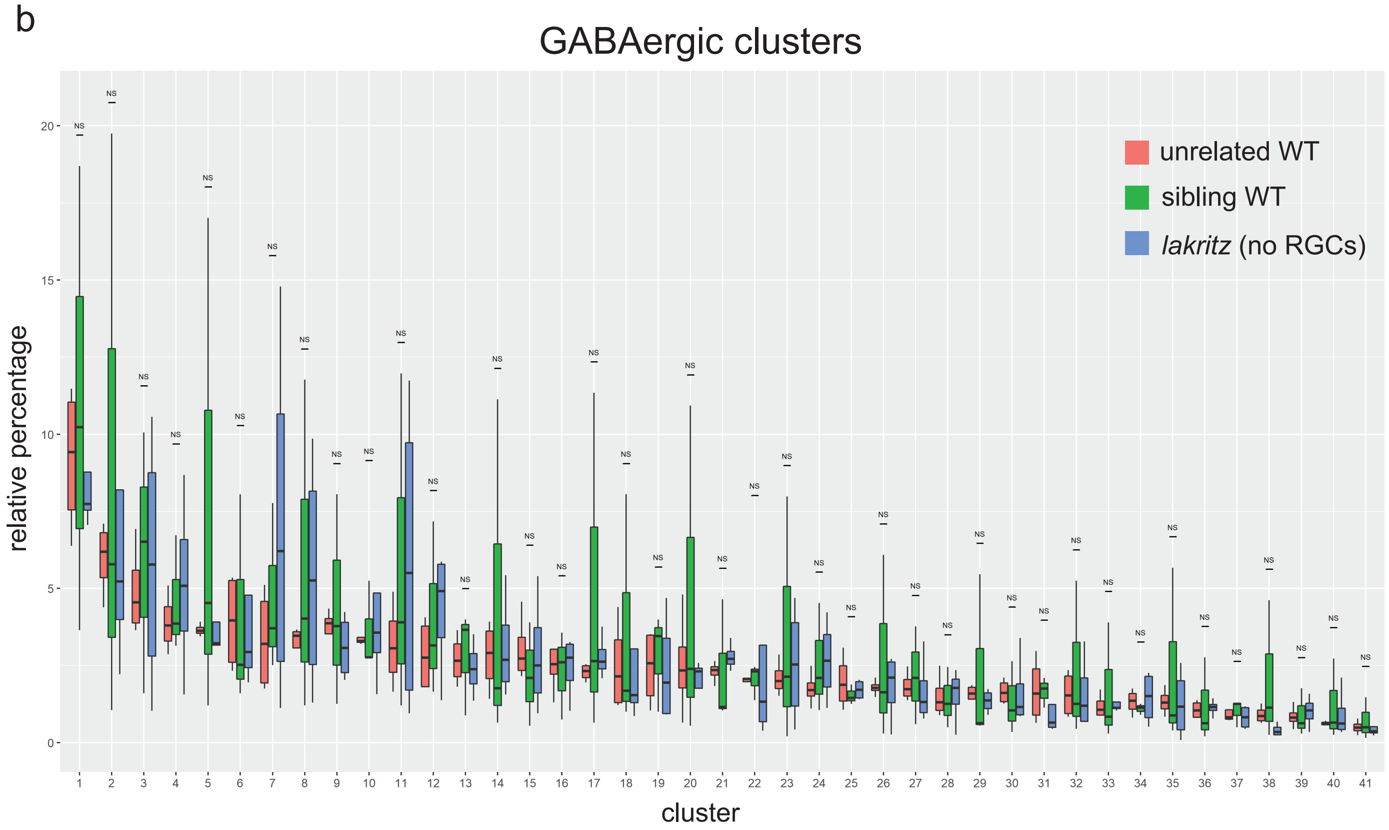

### Supplementary Figure 17

#### Supplementary Figure 17

a

Glutamatergic neurons

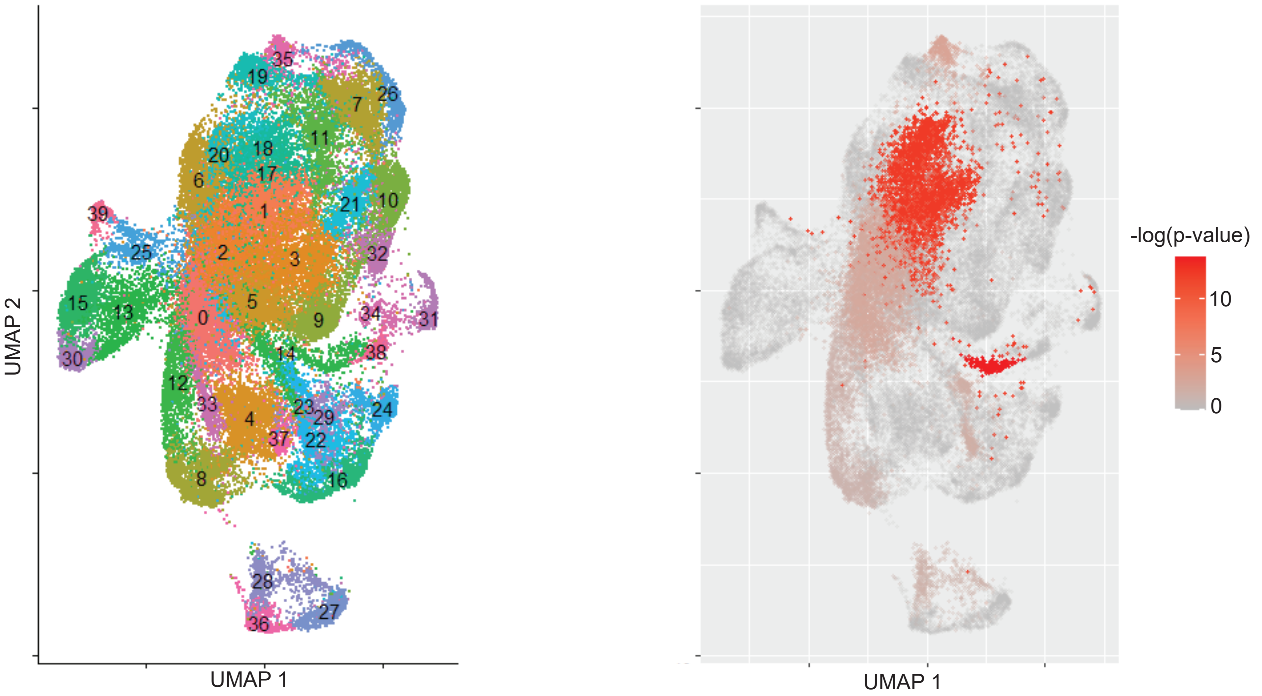


b

GABAergic neurons

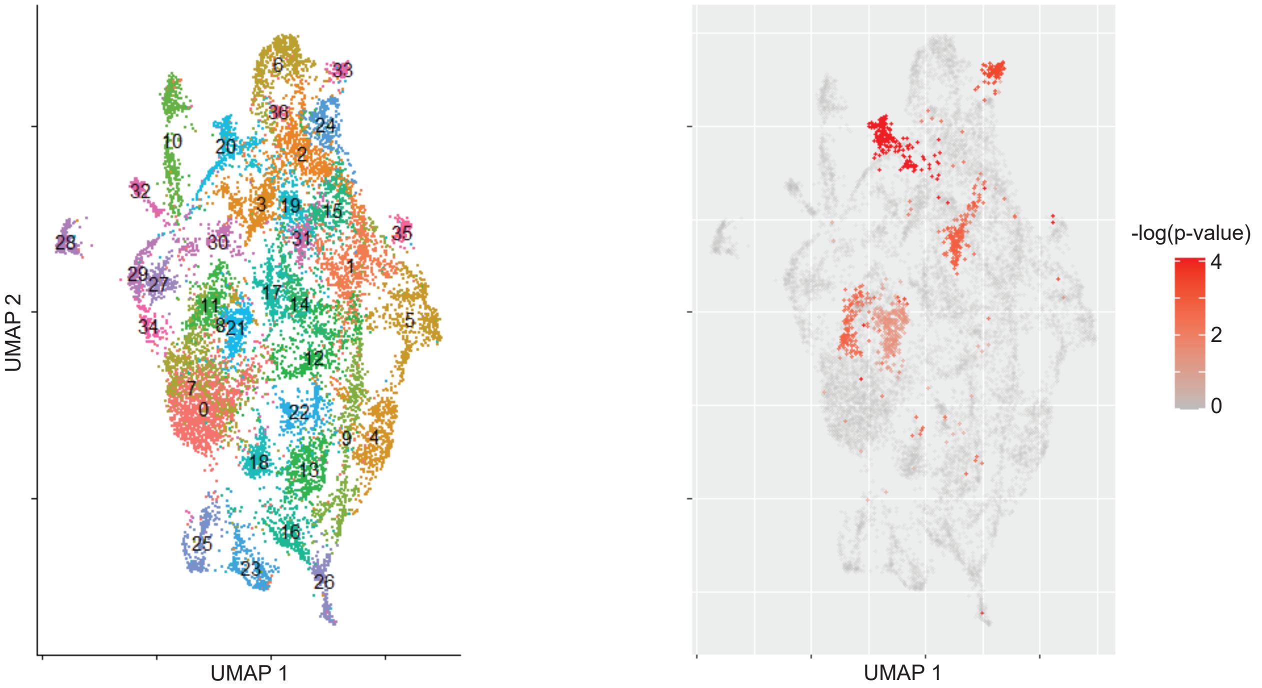

### Supplementary Figure 18

#### Supplementary Figure 18

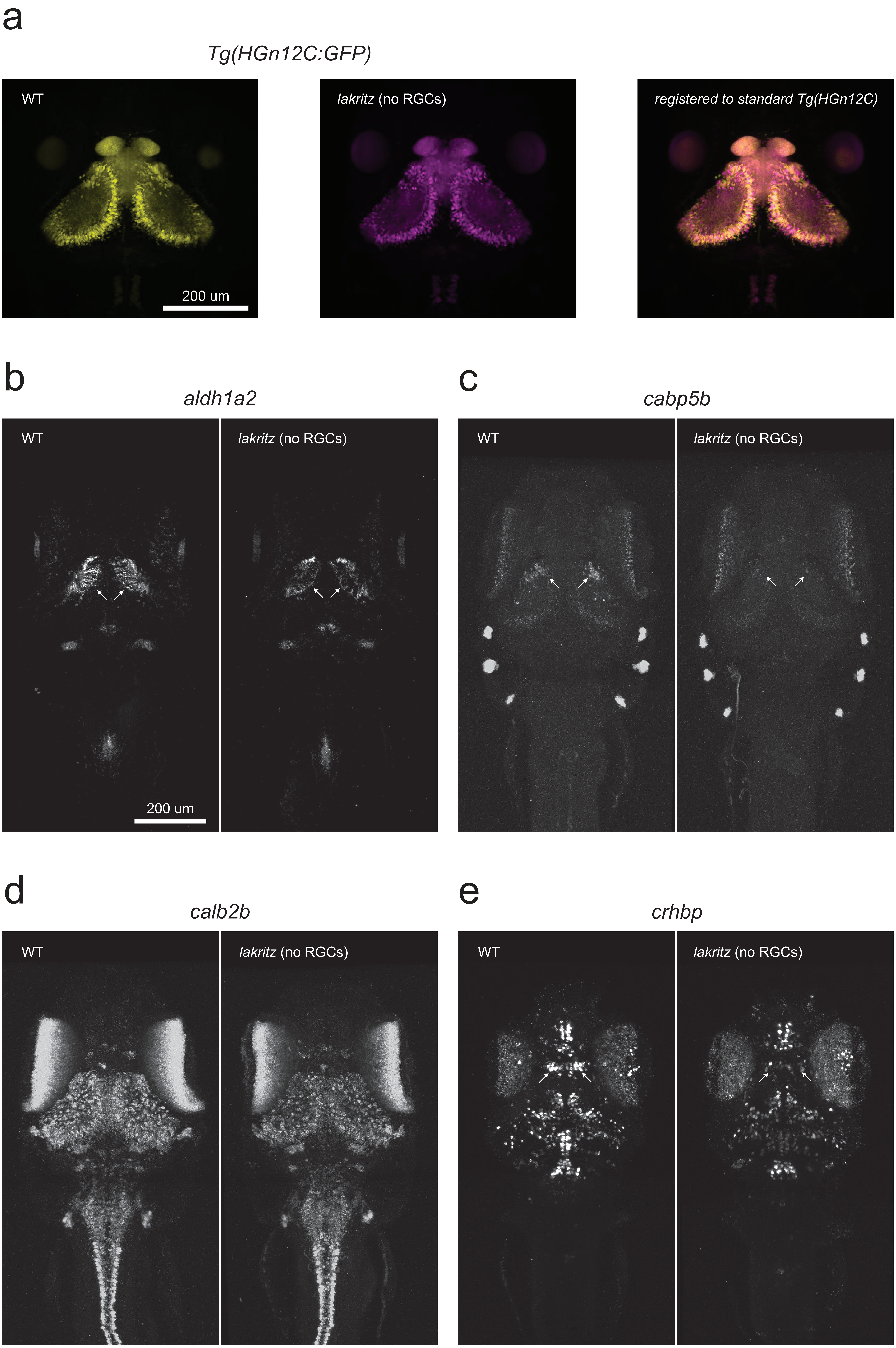

### Supplementary Figure 19

Supplementary Figure 19

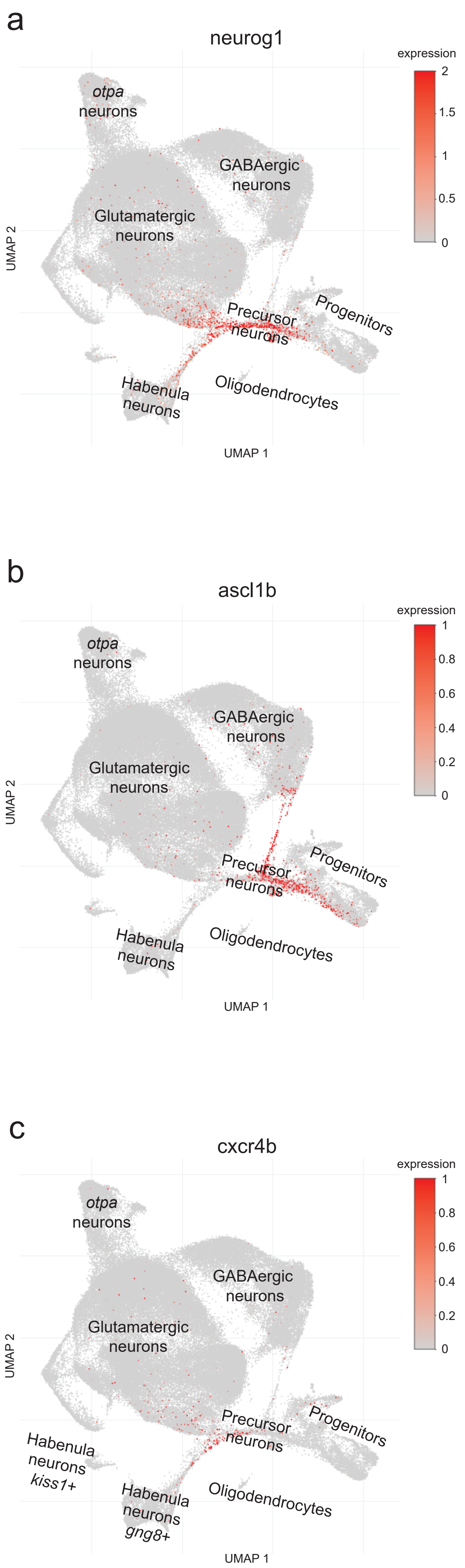

### Supplementary Figure 20

#### Supplementary Figure 20

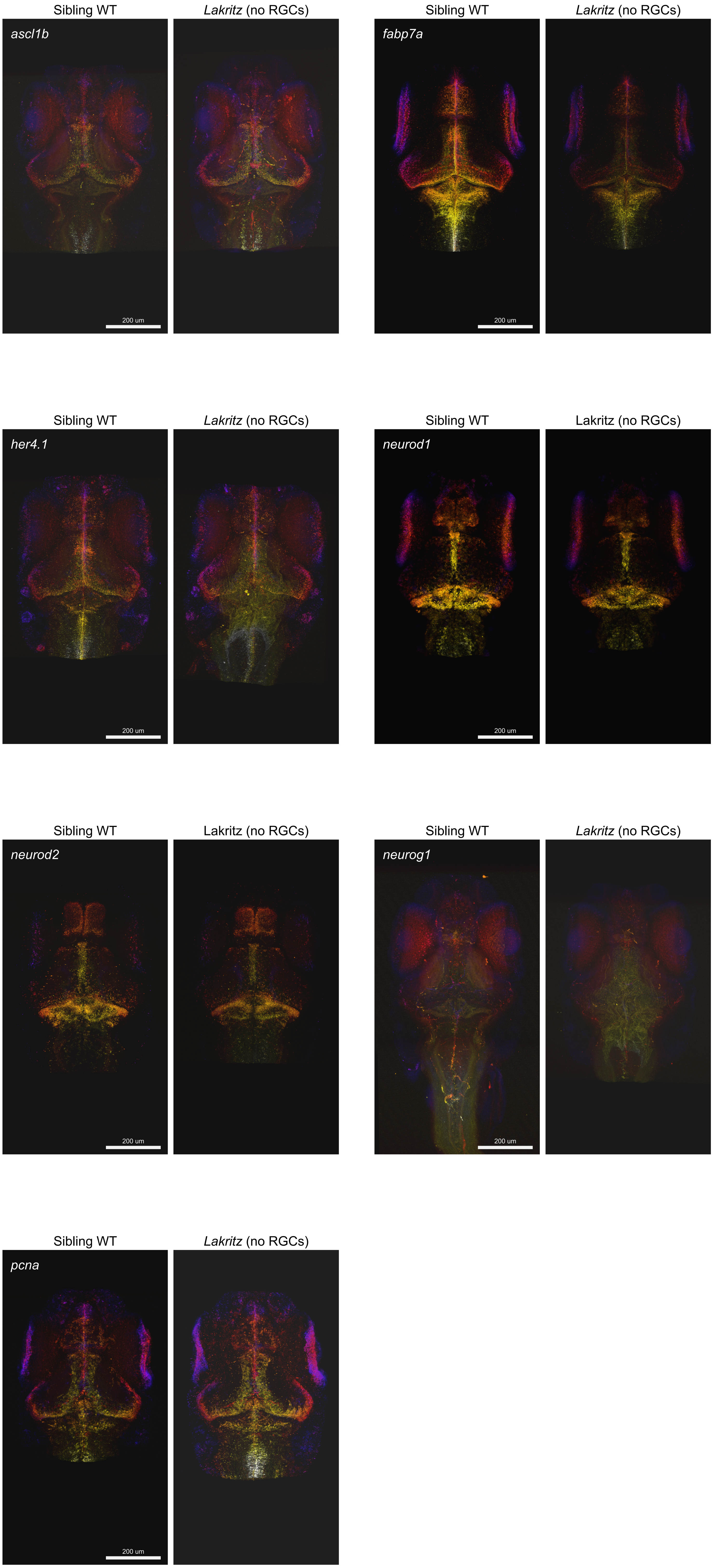

### Supplementary Figure 21

Supplementary Figure 21

a

*lakritz* (no RGCs), ChR2(-)

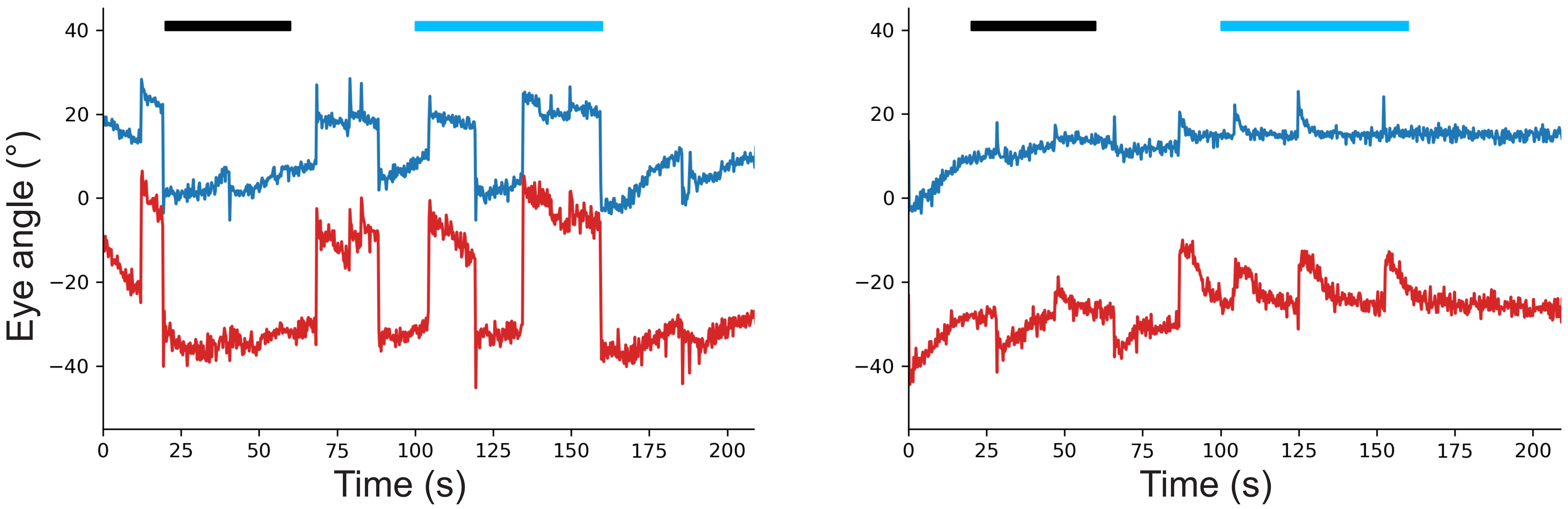


b

*lakritz* (no RGCs), ChR2(+)

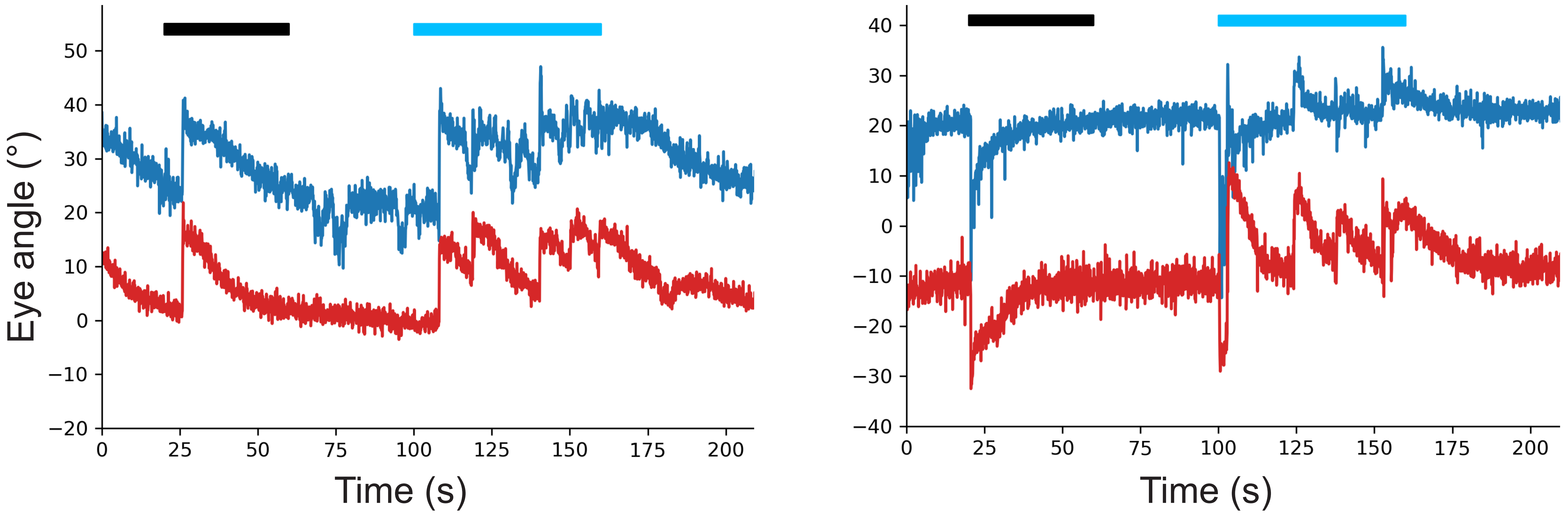


c

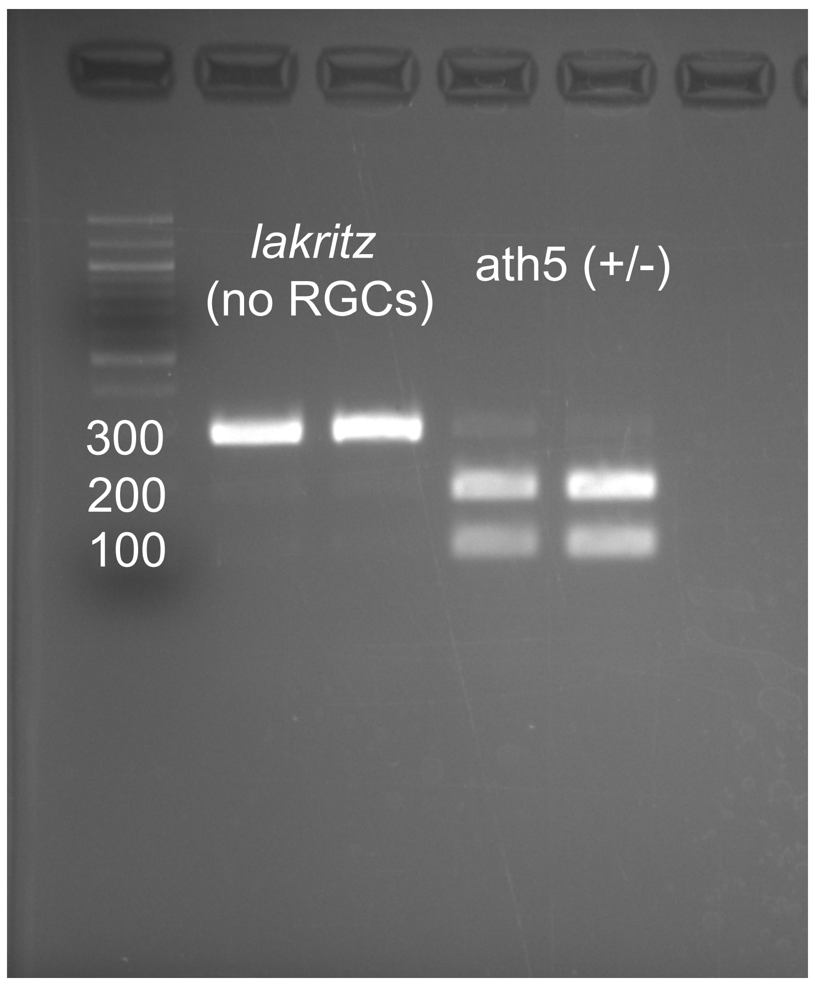

### Supplementary Figure 22

Supplementary Figure 22

a

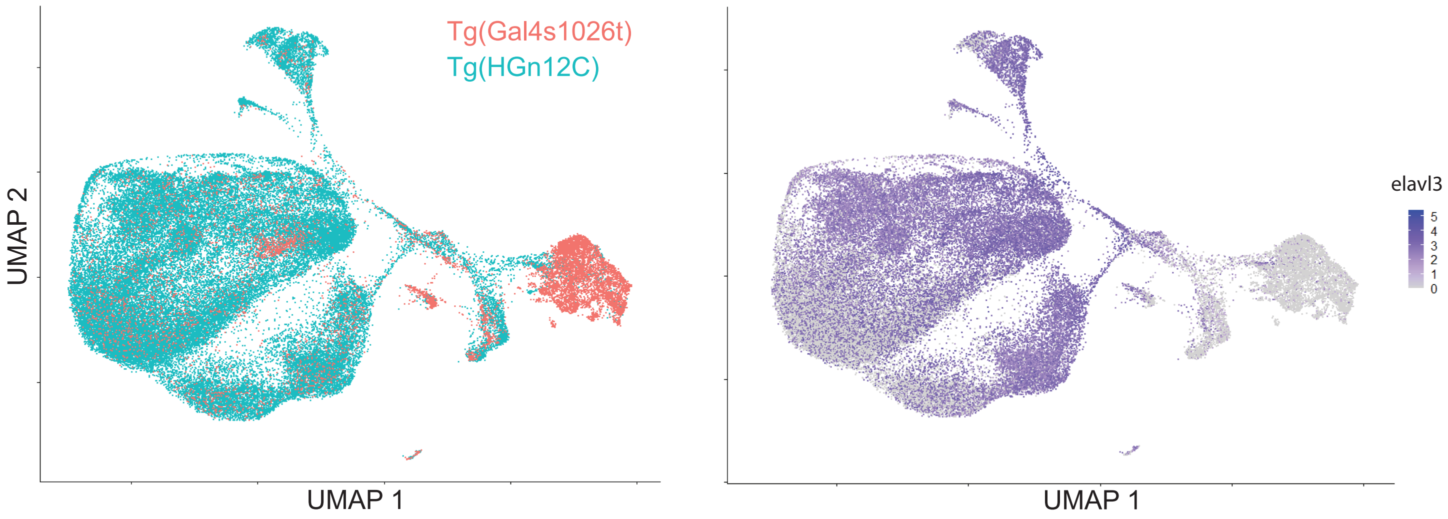


b

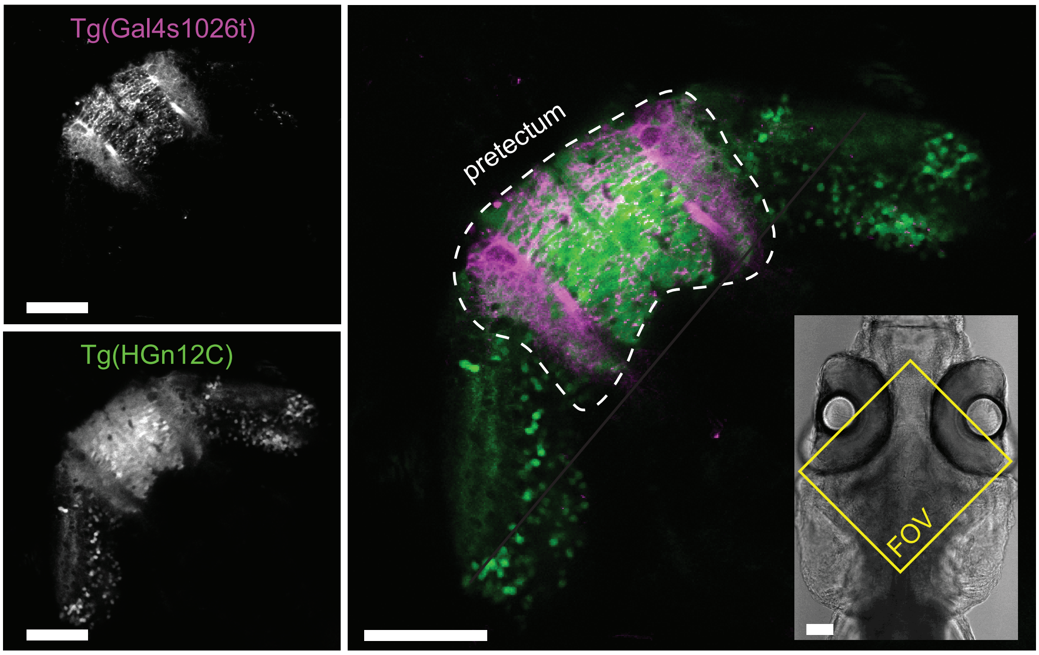


c

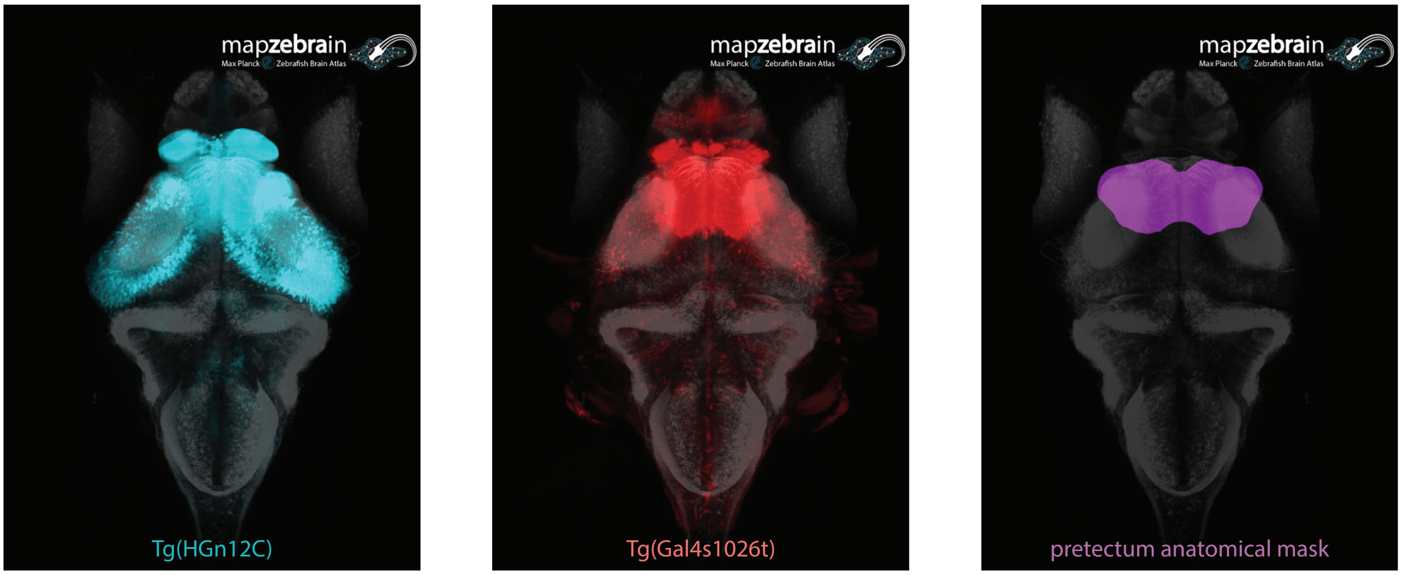
