## Supplementary Table 1 for "Retinal input influences pace of neurogenesis but not cell-type configuration of the visual forebrain"

| gene | top areas |
|---|---|
| aldh1a2 | pretectum__alar_part, area_postrema, inferior_dorsal_medulla_oblongata_stripe_1, retinal_arborization_field_9 |
| ascl1b | dorsal_thalamus_proper, periventricular_layer, medial_tegmentum_(remaining), pretectum__alar_part |
| atf5b | periventricular_layer, sac__spv, inferior_dorsal_medulla_oblongata_stripe_5 |
| bhlhe23 | dorsal_telencephalon_(pallium), dorsal_thalamus_proper, epiphysis, periventricular_layer, cerebellum |
| BX088 | cerebellum, nucleus_isthmi, olfactory_epithelium, superior_dorsal_medulla_oblongata_stripe_4 |
| cabp5b | stratum_fibrosum_et_griseum_superficiale, sfgs__sgc, pretectum__alar_part |
| calb1 | ventral_telencephalon_(subpallium), pretectum__alar_part, area_postrema, facial_motor_nucleus, vagus_motor_nucleus |
| calb2a | olfactory_epithelium, cerebellum, pretectum__alar_part, retinal_arborization_field_7, inferior_dorsal_medulla_oblongata_stripe_5 |
| calb2b | stratum_fibrosum_et_griseum_superficiale, inferior_dorsal_medulla_oblongata_stripe_2&3, stratum_opticum, posterior_lateral_line_ganglion, retinal_arborization_field_7 |
| cart2 | medial_tegmentum_(remaining), inferior_dorsal_medulla_oblongata_stripe_4, vagal_sensory_lobe, inferior_dorsal_medulla_oblongata_stripe_5 |
| cart3 | pretectum__alar_part, olfactory_bulb, epiphysis, nucleus_of_the_medial_longitudinal_fascicle_(pretectum,_basal_part) |
| ccka | inferior_dorsal_medulla_oblongata_stripe_5, vagus_motor_nucleus, nucleus_isthmi, intermediate_dorsal_medulla_oblongata_stripe_5 |
| cckb | ventral_habenula, periventricular_layer, dorsal_habenula, dorsal_telencephalon_(pallium), olfactory_bulb |
| chodl | area_postrema, vagus_motor_nucleus, olfactory_bulb, dorsal_telencephalon_(pallium), inferior_dorsal_medulla_oblongata_stripe_5 |
| cort | olfactory_bulb, dorsal_thalamus_proper, ventral_thalamus__alar_part, posterior_tuberculum_(basal_part_of_prethalamus_and_thalamus) |
| CR361 | periventricular_layer, cerebellum |
| CR848 | dorsal_thalamus_proper, pretectum__alar_part, olfactory_epithelium, inferior_dorsal_medulla_oblongata_stripe_5 |
| crhb | dorsal_telencephalon_(pallium), dorsal_thalamus_proper, eminentia_thalami_(remaining), pretectum__alar_part, ventral_thalamus__alar_part |
| crhbp | ventral_telencephalon_(subpallium), area_postrema, vagus_motor_nucleus, inferior_dorsal_medulla_oblongata_stripe_5 |
| dlx5a | dorsal_telencephalon_(pallium), ventral_telencephalon_(subpallium), eminentia_thalami_(remaining), olfactory_bulb |
| dopamine_transporter | pretectum__alar_part, posterior_tuberculum_(basal_part_of_prethalamus_and_thalamus), olfactory_bulb, dorsal_thalamus_proper, intermediate_hypothalamus_(remaining) |
| drgx | inferior_dorsal_medulla_oblongata_stripe_5, intermediate_dorsal_medulla_oblongata_stripe_5, dorsal_habenula, superior_dorsal_medulla_oblongata_stripe_4, intermediate_dorsal_medulla_oblongata_stripe_4 |
| elavl3 | dorsal_telencephalon_(pallium), periventricular_layer, dorsal_habenula, ventral_habenula, inferior_dorsal_medulla_oblongata_stripe_4, inferior_dorsal_medulla_oblongata_stripe_5 |
| emx2 | periventricular_layer, pretectum__alar_part, eminentia_thalami_(remaining), ventral_entopeduncular_nucleus, torus_longitudinalis, olfactory_epithelium |
| esrrb | cerebellum, periventricular_layer, pretectum__alar_part |
| foxb1a | retinal_arborization_field_7, pretectum__alar_part, periventricular_layer, intermediate_hypothalamus_(remaining) |
| gad1b | stratum_fibrosum_et_griseum_superficiale, inferior_dorsal_medulla_oblongata_stripe_4, inferior_dorsal_medulla_oblongata_stripe_5, stratum_opticum |
| gbx2 | dorsal_thalamus_proper, inferior_dorsal_medulla_oblongata_stripe_2&3, vagal_sensory_lobe, superior_dorsal_medulla_oblongata_stripe_5 |
| gjd2b | stratum_fibrosum_et_griseum_superficiale, sfgs__sgc, stratum_marginale, stratum_opticum |
| grm2b | dorsal_telencephalon_(pallium), olfactory_bulb, pretectum__alar_part, epiphysis, medial_octavolateralis_nucleus |
| gyg1b | cerebellum, epiphysis, torus_longitudinalis, dorsal_habenula, stratum_opticum |
| inhbaa | periventricular_layer, dorsal_thalamus_proper, cerebellum, stratum_marginale, ventral_telencephalon_(subpallium), pretectum__alar_part |
| insm2 | area_postrema, periventricular_layer, intermediate_dorsal_medulla_oblongata_stripe_5, olfactory_bulb, ventral_telencephalon_(subpallium), inferior_dorsal_medulla_oblongata_stripe_5 |
| mafba | intermediate_dorsal_medulla_oblongata_stripe_5, periventricular_layer, intermediate_dorsal_medulla_oblongata_stripe_4, medial_octavolateralis_nucleus, olfactory_bulb |
| mafbb | periventricular_layer, pretectum__alar_part, olfactory_bulb, olfactory_epithelium |
| mc5ra | pretectum__alar_part, dorsal_thalamus_proper, periventricular_layer, ventral_telencephalon_(subpallium) |
| mcm7 | periventricular_layer, dorsal_habenula, cerebellum, dorsal_telencephalon_(pallium), torus_longitudinalis |
| neurog1 | dorsal_thalamus_proper, dorsal_habenula |
| nfil3-6 | dorsal_telencephalon_(pallium), periventricular_layer, ventral_habenula, epiphysis |
| nfixb | periventricular_layer, dorsal_habenula, dorsal_telencephalon_(pallium), cerebellum |
| ngb | dorsal_habenula, dorsal_telencephalon_(pallium), torus_longitudinalis, olfactory_bulb |
| npy | periventricular_layer, dorsal_telencephalon_(pallium), pretectum__alar_part, dorsal_thalamus_proper, medial_tegmentum_(remaining), preoptic_region |
| nsun2 | periventricular_layer, epiphysis, torus_longitudinalis |
| ntn1b | inferior_dorsal_medulla_oblongata_stripe_1, medial_tegmentum_(remaining), olfactory_epithelium |
| onecut1 | ventral_habenula, dorsal_habenula, periventricular_layer, ventral_telencephalon_(subpallium), vagus_motor_nucleus |
| otpa | intermediate_dorsal_medulla_oblongata_stripe_2&3, superior_dorsal_medulla_oblongata_stripe_2&3, preoptic_region, intermediate_dorsal_medulla_oblongata_stripe_1 |
| pax3a | periventricular_layer, cerebellum, pretectum__alar_part, inferior_dorsal_medulla_oblongata_stripe_4, superior_dorsal_medulla_oblongata_stripe_2&3 |
| pax6a | olfactory_bulb, ventral_thalamus__alar_part, ventral_telencephalon_(subpallium) |
| pax7a | pretectum__alar_part, periventricular_layer, dorsal_thalamus_proper, cerebellum |
| pax7b | periventricular_layer, dorsal_habenula, stratum_opticum, pretectum__alar_part, torus_longitudinalis |
| pcp4l1 | cerebellum, epiphysis, olfactory_bulb, superior_dorsal_medulla_oblongata_stripe_4, dorsal_telencephalon_(pallium) |
| penkb | periventricular_layer, dorsal_telencephalon_(pallium), dorsal_thalamus_proper, vagus_motor_nucleus, posterior_tuberculum_(basal_part_of_prethalamus_and_thalamus), ventral_telencephalon_(subpallium) |
| pnocb | nucleus_of_the_medial_longitudinal_fascicle_(pretectum,_basal_part), superior_dorsal_medulla_oblongata_stripe_1_(remaining), trochlear_motor_nucleus, medial_tegmentum_(remaining), olfactory_epithelium, inferior_dorsal_medulla_oblong |
| pou4f2 | periventricular_layer, inferior_dorsal_medulla_oblongata_stripe_5 |
| pth2 | dorsal_thalamus_proper, periventricular_layer, inferior_dorsal_medulla_oblongata_stripe_5 |
| sema3fb | nucleus_isthmi, periventricular_layer, torus_semicircularis, epiphysis |
| six3b | ventral_thalamus__alar_part, dorsal_telencephalon_(pallium) |
| sox14 | periventricular_layer, torus_longitudinalis, cerebellum, torus_semicircularis |
| sox1b | dorsal_telencephalon_(pallium), ventral_telencephalon_(subpallium), olfactory_bulb, periventricular_layer, ventral_habenula |
| sox7 | intermediate_dorsal_medulla_oblongata_stripe_4, olfactory_epithelium, intermediate_dorsal_medulla_oblongata_stripe_5, medial_octavolateralis_nucleus |
| sp9 | dorsal_telencephalon_(pallium), ventral_telencephalon_(subpallium), olfactory_bulb, eminentia_thalami_(remaining) |
| sst1.2 | dorsal_thalamus_proper, inferior_dorsal_medulla_oblongata_stripe_5, inferior_dorsal_medulla_oblongata_stripe_4, vagus_motor_nucleus |
| tac1 | dorsal_telencephalon_(pallium), olfactory_bulb, superior_dorsal_medulla_oblongata_stripe_2&3, vagus_motor_nucleus |
| tac3b | olfactory_bulb, dorsal_telencephalon_(pallium), dorsal_thalamus_proper, rostral_hypothalamus |
| tfap2a | periventricular_layer, inferior_dorsal_medulla_oblongata_stripe_4, medial_octavolateralis_nucleus, inferior_dorsal_medulla_oblongata_stripe_5 |
| tfap2b | periventricular_layer, pretectum__alar_part, medial_octavolateralis_nucleus, torus_longitudinalis, inferior_dorsal_medulla_oblongata_stripe_4, inferior_dorsal_medulla_oblongata_stripe_5 |
| tfap2d | periventricular_layer, dorsal_thalamus_proper, torus_semicircularis, nucleus_isthmi, pretectum__alar_part |
| tfap2e | periventricular_layer, torus_longitudinalis, cerebellum, olfactory_bulb, pretectum__alar_part |
| th | area_postrema, pretectum__alar_part, vagus_motor_nucleus, vagal_sensory_lobe |
| tph2 | superior_raphe, epiphysis, dorsal_habenula, dorsal_thalamus_proper |
| txn | olfactory_epithelium, facial_motor_nucleus, superior_raphe |
| uts1 | trochlear_motor_nucleus, nucleus_of_the_medial_longitudinal_fascicle_(pretectum,_basal_part), medial_tegmentum_(remaining), locus_coeruleus |
| vglut2a | dorsal_habenula, ventral_habenula, epiphysis, inferior_dorsal_medulla_oblongata_stripe_5 |
| zic1 | cerebellum, eminentia_thalami_(remaining), dorsal_habenula, torus_longitudinalis, ventral_habenula |
| zic2a | dorsal_habenula, cerebellum, torus_longitudinalis, dorsal_thalamus_proper, ventral_habenula |
| zic4 | dorsal_habenula, cerebellum, torus_longitudinalis, eminentia_thalami_(remaining), ventral_telencephalon_(subpallium), ventral_habenula |
