## Supplementary Figure 6 for "Retinal input influences pace of neurogenesis but not cell-type configuration of the visual forebrain"

### Unrelated WT glutamatergic clusters

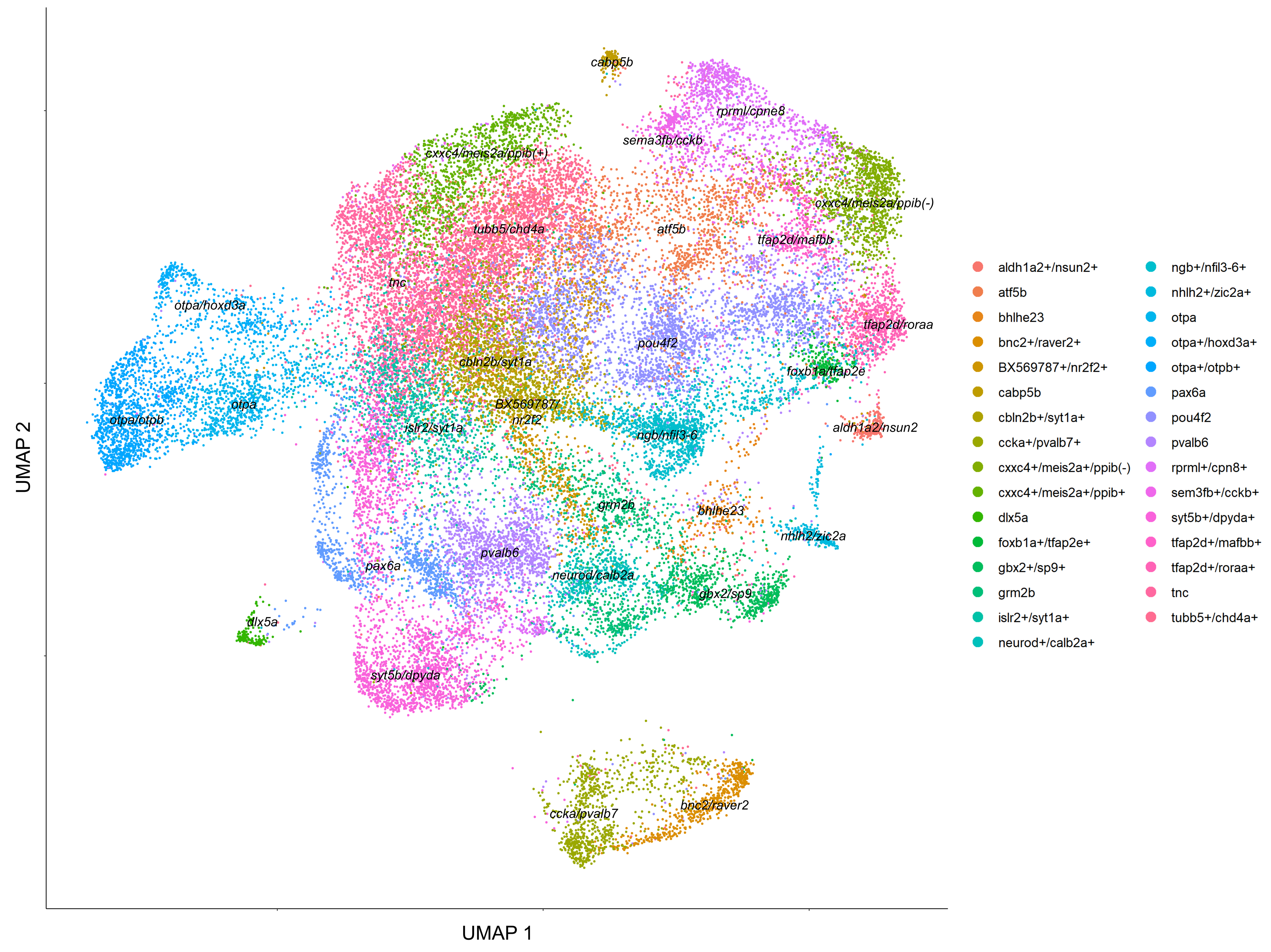


### Sibling WT glutamatergic clusters

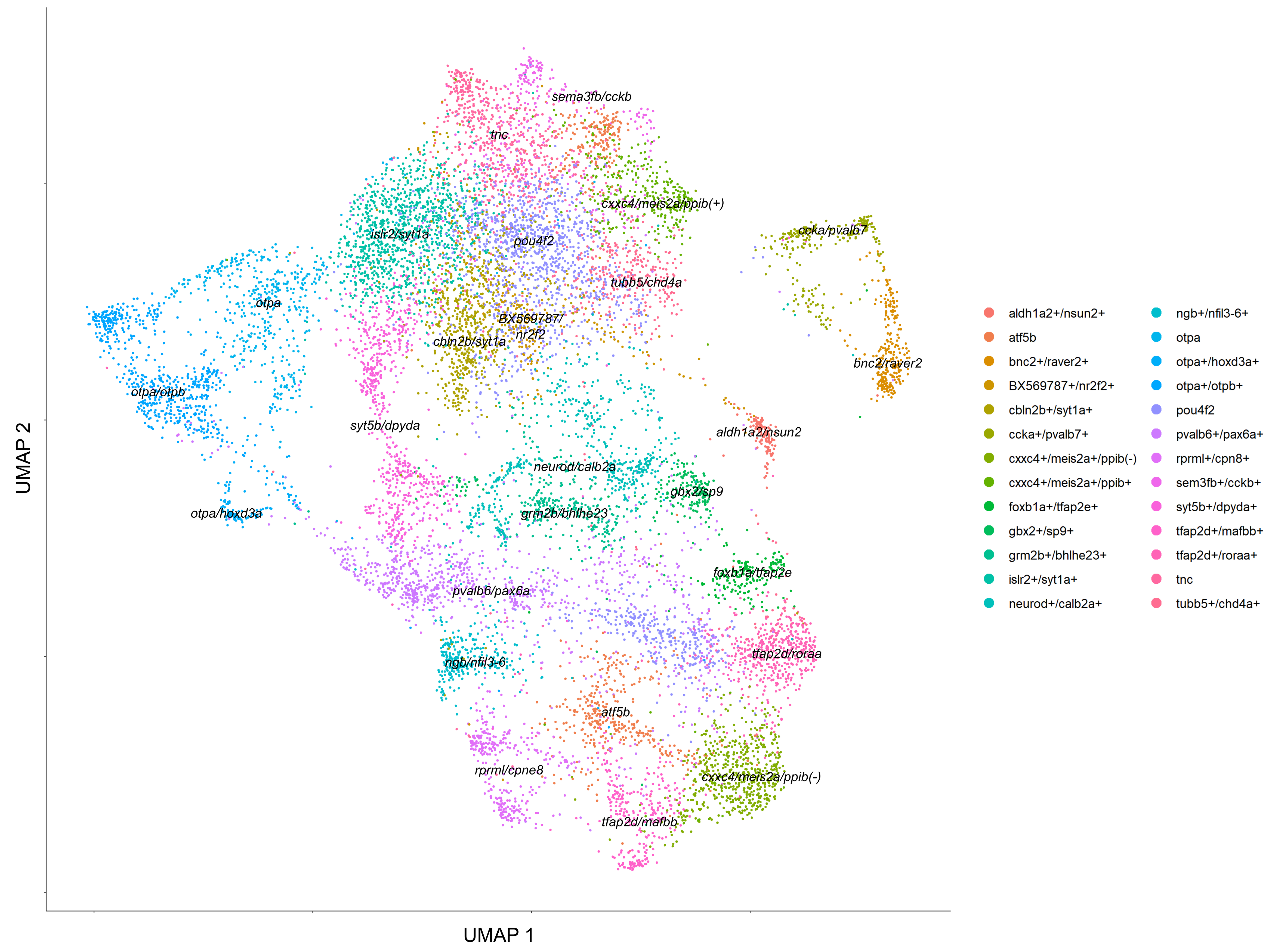


### *Lakritz* (no RGCs) glutamatergic clusters

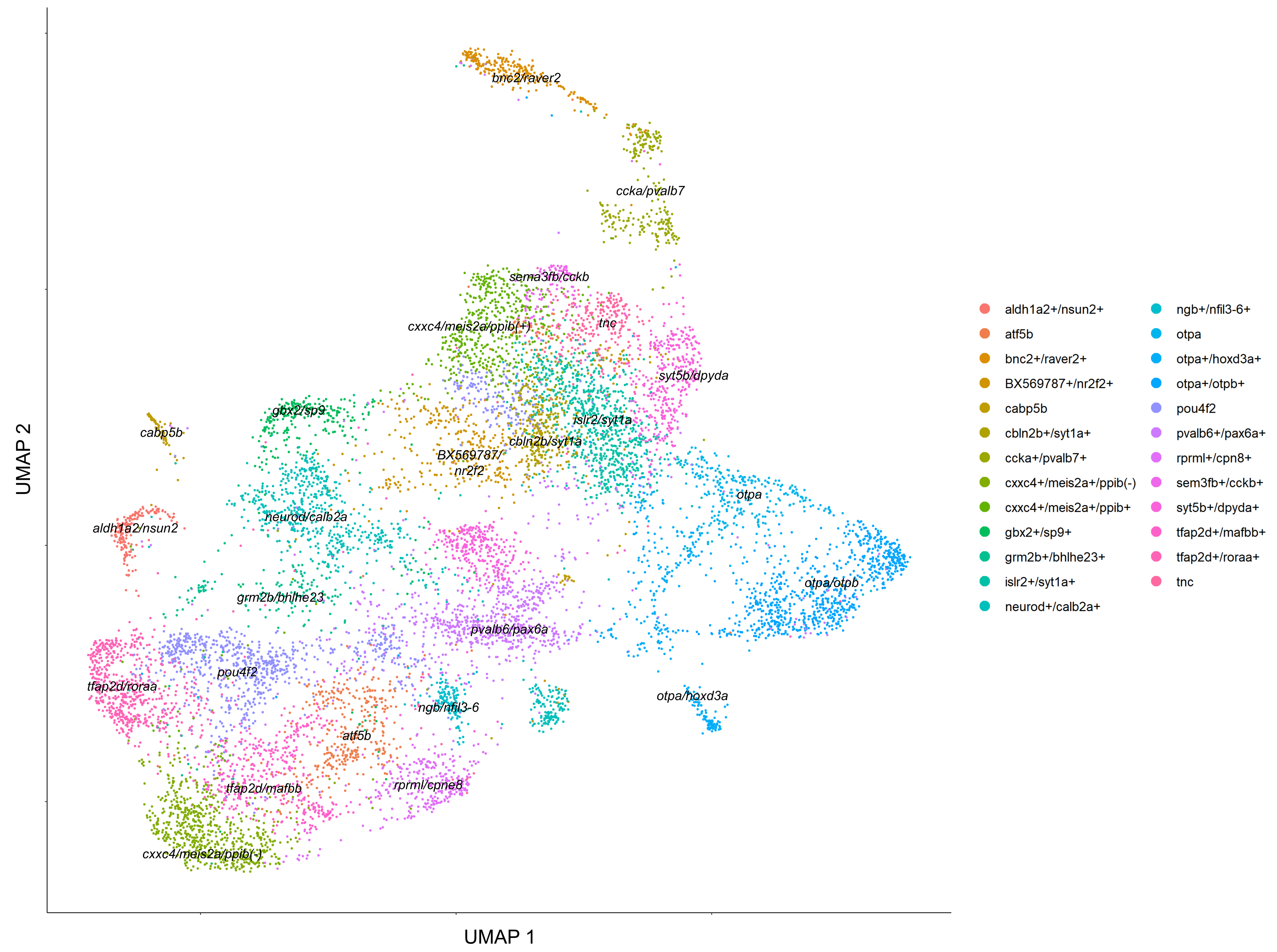
