## Supplementary Figure 8 for "Retinal input influences pace of neurogenesis but not cell-type configuration of the visual forebrain"

### Unrelated WT GABAergic clusters

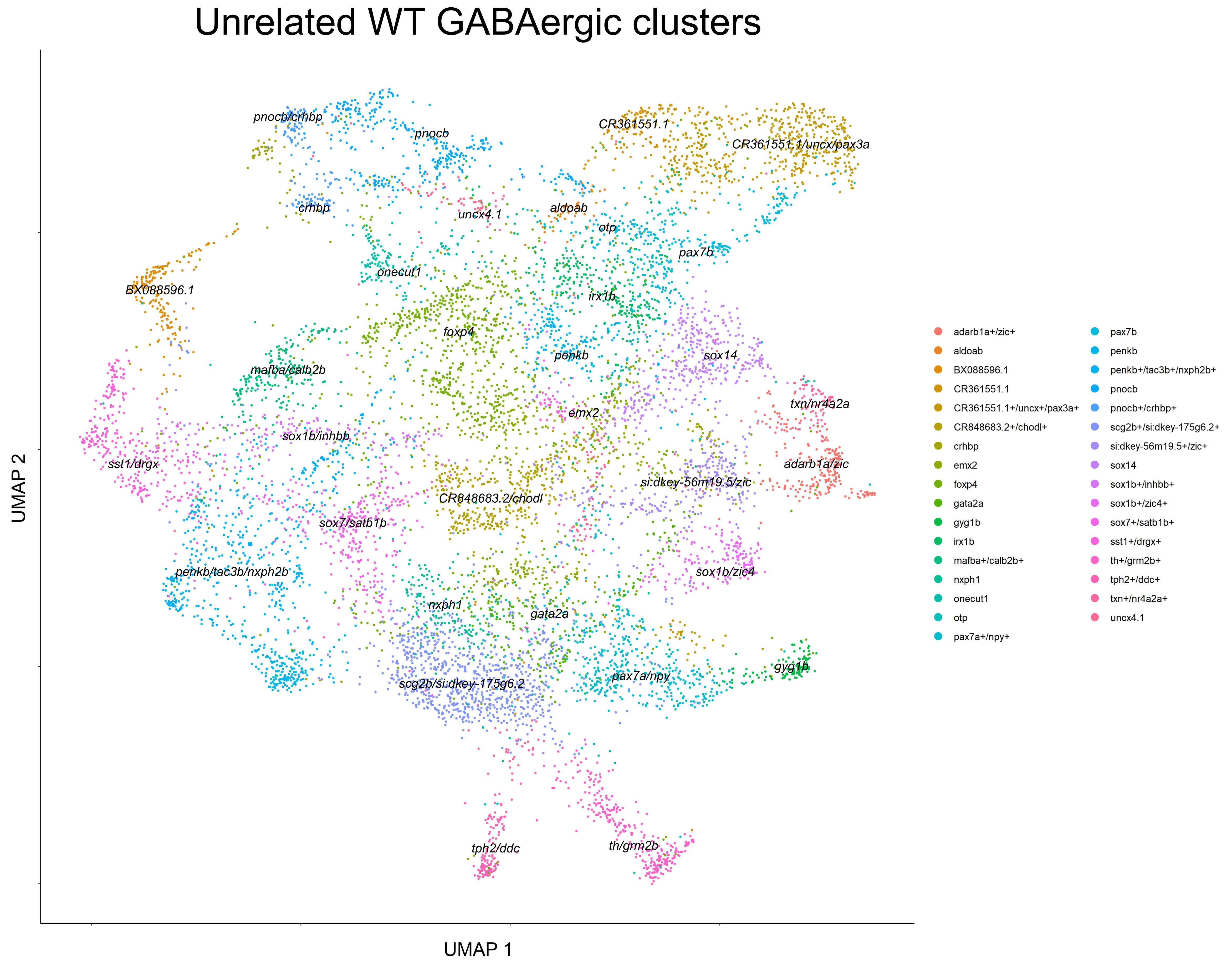


### Sibling WT GABAergic clusters

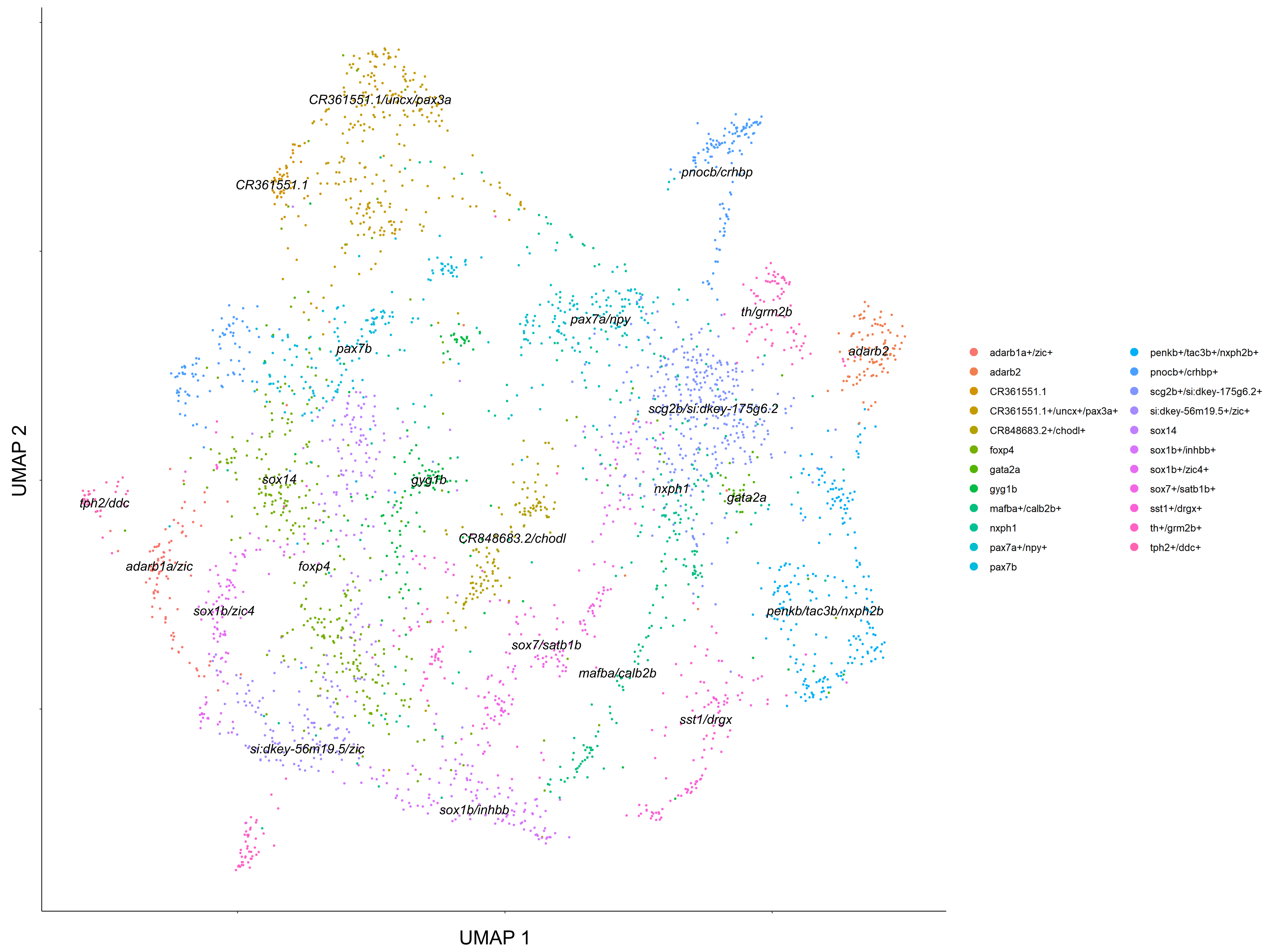


### *Lakritz* (no RGCs) GABAergic clusters

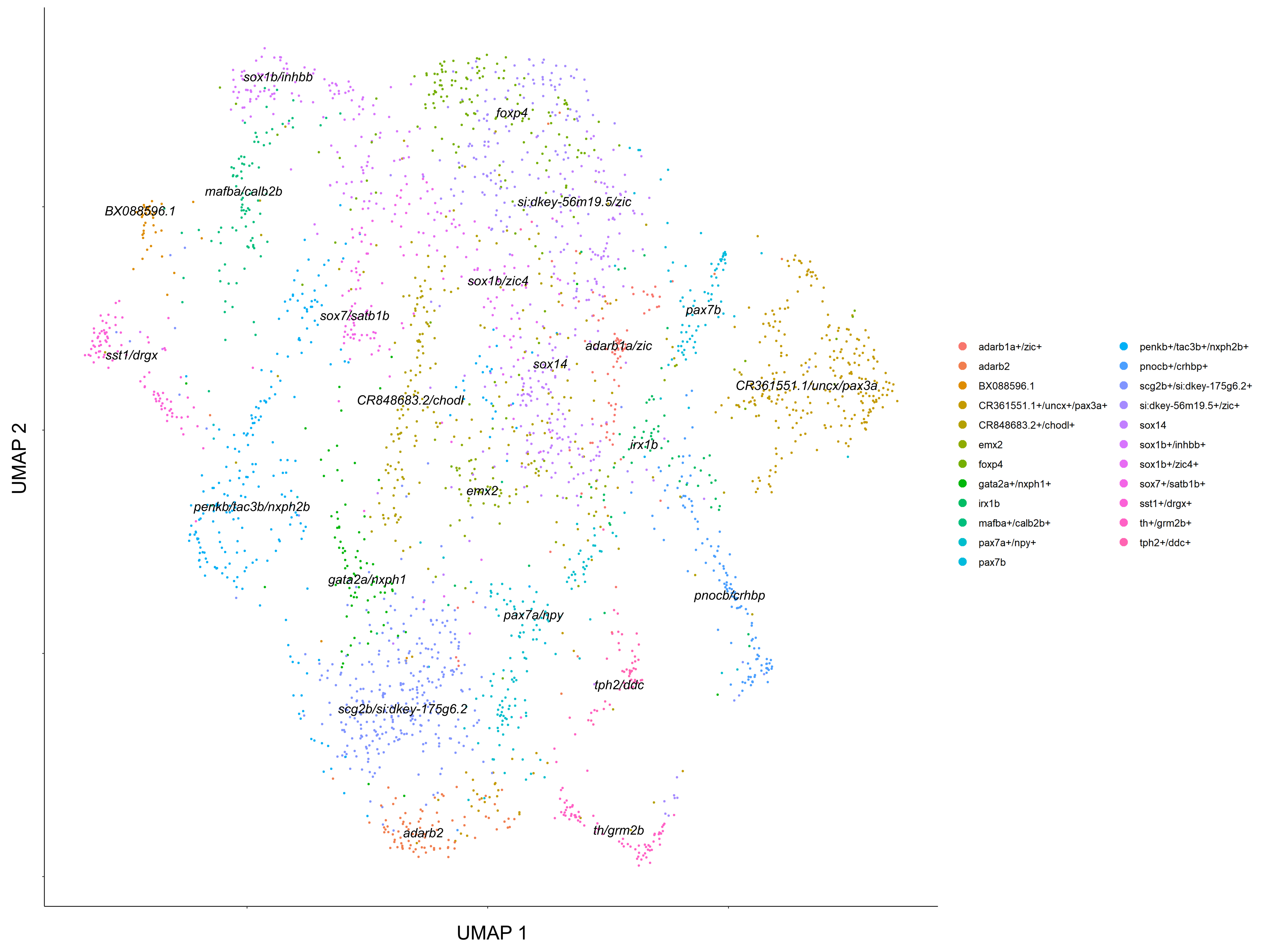
