## Supplementary Table 2 for "Retinal input influences pace of neurogenesis but not cell-type configuration of the visual forebrain"

| | trajectory to: | | | | | | | labels in clusters: | | | | |
|---|---|---|---|---|---|---|---|---|---|---|---|---|
| gene | progenitors | gabaergic clusters | glutamatergic clusters | general habenula | gng8 | kiss1 | | progenitors | gabaergic clusters | glutamatergic clusters | gng8 | kiss1 |
| dla | yes | yes | yes | yes | no | no | | no | no | no | no | no |
| hmgb2a | yes | yes | yes | yes | yes | yes | | yes | no | yes | no | no |
| abhd6a | no | yes | yes | no | no | no | | no | no | no | no | no |
| neurog1 | no | no | yes | yes | yes | no | | no | no | no | no | no |
| hmga1a | no | yes | yes | yes | yes | yes | | no | yes | yes | yes | yes |
| sox3 | no | yes | ambiguous | no | no | no | | no | no | no | no | no |
| notch1a | no | yes | yes | no | no | no | | no | no | no | no | no |
| si:ch73-21g5.7 | no | no | no | no | no | no | | no | no | no | no | no |
| hmgb2b | yes | yes | yes | yes | yes | yes | | yes | yes | yes | yes | yes |
| stmn1a | no | yes | yes | yes | no | no | | no | ambiguous | ambiguous | no | no |
| dlb | no | yes | yes | yes | no | no | | no | no | no | no | no |
| notch3 | yes | no | no | no | no | no | | yes | no | no | no | no |
| id1 | yes | no | no | no | no | no | | yes | no | no | no | no |
| sox11b | no | yes | yes | yes | no | no | | no | yes | yes | no | no |
| si:dkey-151g10.6 | yes | yes | yes | yes | yes | yes | | yes | yes | yes | yes | yes |
| ccnd1 | yes | no | no | no | no | no | | yes | no | no | no | no |
| msi1 | yes | ambiguous | no | yes | yes | no | | yes | no | no | ambiguous | no |
| CU467822.1 | yes | yes | yes | no | no | no | | yes | yes | yes | no | no |
| ascl1b.1 | no | yes | no | no | no | no | | no | no | no | no | no |
| tpt1 | yes | yes | yes | yes | yes | yes | | yes | yes | yes | yes | yes |
| si:ch211-222l21.1 | yes | yes | yes | yes | yes | yes | | yes | yes | yes | yes | yes |
| insm1b | no | yes | yes | yes | no | no | | no | ambiguous | ambiguous | no | no |
| rtca | no | yes | yes | no | no | no | | no | no | no | no | no |
| lfng | yes | ambiguous | no | no | no | no | | yes | no | no | no | no |
| ran | yes | yes | yes | yes | yes | yes | | yes | yes | yes | yes | yes |
| BX465834.1 | yes | yes | yes | no | no | no | | yes | yes | yes | no | no |
| ebf2 | no | no | yes | ambiguous | no | no | | no | no | no | no | no |
| ascl1a | no | yes | no | no | no | no | | no | no | no | no | no |
| rack1 | yes | yes | yes | yes | yes | yes | | yes | yes | yes | yes | yes |
| chd7 | no | yes | yes | yes | yes | yes | | no | no | no | yes | yes |
| cdkn1ca | no | yes | yes | yes | no | yes | | no | no | no | no | yes |
| zbtb18 | no | no | yes | yes | yes | no | | no | no | yes | no | no |
| ddx21 | yes | no | no | no | no | no | | yes | no | no | no | no |
| sox19a | yes | no | no | no | no | no | | yes | no | no | no | no |
| hes6 | no | yes | yes | yes | no | no | | no | ambiguous | ambiguous | no | no |
| naca | yes | yes | yes | yes | yes | yes | | yes | yes | yes | yes | yes |
| si:dkey-85k7.7 | yes | no | no | no | no | no | | yes | no | no | no | no |
| si:dkey-42i9.4 | yes | yes | yes | yes | yes | yes | | yes | yes | yes | no | no |
| inavaa | no | ambiguous | ambiguous | no | no | no | | no | no | no | no | no |
| snrpf | no | yes | yes | yes | yes | yes | | no | yes | yes | no | no |
| adh5 | yes | yes | yes | yes | yes | yes | | ambiguous | ambiguous | yes | no | no |
| si:ch211-212k18.5 | yes | no | no | no | no | no | | yes | no | no | no | no |
| btf3 | yes | yes | yes | yes | yes | yes | | yes | yes | yes | yes | yes |
| nfia | ambiguous | yes | yes | no | no | no | | no | no | yes | no | no |
| serbp1a | yes | yes | yes | yes | yes | yes | | yes | yes | yes | yes | yes |
| sb:cb81 | yes | no | no | no | no | no | | yes | no | no | no | no |
| fabp7a | yes | no | no | no | no | no | | yes | no | no | no | no |
| msna | yes | no | no | ambiguous | no | no | | yes | no | no | no | no |
| tspan7 | no | no | no | no | no | no | | no | no | no | no | no |
| tcf12 | yes | ambiguous | ambiguous | yes | ambiguous | no | | ambiguous | no | no | no | no |
| cdk6 | no | no | ambiguous | no | no | no | | no | no | no | no | no |
| im:7152348 | yes | no | no | yes | yes | no | | yes | no | no | yes | no |
| fgfbp3 | yes | yes | no | no | no | no | | yes | no | no | no | no |
| nop58 | ambiguous | no | no | no | no | no | | no | no | no | no | no |
| CR751602.2 | yes | no | no | no | no | no | | yes | no | no | no | no |
| atp6v0e1 | yes | yes | yes | yes | yes | yes | | yes | yes | yes | no | no |
| sox2 | yes | ambiguous | no | no | no | no | | yes | no | no | no | no |
| txnipa | yes | no | no | no | no | no | | yes | no | no | no | no |
| ptprz1a | yes | no | no | no | no | no | | yes | no | no | no | no |
| otx2a | ambiguous | ambiguous | no | no | no | no | | no | no | no | no | no |
| zeb2a | yes | ambiguous | no | no | no | no | | no | no | no | no | no |
| snu13b | yes | no | no | no | no | no | | no | no | no | no | no |
| selenoh | yes | no | no | no | no | no | | yes | no | no | no | no |
| sinhcafl.1 | yes | yes | yes | yes | no | no | | no | yes | yes | no | no |
| nrarpa | yes | no | no | no | no | no | | yes | no | no | no | no |
| mdka | yes | no | no | no | no | no | | yes | no | no | no | no |
| pax6a | yes | no | no | no | no | no | | yes | no | no | no | no |
| ak2 | yes | no | no | no | no | no | | yes | no | no | no | no |
| ahcy | ambiguous | no | no | no | no | no | | no | no | no | no | no |
| tfdp2 | no | no | no | no | no | no | | no | no | no | no | no |
| cldn5a | yes | no | no | no | no | no | | yes | no | no | no | no |
| smarcb1b | no | yes | yes | ambiguous | no | no | | no | yes | yes | no | no |
| lima1a | no | ambiguous | no | yes | no | no | | no | no | no | no | no |
| si:dkey-239h2.3 | yes | no | no | no | no | no | | yes | no | no | no | no |
| pno1 | yes | no | no | no | no | no | | yes | no | no | no | no |
| si:dkey-56m19.5 | yes | yes | yes | yes | yes | no | | yes | yes | yes | yes | no |
| nr2f2 | yes | yes | yes | yes | yes | no | | yes | yes | yes | yes | no |
| zgc:110796 | no | no | no | no | no | no | | no | no | no | no | no |
| tgif1 | no | yes | yes | yes | ambiguous | no | | no | yes | yes | no | no |
| npm1a | yes | no | no | no | no | no | | yes | no | no | no | no |
| gng5 | ambiguous | no | no | yes | no | no | | ambiguous | no | ambiguous | no | no |
| nop56 | yes | ambiguous | ambiguous | ambiguous | no | no | | yes | ambiguous | ambiguous | no | no |
| fgfr3 | yes | no | no | no | no | no | | yes | no | no | no | no |
| gtpbp4 | yes | yes | yes | yes | no | no | | yes | yes | yes | no | no |
| ybx1 | yes | yes | yes | yes | yes | yes | | yes | yes | yes | yes | yes |
| prdx2 | yes | yes | yes | yes | yes | yes | | yes | yes | yes | yes | ambiguous |
| CR318588.4 | yes | yes | yes | yes | yes | no | | yes | yes | yes | yes | no |
| her6 | yes | no | no | no | no | no | | yes | no | no | no | no |
| cebpd | yes | no | no | no | no | no | | yes | no | no | no | no |
| hmgn6 | yes | yes | yes | yes | yes | yes | | yes | yes | yes | yes | yes |
| cct2.1 | yes | yes | yes | yes | yes | yes | | yes | yes | yes | yes | no |
| nme2b.1 | yes | yes | yes | yes | yes | yes | | yes | yes | yes | yes | yes |
| insm1a | no | yes | yes | yes | no | no | | no | no | no | no | no |
| ppiaa | yes | yes | yes | yes | yes | yes | | yes | yes | yes | yes | yes |
| shox2 | no | no | yes | no | no | no | | no | no | ambiguous | no | no |
| h2afva | yes | yes | yes | yes | yes | yes | | ambiguous | yes | yes | yes | yes |
| khdrbs1a | yes | yes | yes | yes | yes | yes | | yes | yes | yes | yes | yes |
| rasgef1ba | yes | ambiguous | yes | no | no | no | | no | yes | yes | no | no |
| pou3f2b | no | yes | yes | yes | no | no | | no | no | yes | no | no |
| ddx39ab | yes | yes | yes | yes | no | no | | no | yes | yes | no | no |
| h2afvb | yes | yes | yes | yes | yes | yes | | yes | yes | yes | yes | yes |
| snrpe | yes | yes | yes | yes | yes | yes | | no | yes | yes | ambiguous | no |
| nop10 | yes | no | no | no | no | no | | yes | no | no | no | no |
| anp32b | ambiguous | no | no | no | no | no | | no | no | no | no | no |
| lmnb2 | no | no | ambiguous | no | no | no | | no | no | no | no | no |
| smc1al | no | yes | yes | yes | no | no | | no | yes | yes | no | no |
| cct3 | yes | yes | yes | yes | yes | yes | | no | yes | yes | no | no |
| si:dkey-67c22.2 | yes | yes | yes | yes | yes | ambiguous | | yes | yes | yes | yes | no |
| rad21a | yes | yes | yes | yes | yes | yes | | no | yes | yes | yes | no |
| CR848812.1 | ambiguous | no | no | no | no | no | | no | no | no | no | no |
| TXN | yes | yes | yes | yes | yes | yes | | yes | yes | yes | yes | no |

| | | | | | | | | | | | | |
|---|---|---|---|---|---|---|---|---|---|---|---|---|
| hmgn2 | yes | yes | yes | yes | yes | yes | | yes | yes | yes | yes | yes |
| ranbp1 | ambiguous | no | no | no | no | no | | no | no | no | no | no |
| syncrip | yes | yes | yes | yes | yes | yes | | ambiguous | yes | yes | yes | yes |
| si:ch211-288g17.3 | yes | yes | yes | yes | yes | yes | | no | yes | yes | yes | yes |
| seta | yes | yes | yes | yes | yes | yes | | ambiguous | yes | yes | yes | yes |
| si:ch73-281n10.2 | yes | yes | yes | yes | yes | yes | | yes | yes | yes | yes | yes |
| tuba8l4 | yes | yes | yes | yes | yes | yes | | yes | yes | yes | yes | yes |
